# The explosive radiation of the Neotropical *Tillandsia* subgenus *Tillandsia* (Bromeliaceae) has been accompanied by pervasive hybridization

**DOI:** 10.1101/2023.11.16.567341

**Authors:** Gil Yardeni, Michael H. J. Barfuss, Walter Till, Matthew R. Thornton, Clara Groot Crego, Christian Lexer, Thibault Leroy, Ovidiu Paun

**Affiliations:** Department of Botany and Biodiversity Research, University of Vienna, Vienna, Austria; Institute of Computational Biology, Department of Biotechnology, University of Life Sciences and Natural Resources, Vienna, Austria; Vienna Graduate School of Population Genetics, Vienna, Austria; GenPhySE, Université de Toulouse, INRAE, ENVT, 31326, Castanet Tolosan, France

**Keywords:** *Tillandsia*, gene tree discordance, hybridization, phylogenomics, bromeliad, Neotropical diversity, species network

## Abstract

The recent rapid radiation of *Tillandsia* subgenus *Tillandsia* (Bromeliaceae) provides an attractive system to study the drivers and constraints of species diversification. This species-rich Neotropical monocot clade includes predominantly epiphytic species displaying vast phenotypic diversity. Recent in-depth phylogenomic work revealed that the subgenus originated within the last 7 MY, with one major expansion from South into Central America within the last 5 MY. However, disagreements between phylogenies and lack of resolution at shallow nodes suggest that hybridization may have occurred throughout the radiation, together with frequent incomplete lineage sorting and rapid gene family evolution. We used whole-genome resequencing data to explore the evolutionary history of representative ingroup species employing both tree-based and network approaches. Our results indicate that lineage co-occurrence does not predict relatedness and confirm significant deviations from a tree-like structure, coupled with pervasive gene tree discordance. Focusing on hybridization, ABBA-BABA and related statistics were used to infer the rates and relative timing of events, while topology weighting uncovered high heterogeneity of the phylogenetic signal along the genome. High rates of hybridization within and among subclades suggest that, contrary to previous hypotheses, the expansion of subgenus *Tillandsia* into Central America proceeded through several dispersal events, punctuated by episodes of diversification and gene flow. Network analysis revealed reticulation as a plausible propeller during radiation and establishment across different ecological niches. This work contributes a plant example of prevalent hybridization during rapid species diversification, supporting the hypothesis that interspecific gene flow facilitates explosive diversification.

---

Rapid evolutionary radiations are characterised by accelerated and substantial diversification, usually following dispersal into a novel geographic area. As a myriad of diverse ecological spaces become available, a lineage can opportunistically adapt to quickly occupy them in a process termed adaptive radiation (Hughes et al. 2015; Linder 2008; Naciri and Linder 2020; Rundell and Price 2009; Stroud and Losos 2016). Rapid radiations provide ample opportunities to investigate the mechanisms driving diversification. While not all rapid radiations are adaptive, i.e., not necessarily associated with an increase in ecological occupancy (Givnish 2015; Schluter 2000; Stroud and Losos 2016), evolutionary radiations are generally connected with habitat heterogeneity (Paun et al. 2016; Soltis et al. 2019), climatic fluctuations (Slovak et al. 2023), landscape fragmentation (Hughes and Atchison 2015) and/or orogenesis (Boschman and Condamine 2022). Ubiquitous among angiosperms in general and in the Neotropics in particular, plant radiations remain to date understudied and elusive, especially when compared to animal systems (but see e.g., Drummond et al. 2012; Lagomarsino et al. 2016; Linder 2008; Paun et al. 2016; Pérez-Escobar et al. 2017; Richardson et al. 2001).

Phylogenomic studies of young, rapid radiations face multiple challenges. Low sequence divergence and incomplete reproductive barriers increase the likelihood of short internal branches and impede phylogenomic resolution (Giarla and Esselstyn 2015; Straub et al. 2014; Whitfield and Lockhart 2007). Furthermore, population-level processes during rapid radiations - such as fluctuations in population size and incomplete lineage sorting (ILS) - generate high rates of gene-tree conflict, potentially reducing the phylogenetic signal (Oliver 2013; Parins-Fukuchi et al. 2021). Such processes were traditionally deemed as analytical noise when aiming to reconstruct bifurcating species trees. Yet an increasing amount of research indicates that episodes of phylogenomic conflict represent typical signatures of microevolutionary processes and are often associated with rapid phenotypic shifts (Filiault et al. 2018; Parins-Fukuchi et al. 2021). Hence, non-bifurcating relationships should be regarded and studied as fundamental biological phenomena of diversification, underlying the formation of species (Malinsky et al. 2018; Slovak et al. 2023; Wogan et al. 2023).

Historically, hybridization and introgression were viewed as processes that impede speciation by homogenizing distinct genomes, introducing potentially maladaptive alleles, and disrupting locally adapted gene networks. However, they recently emerged also as key drivers of species diversification (e.g. Abbott et al. 2013; Seehausen 2014), with gene flow now regarded as an important process during speciation (He et al. 2019; Linck and Battey 2019; Mallet et al. 2016). Gene flow between recently diverged species can increase genetic diversity, offering the substrate for natural selection. Interspecific hybridization can prompt ecological diversification through heterosis and adaptive introgression, a process in which advantageous alleles are transferred between gene pools, enhancing adaptation. Furthermore, introgression can introduce advantageous alleles upon which selection has already acted or create novel allele combinations, thus catalysing evolutionary radiations (Abbott et al. 2013; Edelman et al. 2019; Harrison and Larson 2014; Meier et al. 2019; Suarez-Gonzalez et al. 2018).

The rapidly radiating and highly diverse Neotropical *Tillandsia* subg. *Tillandsia* provides a particularly relevant study system to investigate the drivers of radiations. Subgenus *Tillandsia* is the most diverse of seven subgenera within genus *Tillandsia* (family Bromeliaceae), comprising more than 250 predominantly epiphytic species. *Tillandsia* species are distributed across a wide range, from the southeastern United States to central Argentina (Barfuss et al. 2016). Members of the subgenus exhibit tremendous morphological diversity of adaptive traits which allows them to occupy disparate habitats, from tropical rainforests to deserts, and from lowlands to highlands (Benzing 2000; Givnish et al. 2011). Adaptations such as adventitious roots, trichomes modified for water and nutrient absorption, a water-impounding leaf rosette tank and Crassulacean acid metabolism (CAM) appear to have undergone in some cases correlated and in others contingent evolution, resulting in adaptive syndromes (Givnish et al. 2014).

Early efforts employing conserved plastid and nuclear markers circumscribed the monophyly of the subgenus and its clades, yet recovered largely inconsistent relationships between subclades (Barfuss et al. 2005; 2016; Chew et al. 2010; Granados Mendoza et al. 2017; Pinzón et al. 2016). Using plastome phylogenomics, Vera-Paz et al. (2023) confirmed the monophyly of the subgenus and the presence of three main clades within the subgenus, including a monophyletic, radiated ‘clade K’ with mostly epiphytic and ornithophilous members distinctly distributed in North and Central America (Fig. 1; Barfuss et al. 2016; Granados Mendoza et al. 2017). They further identified K.1 and K.2 as subclades of clade K. Within this study’s sampling, the *T. punctulata* (K.1) and *T. utriculata* (K.2.1) subclades include widespread or endemic species with intermediate CAM-C3 syndromes or various CAM-like photosynthetic syndromes, respectively. The *T. guatemalensis* subclade (K.2.2) is a small group of widespread and predominantly-C3 plants, exhibiting smooth leaves and a prominent tank habit. Finally, the *T. fasciculata* subclade (previously K.2.3) includes species largely exhibiting CAM photosynthetic syndrome with typical water-absorbing trichomes. In addition, species within this subclade K.2.3 present varying distributions, from endemic to widespread, some extending into the Antilles (Espejo-Serna et al. 2004; Till 2000).

**Figure 1.**
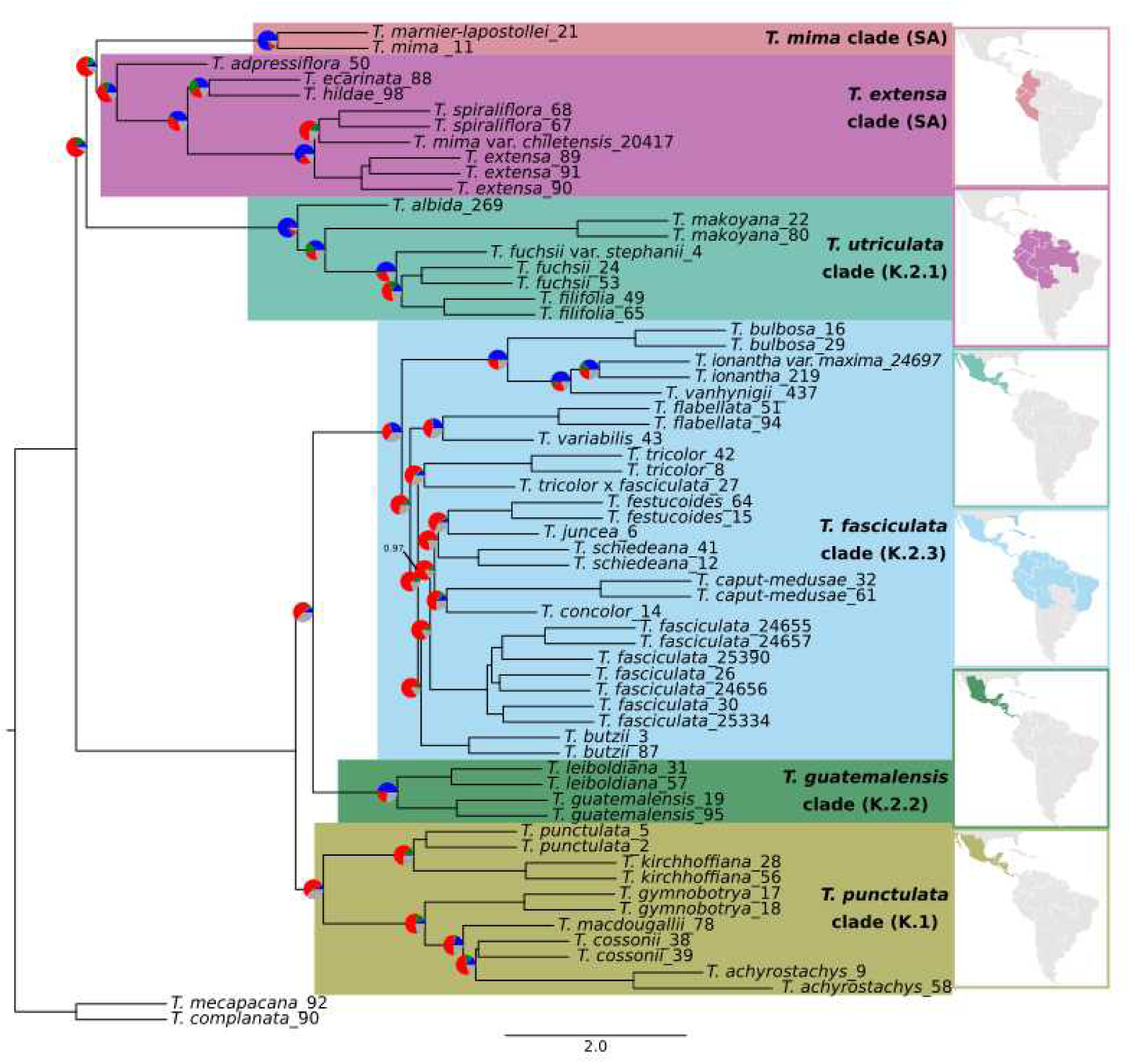
A coalescent-based species tree generated on 15,791 genomic windows with ASTRAL-III for 64 individuals representing 33 species of *Tillandsia* subg. *Tillandsia*, plus two outgroups of subg. *Allardtia*. Branch lengths are given in coalescent units. Node values represent local posterior probabilities for the main topology and are equal to one unless noted otherwise. Pie charts at the nodes show levels of gene tree discordance: the percentages of concordant gene trees (blue), the top alternative bipartition (green), other conflicting topologies (red) and uninformative gene trees (grey). Clade terminology follows previously proposed names (e.g., Pinzón et al. 2016 and Barfuss et al. 2016).

The subgenus *Tillandsia* was estimated to be phylogenetically young: existing Bromeliaceae phylogenies set the crown age of the subfamily Tillandsioideae at 15.2 ± 0.4 Mya and of “core tillandsioids” at 9.6 ± 0.7 Mya (Givnish et al. 2014). Several studies (Givnish et al. 2014; Till 2000; Vera-Paz et al. 2023; Winkler 1990) proposed that the ancestor of ‘clade K’ was most likely South American, expanded to Central America approximately 4.9 Mya and subsequently dispersed multiple times to North America. However, the relationships within ‘clade K’ remained elusive and disagreements between gene trees are common, calling to consider the role of different evolutionary processes in the *Tillandsia* radiation. As a result, hybridization was put forward as especially prevalent in the subgenus (Till 2000; Vera-Paz et al. 2023).

In contrast to phylogenetic reconstruction using a limited number of markers, whole genome sequencing provides an abundance of informative data, which increases resolution and helps disentangle major causes of incongruence (e.g., Liu et al. 2015; Guo et al. 2023). In comparison to plastid markers, nuclear loci are characterised by higher evolutionary rates and biparental inheritance, rendering them suitable for resolving rapid radiations and investigating hybridization (Léveillé-Bourret et al. 2018; McKain et al. 2018; Debray et al. 2019). Due to costs and the complexities of molecular lab work, a trade-off often exists between the scale of sequencing and the extent of taxon sampling in phylogenomic studies (Mitchell et al. 2000). Regardless, genome-wide molecular phylogenetics generally results in greater phylogenetic informativeness and improved accuracy (Rokas et al. 2003; Wortley et al. 2005; Léveillé-Bourret et al. 2018; Debray et al. 2019).

Using whole-genome resequencing data and a recently published reference genome, our study aims to shed light on the evolutionary history of *Tillandsia* subgenus *Tillandsia*, focusing on Central American groups within the radiated ‘clade K’ and representatives of South American species. Specifically, we seek to infer relationships between representative species and investigate signals of phylogenetic conflict, with the ultimate aim to examine patterns of gene flow which may have contributed to adaptive trait shifts within this Neotropical rapid radiation. Our detailed analyses reveal a reticulated history within the subgenus and at least two colonisation events from South into Central America, surprisingly indicating that the previously described ‘clade K’ (Barfuss et al. 2016; Granados Mendoza et al. 2017; Vera-Paz et al. 2023) may not constitute a monophyletic group.

## Materials and Methods

### Plant Material

Leaf material was collected from 69 individuals belonging to 36 *Tillandsia* species. Material from field collections was cut lengthwise and immediately dried in powdered silica gel. Samples from the botanical collection of the University of Vienna were extracted fresh (Supporting Tables S1, S3). 67 accessions correspond to 34 species of *Tillandsia* subg. *Tillandsia*, while the remaining two belong to subg. *Allardtia* sensu Smith & Downs (1977) and were used as outgroups. Of the 34 ingroup species, 26 represent the Central-North American ‘clade K’ radiation and eight species represent South American *Tillandsia* (henceforth: SA clades; Supporting Fig. 1, Supporting Table S1). Albeit not exhaustive, our sampling was performed with the intention to represent the variety of morphological and physiological syndromes within clades SA and ‘K’ (Supporting Table S2). We sampled species from all subclades of clade K previously introduced: 6, 5, 2, and 13 from subclades *T. punctulata*, *T. utriculata*, *T. gutemalensis* and *T. fasciculata*, respectively.

### DNA Library Preparation

DNA extractions were performed as previously described (Yardeni et al. 2022) using a modified CTAB protocol (Doyle and Doyle, 1987) or the QIAGEN DNeasy® Plant Mini Kit (Qiagen, USA). They were further purified using the Nucleospin® gDNA cleanup kit from Macherey-Nagel (Hudlow et al. 2011). The extracts were subsequently diluted in water and quantified with a Qubit® 3.0 Fluorometer (Life Technologies, Carlsbad, CA, USA). Prior to library preparation, a maximum of 600 ng of DNA per accession was sheared at 4°C to a target average length of 400 bp using a Bioruptor® Pico sonication device (Diagenode, Denville, NJ, USA). Illumina libraries were prepared using a modified KAPA protocol with the KAPA LTP Library Preparation Kit (Roche, Basel, Switzerland) with adaptor ligation and adaptor fill-in reactions based on Meyer and Kircher (2010). Some libraries have been instead prepared with the NEBNext Ultra II DNA PCR-free Library Prep Kit (New England Biolabs, Ipswich, MA, USA). Samples were either double indexed with a set of 60 dual-index primers, as recommended by Kircher et al. (2012) and described in Loiseau et al. (2019), or with Illumina TrueSeq PCR-free dual indexes. The libraries underwent size selection steps using AMPure Beads (Agencourt). Finally, the libraries were pooled and sequenced at the Vienna BioCenter Core Facilities on Illumina HiSeqV4 PE125 or on NovaSeqS1 PE150. Protocol details for all accessions are listed in Supporting Table S3.

### Data Processing

The raw sequence data was demultiplexed using deML v.1.1.3 (Renaud et al. 2015) and bamtools v.2.5.1 (Barnett et al. 2011), and converted from BAM to FASTQ format using bedtools v.2.29.2 (Quinlan and Hall 2010). Reads were trimmed for adapter content and quality using trimgalore v.0.6.5 (Krueger 2019), a wrapper tool around fastqc and cutadapt, using --fastqc --retain unpaired. Sequence quality and adapter removal were confirmed with FastQC v.0.11.9 (Andrews 2010).

Quality- and adapter-trimmed reads were aligned to the *T. fasciculata* reference genome v.1.0 (GenBank Assembly accession GCA_029168755.1, Groot Crego et al., 2024) using bowtie2 v.2.3.5.1 (Langmead and Salzberg 2012) with the --very-sensitive-local option to maximise alignment rates. Low-quality mapped reads (mapQ < 10) were removed, alignments were sorted by position using samtools v.1.15.1 (Li et al. 2009), and PCR duplicates were marked using MarkDuplicates from PicardTools v.2.25.2 (Picard Toolkit 2018). The average coverage was calculated with samtools v.1.15.1 (Li et al. 2009).

To call variants, we used GATK HaplotypeCaller v.4.1.9.0 followed by joint calling with GenotypeGVCFs (Poplin et al. 2018). We proceeded to confirm technical replicates and to identify high relatedness between accessions with kinship coefficients inferred through KING v.2.2.6 (Manichaikul et al. 2010). BAM files of technical replicates were merged before we called a final variant call file (VCF). Samples exhibiting a first degree relationship with another sample (consistent with full-sib or direct parent-offspring relationships; i.e., kinship > 0.177) or having an average genome-wide coverage below 3.5x were removed from subsequent analyses (Supporting Table S3). The resulting VCF was filtered using bcftools v.1.15 (Danecek et al., 2021) and GATK SelectVariants (DePristo et al. 2011), to exclude indels and any SNPs located within 3bp of indels. Regions annotated as transposable elements were excluded using bedtools intersect (Quinlan and Hall 2010). We generally used the transition/transversion ratio for guidance to define our filtering parameters, aiming for values leading to a set of SNPs exhibiting the highest ratio. Ts/Tv ratios were calculated with SnpSift (Cingolani et al. 2012). We finally used the following parameters for filtering: mapping quality (MQ) < 15, read depth coverage (DP) < 4, quality by depth (QD) < 4, Fisher strand bias (FS) > 40, strand odds ratio (SOR) > 3, minor allele frequency (MAF) < 0.045 (corresponding to the presence in at least six chromosomal complements) and missing rate < 0.2. We prepared additional files with different MAF filtering thresholds, to examine the possible bias in branch lengths (see below and supporting file 3). Summary statistics were generated using bcftools stats (Danecek et al., 2021). The full variant calling pipeline is available at https://github.com/giyany/TillandsiaPhylo/notebooks/raw_to_vcf_GATK.md.

### Phylogenetic Tree Inference

We inferred phylogenetic relationships for all samples using both a maximum-likelihood (ML) and a coalescent-based method. We included a coalescent-based method in order to explicitly account for ILS, which may otherwise result in high support for an incorrect topology (Kubatko and Degnan 2007). Gene tree incongruence further provides insight into molecular genome evolution, including the extent of incomplete lineage sorting and genomic processes such as hybridization and introgression (Galtier and Daubin 2008; Hibbins et al. 2020; Morales-Briones et al. 2021; Wendel and Doyle 1998). Both analyses were repeated on data-sets with different MAF filtering thresholds to account for possible bias in branch lengths (supporting files 3). The ML tree was inferred on a concatenated supermatrix, including both variant and invariant sites. First, a concatenated matrix was obtained by converting the VCF into a phylip file with vcf2phylip (Ortiz 2019). We then inferred a phylogeny with IQ-TREE v.2.1.3, using IQ-TREE’s ModelFinder to select the best-fitting partitioning scheme and models for each genic region (Nguyen et al. 2015, Kalyaanamoorthy et al., 2017). Node support was inferred with 1,000 nonparametric ultrafast bootstrap replicates (Chernomor et al. 2016). During the first inference, the available *T. zoquensis* accession grouped within *T. fasciculata* (Supporting Fig. S2), suggesting a need to revise its species status. We removed this sample from the data-set and all analyses hereafter. We then repeated the analysis, constructing a whole-genome ML tree as described, as well as separate ML trees for each reference chromosome.

To reconstruct a species tree, ML trees were estimated on non-overlapping genomic windows of 10 kb. Windows with fewer than 40 SNPs were excluded. Since loci with insufficient signal may reduce the accuracy of species tree estimation (Mirarab 2019), nodes with a bootstrap support below ten were collapsed across all trees with Newick utilities v.1.1.0 (Junier and Zdobnov 2010). ASTRAL-III v.5.7.8 (hereafter: ASTRAL) was then used to infer a species tree, measuring branch support as posterior probabilities (Zhang et al. 2018). To explore gene tree discordance, we used phyparts and calculated quartet support for the main, as well as the first and second alternative topologies. We calculated the number of concordant and discordant bipartitions on each node, imposing a cutoff of 70 for informative bootstrap values (-s 70; Smith et al. 2015). Gene tree discordance was visualised with phypartspiecharts.py (available from https://github.com/mossmatters/MJPythonNotebooks).

### Investigating Monophyly and Deviations from a Bifurcating Tree Structure

We used several variations of the ABBA-BABA test (D-statistic) to further explore the relationships within and between subclades and to assess deviations from tree structure (Durand et al. 2011; Green et al. 2010; Martin et al. 2015). The D-statistic we implemented in Dsuite v.0.5r45 (Malinsky et al. 2021) specifically uses allele frequency estimates, which allows to include several individuals per taxon and does not require the implicit assumption that the outgroup is fixed for the ancestral allele. For all D-statistic-related analyses we used the full VCF file with *T*. *complanata* as an outgroup. Significance was assessed using a jackknife procedure with 200 kb window size and family-wise error rate (FWER) was calculated and corrected for multiple comparisons following the Holm–Bonferroni method with the p.adjust function in R (R Core Team 2020). We first examined the consistency of assigning individuals to a subclade, following the approach of Malinsky et al. (2018), by testing whether individuals assigned to the same subclade always share more derived alleles with each other than with individuals from other subclades. For each individual A, we calculated D(A,G_1_;G_2_,O), where A is assigned to one subclade and G_1_ and G_2_ to another. Values greater than zero indicate that allele sharing is greater within groups compared to across groups.

To characterise the relationships among subclades, we examined the consistency of subclade monophyly in genomic windows. We used the set of window-based trees previously generated for the ASTRAL analysis and checked if a subclade was monophyletic in each window with the check_monophyly function in the ETE Toolkit v.3 (Huerta-Cepas et al. 2016). Next, a cloudogram of ML trees was visualised. Window-based trees were modified using R packages phytool v.1.0-3 (Revell 2012) and treeio v.1.20.0 (Wang et al. 2020) to force branch lengths into an ultrametric structure for visualisation purposes. A cloudogram was generated with ggdensitree from the R package ggtree v.3.4.0 (Yu 2020). We next used D_min_, another variation of the D-statistic (Malinsky et al. 2018), to assess if allele-sharing among subclades is consistent with a tree-like structure. D_min_ yields a conservative inference of gene flow within a trio by providing the minimal absolute value of D across all possible arrangements within it. A significantly positive score signifies allele sharing that is inconsistent with a single species tree. To identify and visualize the genetic structure between subclades, we performed a principal components analysis (PCA) on a reduced data-set. This data-set was pruned for linkage disequilibrium by randomly selecting SNPs that are at least 10 kb apart among all the biallelic SNPs with <10% missing data. The PCA analysis was performed with SNPRelate v.1.20.1 (Zheng et al. 2012).

Finally, to quantify and visualise the relationship among subclades along the genome we used topology weighting by iterative sampling of subtrees with *Twisst* v.0.2 (Martin and Van Belleghem 2017). Using ML topologies inferred on genomic windows, this method considers all possible topologies and quantifies the contribution of each to the full tree, enabling to locate genomic regions that are associated with certain topologies. We focused on the three largest subclades and reduced the number of possible topologies by including only the subclades *T. extensa*, *T. utriculata* and *T. fasciculata*. For this analysis, subtrees were constructed on windows of 50 SNPs following the strategy of Martin and Van Belleghem (2017). Individual VCF files for each window were produced by partitioning the original VCF with the biostar497922 script from Jvarkit (Lindenbaum 2015) and ML trees were inferred for each window as described previously. A summary and visualisation of all topologies along genomic coordinates were produced with a slightly modified version of the plot_twisst.R script, which excludes windows with high rates of missing data in the computation of the average regional weightings (Martin and Van Belleghem 2017; script is available at https://github.com/giyany/Tillandsia_Phylo_hybridization/blob/main/notebooks/ plot_twisst_mod.R).

### Characterising Hybridization Events

To quantify the rates of hybridization between all species in our dataset, we obtained a genome-wide signal of hybridization using the original implementation of the D-statistic and estimates of admixture fraction f (henceforth: ƒ_4_-ratio) in *Dsuite* v.0.5r45 (Malinsky et al. 2021). Given a certain level of uncertainty regarding the true relationship between species, we set no *a priori* knowledge of taxon relationships. Instead, *Dsuite* ordered each trio so that the BBAA pattern is more common, principally to focus on topologies with minimal discordant patterns. For all D-statistics related analyses below, we again used the full VCF file with *T*. *complanata* as an outgroup. We repeated the analysis separately for each reference chromosome and computed the *D*-statistic for each, obtaining p-values using a jackknife procedure with windows of 150 SNPs.

Since groups that are involved in hybridization may share branches on a phylogenetic tree, a single hybridization event can present multiple correlated instances of a significantly elevated D-statistic. This is especially expected when gene flow involves ancestral lineages, affecting internal branches of a phylogenetic tree. We used *Dsuite* to calculate the *f*-branch metric (henceforth: f_b_(C)), an estimator developed to create a summary of gene flow events with minimized correlation (Malinsky et al. 2021). The f_b_(C) results invited several hypotheses regarding hybridization events, which we examined by inferring a species network under a maximum pseudo-likelihood approach using PhyloNet v.3.8.0 (Cao et al. 2019; Than et al. 2008; Yu and Nakhleh 2015). For PhyloNet, we reduced our sampling to one outgroup and 18 ingroup taxa to include representatives from each highly-supported subclade, then inferred ML trees on non-overlapping windows of 10 kb as previously described. We inferred networks that specified between zero and five reticulation events and repeated the search five times for each network, finally picking the network with the highest pseudo-likelihood score in each search.

### Characterising Hybridization on Chromosome 18

Our previous calculation of the D-statistics for each reference chromosome (see above) pointed at significantly elevated values on chromosome 18. Gene flow particularly involved the species *T. achyrostachys* (*T. punctulata* subclade) and most species in the CAM subclade *T. fasciculata* (see results). Species in the *T. punctulata* subclade putatively possess an intermediate C3-CAM photosynthetic syndrome, expressing intermediary values of stable carbon isotope ratios (δ^13^C; Supporting Table S2, but see also Messerschmid et al. 2021). Within our sampling of this subclade, *T. achyrostachys* was documented to express the strongest CAM phenotype (δ^13^C of −14.7; Crayn et al. 2015). Additionally, all known species in the *T. fasciculata* subclade are putatively strong, constitutively CAM plants. We wished to investigate the possibility that gene flow introduced advantageous genes related to shifts in metabolic syndromes. Hence, we inferred the signature of hybridization along chromosome 18 to locate highly admixed loci. We focused on possible hybridization between *T. punctulata*, *T. butzii* and *T. achyrostachys* (corresponding to the *T. punctulata* subclade, the *T. fasciculata* subclade and the *T. punctulata* subclade, respectively). This allowed us to account for phylogenomic relatedness between *T. achyrostachys* and *T. punctulata* and for similar photosynthetic syndromes between *T. achyrostachys* and *T. butzii*. We used the Dinvestigate function from *Dsuite* on this species trio, performing analysis on windows of 50 SNPs with a step size of ten, obtained as previously described. The D-statistic itself shows large variance when applied to genomic windows (Martin et al. 2015), hence we used f_dM_, a statistic designed to investigate hybridization in small windows which also accounts for allele sharing across all possible taxon arrangements in a trio (Malinsky et al. 2015; Martin et al. 2015). We identified regions exhibiting D-statistic values exceeding the 95% quantile of the distribution and inspected the genes annotated in regions of high admixture, specifically their functional annotation.

## Results

### Read Mapping and Variant Calling

After removing samples with low coverage, those sharing high kinship coefficients, and the *T. zoquensis* accession (see methods) we retained 64 *Tillandsia* accessions, corresponding to 35 recognized species. The average number of reads retained per accession was 5.1x10^7^ (range 4.7x10^6^-1.2x10^8^, SD=2.5x10^7^) and average mapping rates were 89.6% (range 69.4%-97.5%, SD=6.3), with slightly higher rates for members of the *T. fasciculata* subclade and slightly lower for the SA subclades (97.5% and 89.6% respectively). Differences in mapping rates between the six main subclades (Fig. 1) were however not significant (Kruskal–Wallis test, p-value = 0.22), suggesting no or limited biases towards the reference genome. An average coverage of 13.8x (range 4.1x-35.0x, SD=6.5x) was obtained for the samples retained further. After variant calling and filtering we retained 2,162,143 high-quality SNPs.

### Phylogenomic Inference Contradicts clade Monophyly

The concatenated matrix was partitioned into 14,392 regions, whereas the coalescent-based analyses were performed on 15,791 genomic windows. The species tree (Fig. 1) and concatenated maximum-likelihood (ML) tree (Supporting Fig. S1) yielded inconsistent results regarding the main clades and relationships. In both trees, the Central American ‘K’ subclades did not form a monophyletic group, contrary to among-subclade relationships previously reported for this subgenus (Barfuss et al. 2016; Granados Mendoza et al. 2017; Rivera 2019; Vera-Paz et al., 2023). Specifically, our results are incongruent with a monophyly of the previous inferred ‘subclade K.2’. Instead, the *T. utriculata* subclade (K.2.1) was recovered as a sister to one or both of the South American subclades (in the ML tree or the species tree, respectively). The assignment of species to subclades and all other relationships between subclades remained overall congruent with previous phylogenies (Granados Mendoza et al. 2017; Vera-Paz et al., 2023). Several, relatively minor within-clade topological differences appeared among trees inferred with different methods, especially within the *T. fasciculata* subclade. Relationships between species in this subclade were highly supported yet coupled with high levels of gene tree incongruence, suggesting substantial allele sharing and high rates of gene flow within this group. While the South American taxa in our sampling formed a monophyletic group in the species tree (Fig. 1), in the ML tree (supporting Fig. S1) these were separated into two subclades: one consisted of endemic Peruvian species and the widespread species *T. adpressiflora*, while a second subclade contained *T. marnier-lapostollei* and *T. mima*. The latter was retrieved as paraphyletic to the *T. utriculata* subclade and all remaining South American taxa. As the subclades we retrieved do not correspond to the previously-inferred *T. paniculata* and *T. secunda* clades (Vera-Paz et al. 2003), we renamed them as *T. extensa* and *T. mima* subclades, respectively (supporting table S1). We additionally renamed the Mexican ‘K’ subclades to reflect the new assignment (Fig. 1). Gene tree discordance was widespread within the dataset, affecting both deep and shallow nodes (Fig. 1; Supporting Fig. S3). The relationships between the South American species and the *T. utriculata* subclade were particularly characterised by short internode distances and many alternative topologies. Frequently, the majority of inferred gene tree topologies were discordant with a single main topology: for example, in the node preceding the separation of the SA subclades only 3,148 (19.9%) of all gene windows supported the main topology (Fig. 1; Supporting Fig. S3). Similar levels of discordance characterised the internal nodes within the *T. fasciculata* subclade and high levels of discordance were also found within the *T. punctulata* subclade (Fig. 1; Supporting Fig. S3).

Maximum-likelihood trees constructed separately for concatenated matrices of each reference chromosome retrieved many different topologies: solely considering relationships between subclades we recognized ten different topologies among the different 25 trees (see Supporting file 1). The placement of the *T. mima* subclade showed the greatest incongruence between trees, as well as the placement of *T. adpressiflora*. In three trees, the relationships between subclades were similar to the topology retrieved from the whole-genome concatenated matrix. The South American species formed a monophyly in six chromosome trees and ‘clade K’ was recovered as monophyletic in two. While each chromosome tree contains high amounts of gene tree discordance, the abundance of different topologies imply that several evolutionary histories can be traced along the genome of *Tillandsia*, as disparate genomic processes contribute to the differences between gene trees and the true species relationships. Trees inferred on data-sets with different MAF filtering thresholds did not substantially differ in topology, whereas gene tree discordance was minimally affected (supporting file 3). Branch lengths were however affected, as minor alleles contribute to long branches of specific species. We again retrieved inconsistency in the placement of the *T. mima* subclade when using different filtering parameters, indicating the subclade cannot be confidently placed in our current data-set.

### Lack of Monophyly and Deviations from Tree-like Structure

The multitude of tree topologies along the genome led us to hypothesise that a single bifurcating tree misrepresents the true relationships among species in the subgenus *Tillandsia*, in particular affecting ‘clade K’ (Supporting Fig. S4). To investigate deviations from a tree-like structure, we analysed the patterns of allele sharing between species and between subclades. Assuming no interspecific gene flow, allele sharing is expected to be consistent with the main tree topology, whereas asymmetrical allele sharing indicates deviations from that assumption.

We first used the D-statistic framework to test the robustness of the assignment of species to subclades. We calculated allele sharing between species, utilising the D-statistic’s allele frequency estimates. We found high support for subclade monophyly, as species assigned to the same subclade shared more alleles with other species within the subclade than with species from other subclades. However, we observed low support for subclade monophyly based on the analyses in genomic windows performed with ETE: 52.4%, 34.4% and 14.11% of the windows supported monophyly for subclades *T. guatemalensis*, *T. fasciculata* and *T. punctulata*, respectively. A notable exception was observed for *T. utriculata*, for which 86.6% of the windows were consistent with a monophyly.

We next characterised allele sharing between subclades by computing the D_min_ statistic on a total of 7,141 trios (Supporting Fig. S5a). We found widespread deviations from a tree-like structure, mostly driven by allele sharing between Central American taxa (previous ‘clade K’): D_min_ values were significantly elevated in 3,650 comparisons (P<0.05) and more than 95% of those were highly significant (P<0.01). The rate of significant D values was highest for comparisons involving accessions assigned to *T. utriculata* subclade (58.7% of the trios), followed by *T. punctulata*, *T. guatemalensis* and *T. fasciculata* (57.9%, 56.1% and 51.4%, respectively). We also employed a multivariate technique with PCA to investigate the interspecific genetic structure (Supporting Fig. S5b). After distance-pruning and removing SNPs with <10% missing data we retained 16,204 SNPs. We found a consistent interspecific genetic structure, separating the South American subclades and the *T. utriculata* subclade, while the remainder of the Central American subclades clustered densely - reflecting the distinct subclades found in tree-based and D-statistics approaches.

Finally, to obtain a better understanding of how subclade relationships vary along the genome, we considered three distinct tree topologies for topology weighting with *Twisst* (Supporting Fig. S6): the most frequent topology was congruent with the one recovered in the species tree, which placed the *T. extensa* subclade as sister to the *T. utriculata* subclade in 42% of the genomic windows. A second topology, with *T. fasciculata* and *T. utriculata* as sister subclades, was recovered from 35% of the genomic windows, while a third topology appeared in 23%. The three topologies were broadly equally distributed along the genome – however, in some chromosomes (for example, chromosome 4; Supporting Fig. S6) the first topology was more frequent within centromeric regions. Given that topology weighing remains a descriptive method, it does not allow explicit testing for hybridization or ILS. Regardless, the prevalence of the main topology in regions of low recombination rates and reduced genic density suggests that it represents the backbone phylogeny, while other topologies are likely the result of gene flow or deep coalescence (Carneiro et al 2009; Christmas et al 2021).

### Correlated and Widespread Gene flow Events

D-statistics results for all possible trios indicated that all subgenus *Tillandsia* species in our dataset were involved in potential hybridization (Supporting Fig. S7). Out of 7,141 tests in total, 4,331 returned significantly elevated values, with D values ranging between 0.021 and 0.581. The signal was not localised to a specific chromosome according to separate calculations on each chromosome (Supporting file 2). A prominent signal revealed gene flow between species in the *T. utriculata* subclade and all other Central American subclades. Notable gene flow signals were also found within the *T. fasciculata* subclade. ƒ4-ratio scores ranged between 0.0017 and 0.357, but for most hybridization events the proportion of the genome involved was estimated smaller than 10% (Supporting Fig. S8). Larger parts of the genome were admixed in hybridization within the main subclades: for example, *T. caput-medusae* was involved in hybridization events with *T. butzii* and T*. fasciculata* involving ca. 31.7% of the genome.

Further analysis confirmed that past hybridization events involved the ancestor of the *T. urticulata* subclade and an ancestor of the other three Central American clades. In analysis based on the f-branch metric, elevated f_b_(C) scores were assigned to events involving these subclades with an average of 6.3% (Fig. 2). Among all between-subclades f_b_(C) values, 246 (19.6%) were significantly elevated (P<0.05), although most significant values (412, 62.6%) occurred within subclades. In a network analysis on a total of 15,535 genomic windows, the results repeatedly indicated the involvement of the *T. utriculata* subclade in hybridization with the *T. extensa* and other Central American subclades (Fig. 3). Notably, higher weights were assigned to the former hybridization. Additional gene-flow events occurred within subclades. Analyses allowing for four or more reticulation events resulted in events involving the outgroup or fewer than four events reported, so we present here results for up to three reticulations. Overall, these findings confirm the extensive violations of a tree-like structure revealed in the previous part of the analyses and suggest that inter-specific gene flow occurred frequently throughout the evolutionary history of the radiation.

**Figure 2.**
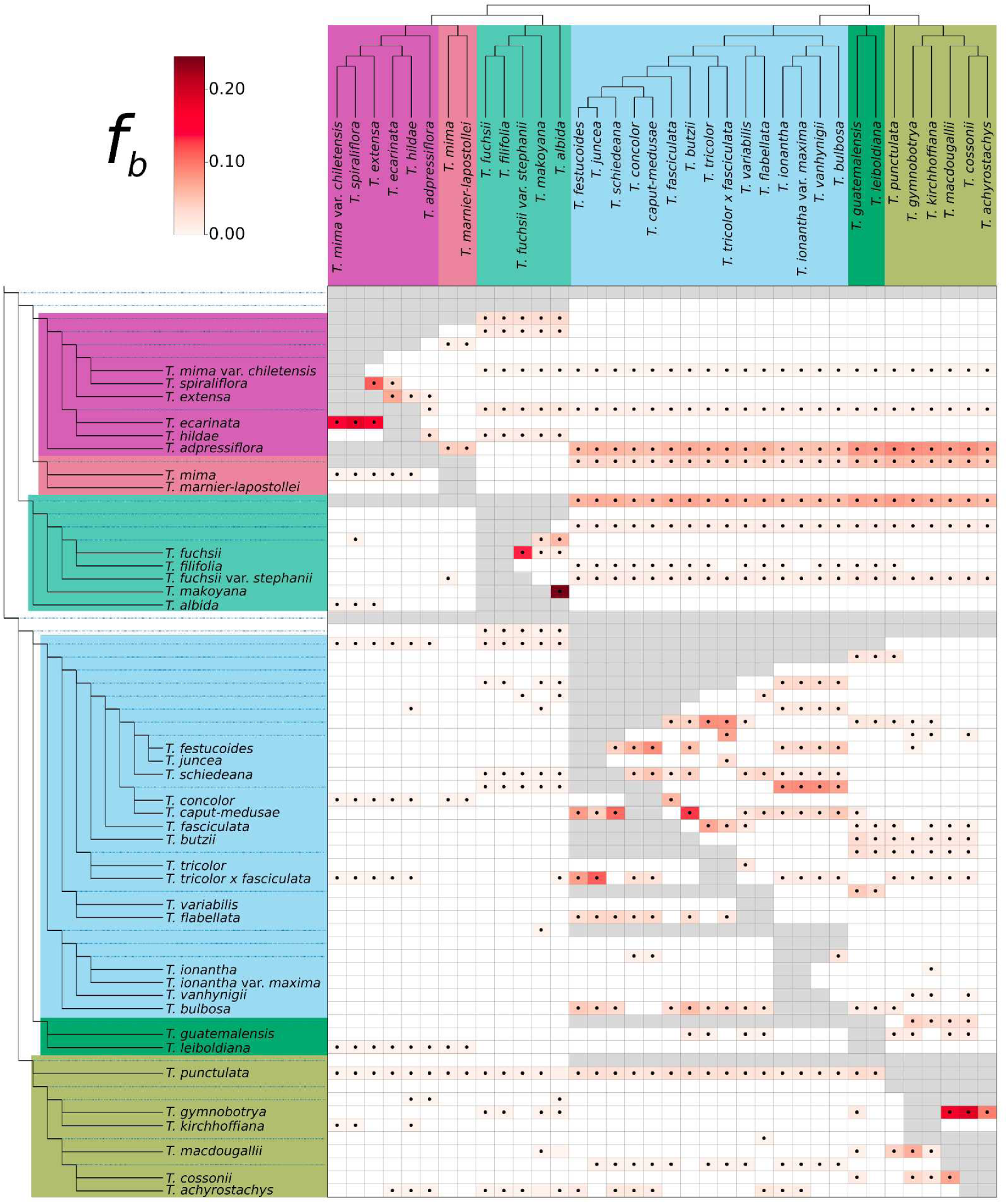
Heatmap summarising the statistic f_b_(C), where excess sharing of derived alleles is inferred between the branch of the tree on the Y axis and the species C on the X axis. The ASTRAL species tree was used as input topology for the branch statistic. The matrix is colored according to f_b_(C) values and grey squares correspond to tests that are inconsistent with the ASTRAL phylogeny. Dots within the matrix denote a significant p-value, estimated using a block jackknife procedure and corrected for family wise error rate. Colours correspond to the clades in Figure 1.

**Figure 3.**
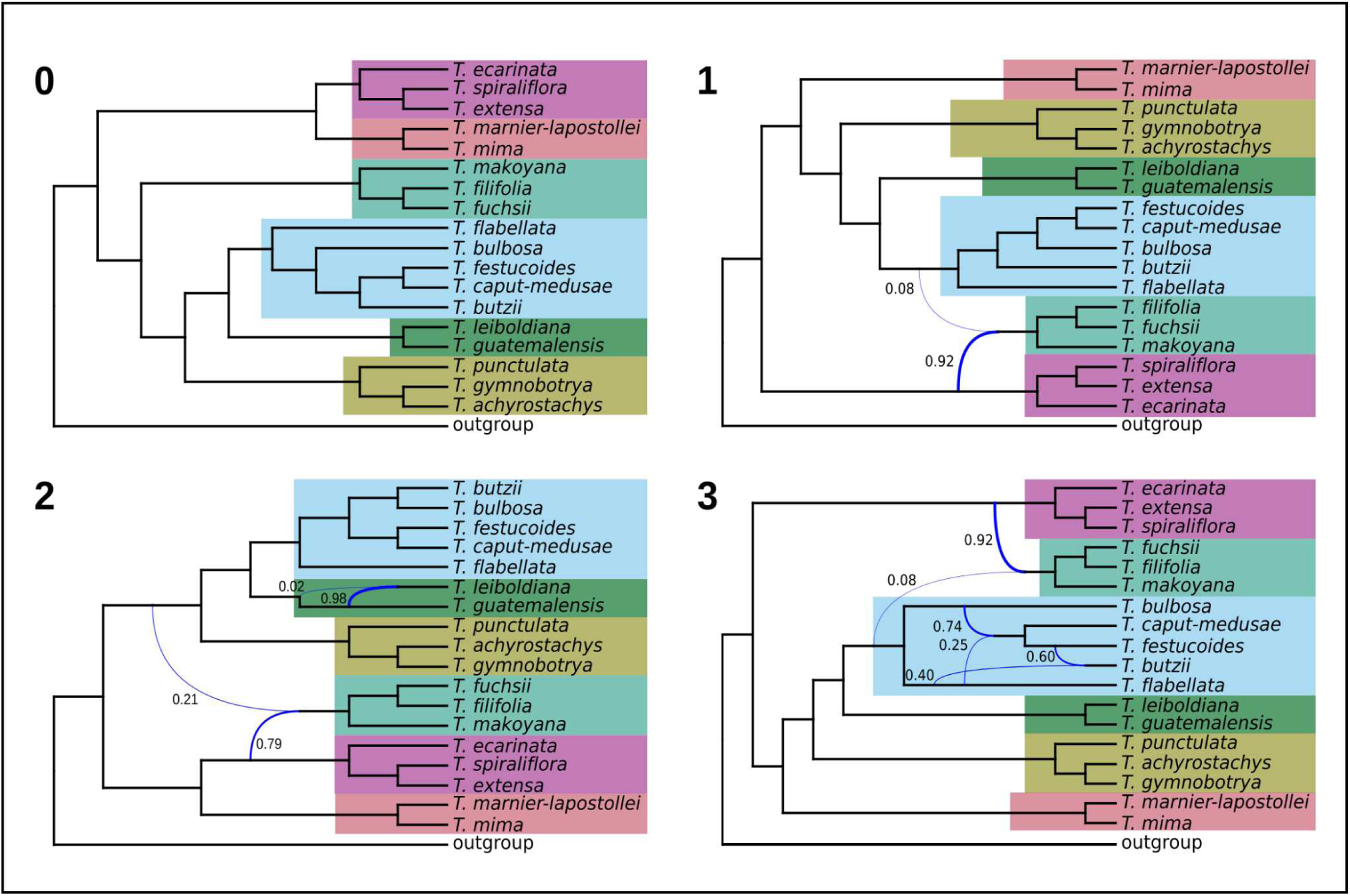
pseudo-likelihood species networks inferred with PhyloNet for zero to three reticulation events (network presented was scored with the highest log-probability). Curved branches indicate reticulation events. Numbers next to curved branches indicate inheritance probabilities for each event. Colours correspond to the clades in Figure 1.

### Local Footprint of Hybridization between CAM-expressing Species

Elevated D-statistics, signifying hybridization, generally affected all chromosomes (Supporting file 2). In contrast, a significantly elevated, but localized D-statistic was found on chromosome 18 involving *T. achyrostachys* (*T. punctulata* subclade) and most of the species in the CAM subclade *T. fasciculata* (Supporting file 2). To further investigate this highly localised signal, f_dM_ statistics were calculated on a total of 9,185 windows between *T. punctulata*, *T. butzii* and *T. achyrostachys*. The first and last species are assigned to the *T. punctulata* subclade whereas *T. butzii* is a member of the *T. fasciculata* subclade. Across all chromosomes, mean values were negative at an average of -0.043 fitting the expectation for higher rates of gene flow between closely related species (Fig. 4a). Chromosome 18 was however characterised by two regions of high positive values, which were also evident with topology weighting (Fig. 4b, 4c). Considering centromere localization, we found that the loci exhibits high f_dM_ values, coincides with low genic density and contains 194 genes (Fig. 4d). In a detailed survey of gene annotations, we found this set of genes was potentially associated with a variety of stress and metabolic functions. For example, serine/threonine-protein kinase prpf4B is known to have a role in pre-mRNA splicing in yeast and humans (Eckert et al. 2016), and was found to be associated with stress response in millet (Parvathi et al. 2019). Another example is the S-adenosyl-L-methionine-dependent methyltransferase superfamily protein (SAM-Mtase), a key enzyme in plant metabolic pathways like the phenylpropanoid and flavonoid pathway (Joshi and Chiang, 1998; Sistla and Rao 2004).

**Figure 4.**
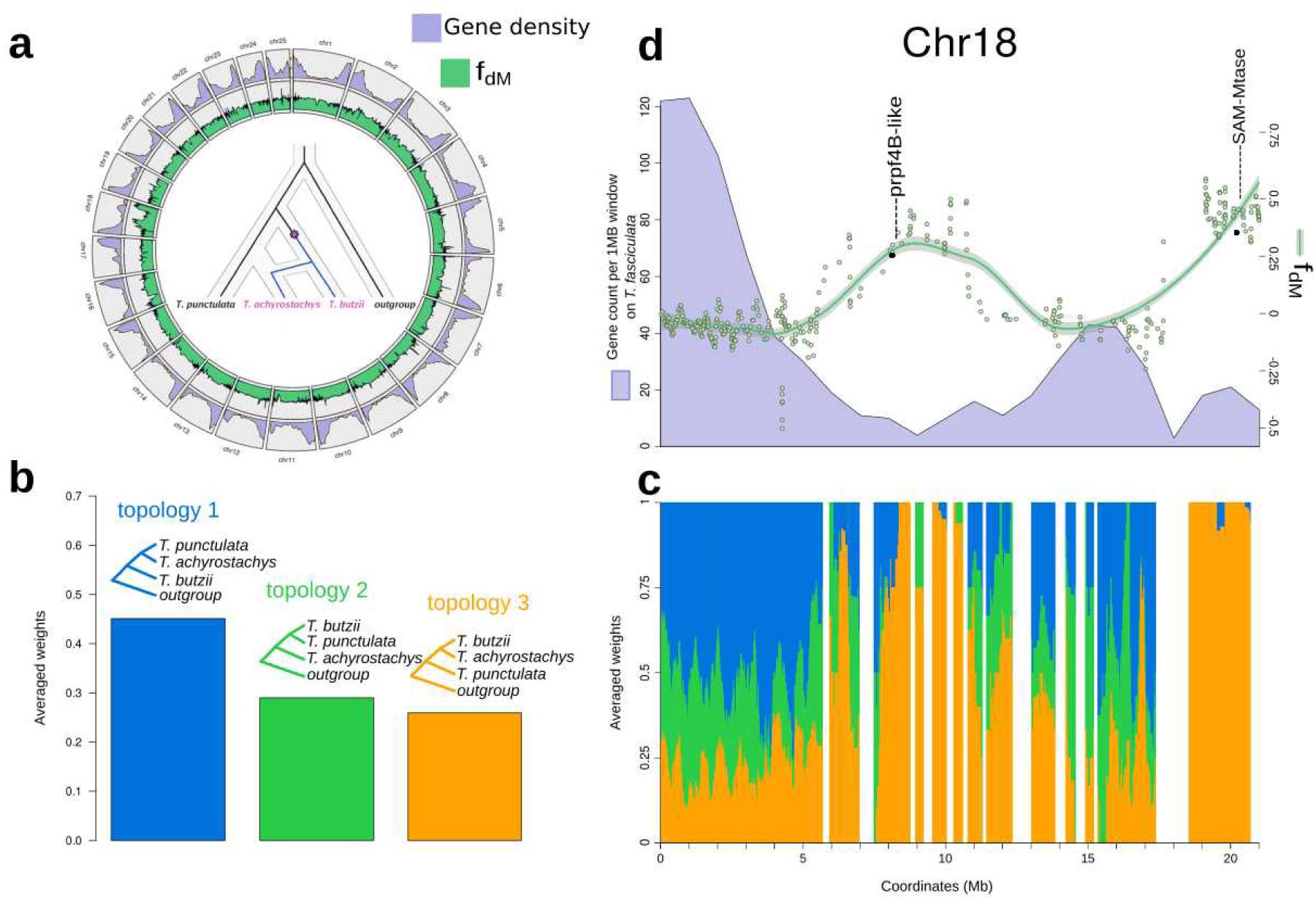
Signature of introgression and topology weighting on chromosome 18 between *T. punctulata*, *T. butzii* and *T. achyrostachys* (P1, P2 and P3; T. *complanata* was used as the outgroup). **A**, values of f_dM_ statistic and gene content for each chromosome calculated between *T. punctulata*, *T. butzii* and *T. achyrostachys*. **B**, Topology weighting by iterative sampling of subtrees in genomic windows of 50 SNPs using *Twisst*. Colours represent the frequency of each topology in **C** along chromosomal position in chromosome 18: white gaps indicate regions excluded due to high levels of missing data. **D,** f_dM_ statistic (green dots and smoothed green line, scale on the right) calculated in genomic windows. Analysis was performed on windows of 50 SNPs with a step size of ten. Shared variation is quantified in positive values when shared between P2 and P3 and as negative values when shared between P3 and P1. Gene content as the number of predicted genes per 1 MB window is in lavender with scale on the left.

## Discussion

Hybridization is extensively studied for its various roles in plant speciation. Using a multi-species genomic dataset and phylogenomic approaches, we present the subgenus *Tillandsia* as a striking example of a highly reticulated radiation. The radiation likely proceeded in the presence of rampant inter-specific gene flow, rather than by refined reproductive barriers (Loiseau et al. 2019; Koch et al. 2022; Vera-Paz 2023). An abundance of sequencing data tends to generate high branch support in both concatenated alignments and a coalescent-based tree-building method (Pease et al. 2016; Salichos and Rokas 2013), yet a detailed investigation in subgenus *Tillandsia* revealed that a bifurcating tree offers an incomplete picture of the true relationships between species. This work thus provides a compelling example to the mounting evidence of the presence, and possibly contribution, of gene flow in species diversification (Arnold et al. 2016; Filiault et al. 2018; Keller et al. 2013; Nosil 2008; Seehausen et al. 2014; Slovak et al. 2023; Stankowski et al. 2019; Zhang et al. 2021).

In stark contrast to previously inferred phylogenies, we retrieved the Central American ‘clade K’ as polyphyletic. First inferred based on morphological characters and later on a limited number of genomic markers, the generic and sub-generic classification of subfamily Tillandsioideae in general and of the genus *Tillandsia* in particular shifted throughout decades of phylogenetic research. Overall, phylogenies based on plastid sequences and on relatively few nuclear genomic regions offered little resolution for shallow phylogenetics in this young genus, particularly within Central American taxa (Pinzón et al. 2016; Barfuss et al. 2016; Granados Mendoza et al. 2017). Recently, a full plastome phylogeny greatly improved the phylogenomic resolution while statistical support remained low in shallow nodes (Vera-Paz et al. 2023). We suggest that incongruence between *Tillandsia* phylogenies reflects both the local scarcity of genomic divergence within this young radiation and the complicating role of hybridization. Moreover, considering the prevalence of recent inter-specific gene flow, the monophyly of the Central American subclades in plastid phylogenies seems particularly driven by chloroplast capture (Pham et al. 2017). Beyond insight on subclade relationship, our analyses produced several novel insights regarding cryptic species: for example, our inference suggests that *T. mima* and *T. mima var. chiletensis* are distinct species, despite morphological similarities. Similarly, *T. fuchsii* and *T. fuchsii* var. *stephanii* did not form a monophyletic group, raising a need to revise their taxonomic status (Fig. 1; Supporting Fig. S1).

Hybridization played a central role in the evolution of Central American *Tillandsia*, reflected in departures from tree structure encompassing all clades, subclades and species. D-statistics revealed both recent and ancient gene flow while allele sharing did not compromise species or subclade boundaries. Notably, the obtained f_b_(C) ranges were in general higher than those previously inferred for ancestral introgressions between species of Malawi cichlids (Malinsky et al. 2018) or of hares (Ferreira et al. 2021), but comparable to other rapid radiations (De-Kayne et al. 2022; Slovak et al. 2023). The interplay of genomic signals and our limited sampling complicate inference on the timing of gene flow events, yet their high prevalence suggests occurrence during both ancestral and recent history.

High rates of gene flow are compatible with *Tillandsia* ecology and evolution: *Tillandsia* is known to produce natural hybrids (Luther 1985; Koch et al. 2022; Till 2000) and bears copious seed adapted to wind dispersal, which may facilitate high rates of inter- and intra-specific gene flow (Mondragon and Calvo-Irabien 2006; Victoriano-Romero et al. 2017). Previous studies proposed that the South American ancestor of *Tillandsia* subg. *Tillandsia* colonised North and Central America in a long-distance dispersal event 4.86 Mya (Barfuss et al. 2016; Granados Mendoza et al. 2017; Vera-Paz et al. 2023) and our results offer intriguing hypotheses on multiple, likely polytopic origins of Central American *Tillandsia*. Instead of a single dispersal event, South American ancestors may have dispersed in several migration events into Central America. Strong population bottlenecks, producing phylogenies with relatively long internal branches, were followed by concomitant episodes of gene admixture and increased isolation. It is further possible that gene flow increased allelic diversity in founder populations, thus fueling adaptation (see below).

Apart from hybridization, other molecular processes can contribute to violations of a strictly bifurcating species tree, such as ILS, paralogy, and gene duplication and loss (Edwards 2009; Galtier and Daubin 2008; Smith et al. 2015). Previous studies on *Tillandsia* subg. *Tillandsia* indeed found evidence for changes in population sizes and elevated rates of gene duplication and loss, specifically associated with photosynthetic syndrome shifts (de La Harpe et al. 2020; Groot Crego et al. 2024; Yardeni et al. 2022). We suggest there is strong evidence for ancestral hybridization as the source of discordance between *T. utriculata* and other Central American subclades: f-statistic and D-statistic tests are robust to the presence of ILS, as is PhyloNet (Malinsky et al. 2021; Martin et al. 2015; Wen et al. 2018) and ancestral population structure is unlikely to segregate through the demographic events that accompanied *Tillandsia*’s dispersal into Central America. Regardless, the generality of the obtained signal could be influenced by our limited sampling; a study employing a wider sampling could further infer if hybridization involved related or ancient lineages, while considering disparate molecular processes.

Genetic variation introduced through hybridization can be maladaptive, neutral or adaptive (Moran et al. 2021; Wong et al. 2022; Zeberg and Pääbo 2020). While its consequences remain difficult to predict and negative consequences may not be observable for long periods, an adaptive role of introgression has been demonstrated in numerous animal and plant taxa (Arnold et al. 2016; Dasmahapatra et al. 2012; Lamichhaney et al. 2015; Leroy et al. 2020; Taylor and Larson 2019). In this study, we did not directly test for the hypothesis that hybridization facilitated adaptation. However, we have identified two candidate regions on chromosome 18, with genomic signatures of hybridization occurring between species that share similar photosynthetic syndromes. Interestingly, the identified regions coincide with low gene density and one of them occurs at a chromosome edge. These findings echo the general expectation regarding the theory, posing that higher rates of hybridization correlate with regions of low gene density, since hybridized fragments in gene-rich regions are more likely to be detrimental (Barton and Bengtsson, 1986; Martin and Jiggins 2017; Sankararaman et al. 2014). The incomplete taxon sampling in our current study limits our ability to draw generalities about the extent of adaptive hybridization and introgression in *Tillandsia*: however, considering the rapid accumulation of morphological and physiological disparity we may hypothesize that hybridization in *Tillandsia* contributed to its success in a wide range of habitats, potentially introducing novel alleles through porous species boundaries and driving this evolutionary radiation. Further efforts will be needed to identify additional potential adaptive regions and uncover the key genes within.

We used whole-genome sequencing to deeply investigate the phylogenomics of a remarkable *Tillandsia* radiation. Recent research addressed questions on the genus’ evolutionary history, simultaneously expanding bromeliad and *Tillandsia* genomic resources and facilitating a range of evolutionary analyses. Ultimately, the complex history of *Tillandsia* remains elusive, calling for further investigations to uncover the interplay of processes that drove this rapid diversification. Future studies employing both a wider sampling and a deeper genomic coverage can characterise the genomic properties associated with diversification and explore the prevalence and consequences of hybridization across different evolutionary scales.

## Acknowledgements

This research was supported with funding from the Christian Lexer professorship start-up BE772002 at the University of Vienna, and in part by the Austrian Science Fund (FWF) [https://doi.org/10.55776/P35275] to O.P. The analyses took advantage of the Vienna Scientific Cluster (VSC). We thank Magnus Nordborg, James Pease and Hannes Svardal for insightful discussions and advice. We thank the members of Swiss SNSF Sinergia project CRSII3_147630 and Juan P. Pinzón for accession sampling.

## Data availability statement

Online-only appendices can be found in the Dryad data repository: DOI: https://datadryad.org/stash/share/HU9AtN96f1Gy36L8N_ja0L8XiuCEM327k-j21dwJUZY

Additional information, scripts and protocols are available from: https://github.com/giyany/TillandsiaPhylo/

All raw sequence reads will be deposited in NCBI-SRA (accession ID XXXX).

## Supporting files and tables

**Supporting file 1 –** maximum-likelihood trees constructed for each of the 25 chromosomes. Eat tree was inferred on a dataset of concatenated SNPs with IQ-TREE, using substitution model TVMe+R2 with ascertainment bias correction. Branch lengths were calculated by number of substitutions per site and branch support was assessed using ultra-fast bootstrap estimation with 1,000 replicates.

**Supporting file 2** - Heatmaps summarizing 7,141 four-taxon D-statistic tests for each of the 25 reference chromosomes, indicated on each figure. *Tillandsia complanata* was used as the outgroup in all tests. The four taxa in each test have been rearranged to always obtain positive D values, and P2 and P3 are shown on the axes. Colour indicates the value of D and log value of p-value, as appears in legend (bottom right).

**Supporting file 3** - Examination of the effects of different filtering thresholds for minor alleles on different parts of the analysis. Using the explicit value of MAC (minor allele count) with values between 0-5, we performed maximum likelihood tree inference, species tree inference with quartet support and genome-wide D-statistic calculation (see main methods).

**Supporting Table S1.** sampled accessions in this study with collection, locality and voucher information. samples marked grey were removed from analysis (see table 3)

**Supporting Table S2.** List of species sampled for this study with clade assignment, phenotype and ecology (if known). Sheet 2 details references to corresponding literature.

**Supporting Table S3.** Samples accessions with information about sequence read number, alignment rates and number of sequences after filtering. samples marked grey were removed from analysis.

**Supporting Table S4.** Description of data-sets used in main analysis in the study, their properties and use in analysis.

