## Supporting Fig for "The explosive radiation of the Neotropical *Tillandsia* subgenus *Tillandsia* (Bromeliaceae) has been accompanied by pervasive hybridization"

### Supporting Figures

**Supporting Figure S1.** Maximum-likelihood (ML) phylogenomic tree inferred on a supermatrix partitioned by genes with IQ-TREE2, using ModelFinder to select the best-fitting partitioning scheme and models. Branch lengths were calculated by number of substitutions per site and branch support was assessed using ultra-fast bootstrap estimation with 1,000 replicates. Colors follow the clades in Figure 1.

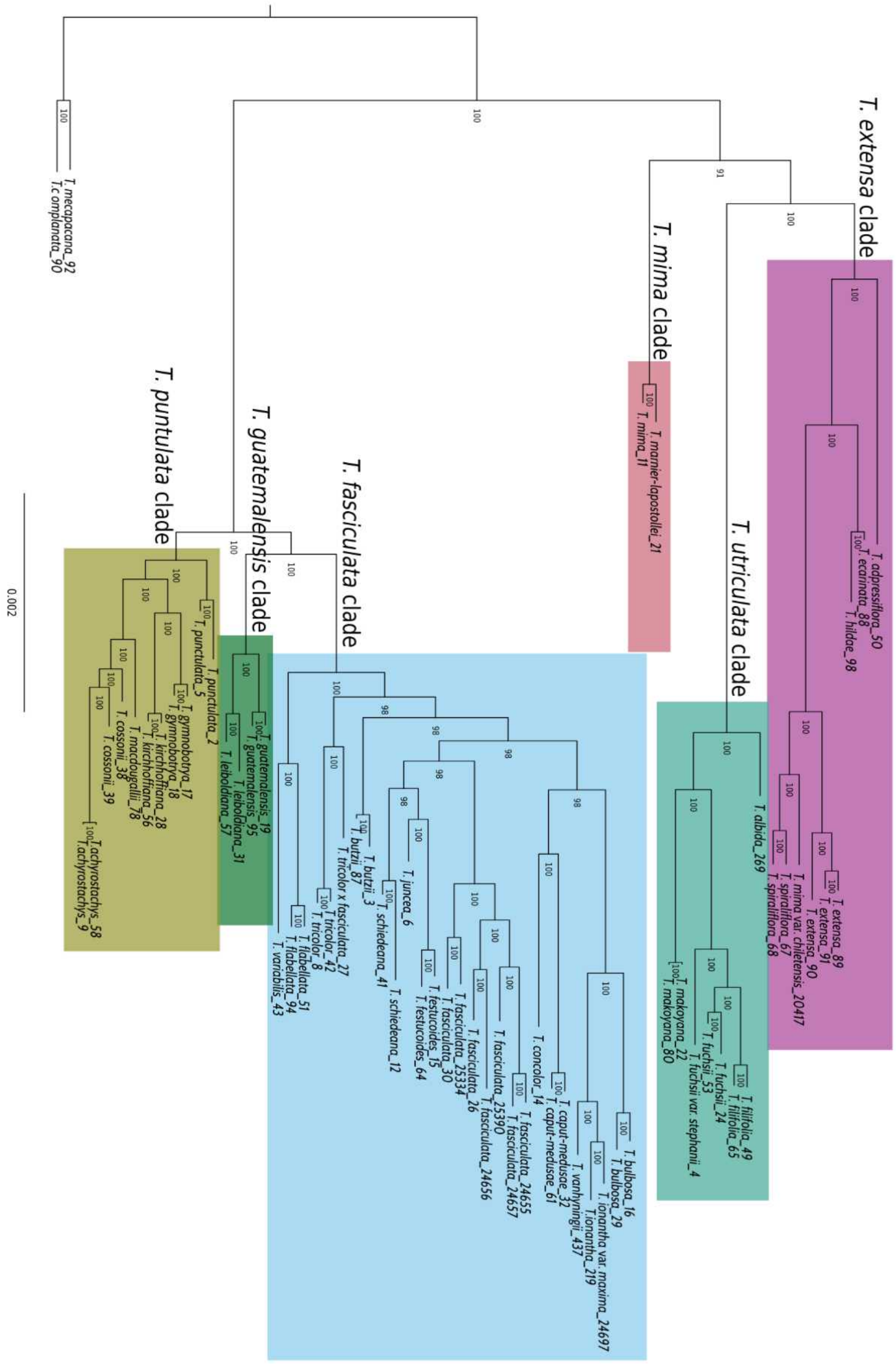

Pervasive hybridization in radiated *Tillandsia*

**Supporting Figure S2.** *Tillandsia zoquensis* monophyletic with *T. fasciculata*. A maximum-likelihood tree was inferred on a dataset of concatenated SNPs with IQ-TREE, using substitution model TVMe+R2 with ascertainment bias correction. For sample collection details, see Supporting Table S2.

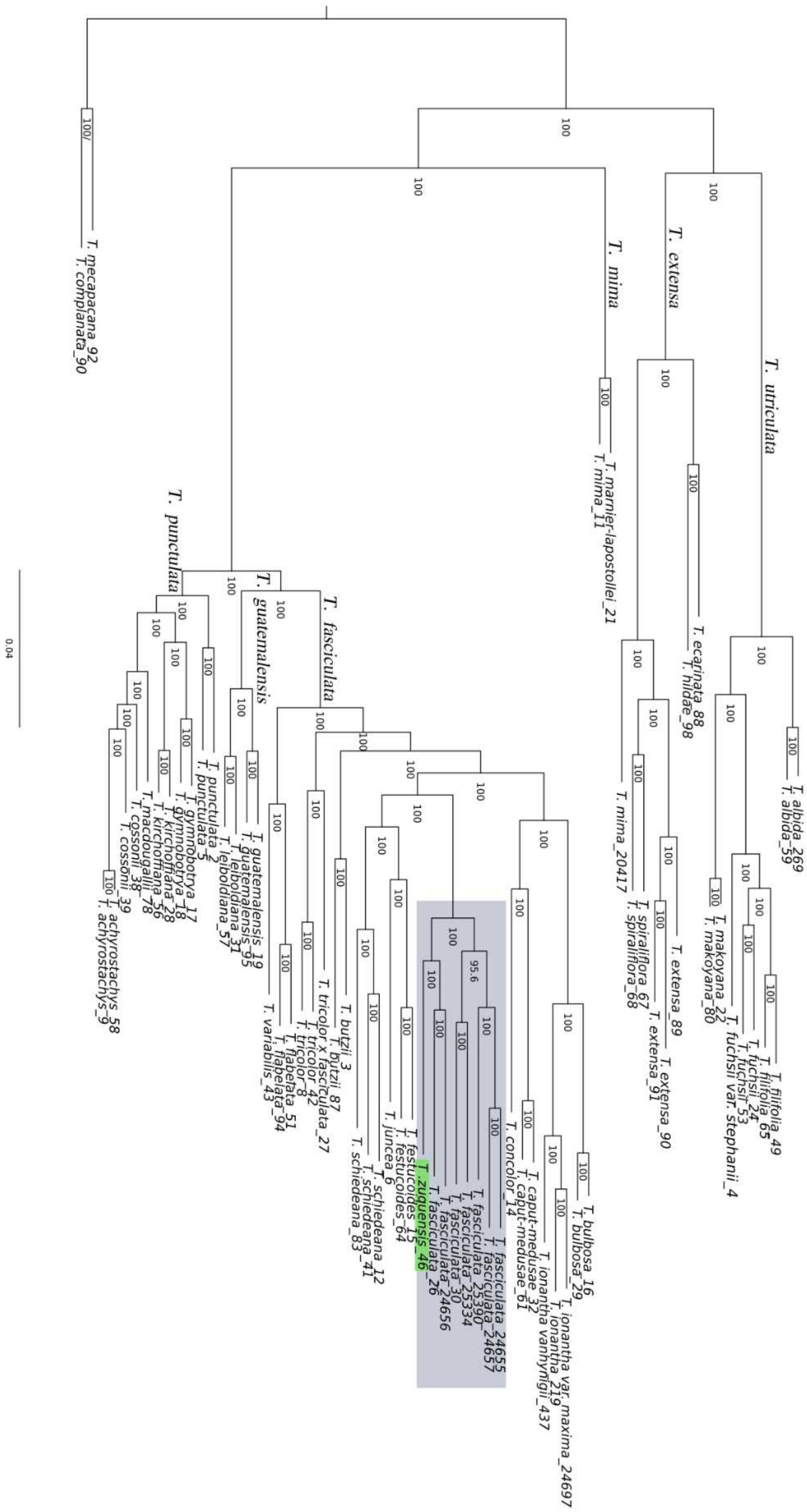

**Supporting Figure S3.** Visualisation of gene tree discordance. Coalescent-based species tree was generated on 15,791 genomic windows using ASTRAL-III. For each branch, the top number indicates the number of gene trees concordant with the species tree at that node and the bottom number indicates the number of gene trees in conflict with that species tree. Pie charts at the nodes show levels of gene tree discordance: the percentages of concordant gene trees (blue), the top alternative bipartition (green), other conflicting topologies (red) and uninformative gene trees (grey).

Pervasive hybridization in radiated *Tillandsia*

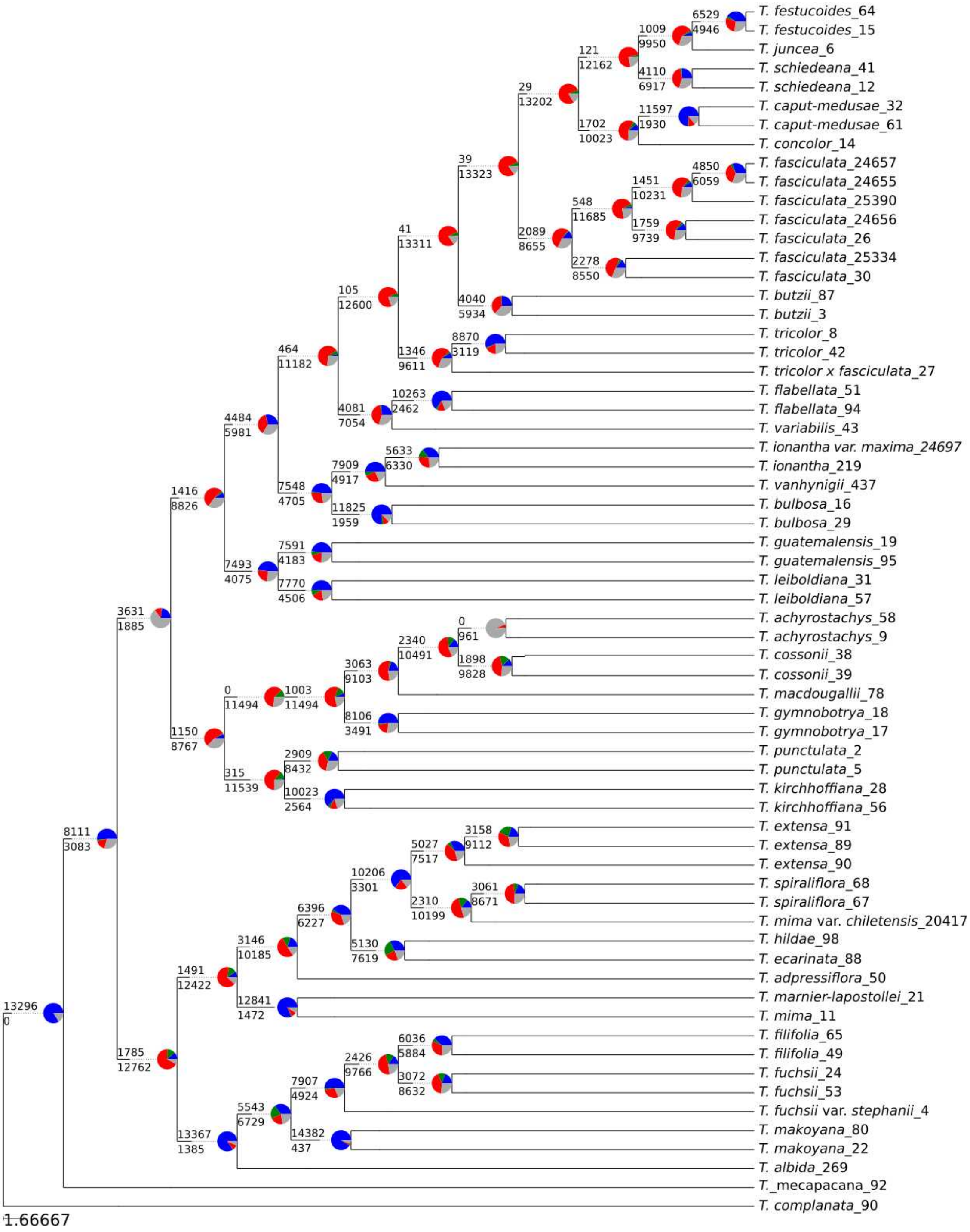

**Supporting Figure S4.** Widespread deviations from tree-like structure in *Tillandsia* phylogeny. A Cladogram of 100 randomly selected ML phylogenetic trees inferred from non-overlapping 10kb genomic windows. Branch lengths were forced into ultrametric structure for visualisation purposes only. Colours follow the legend and Figure 1.

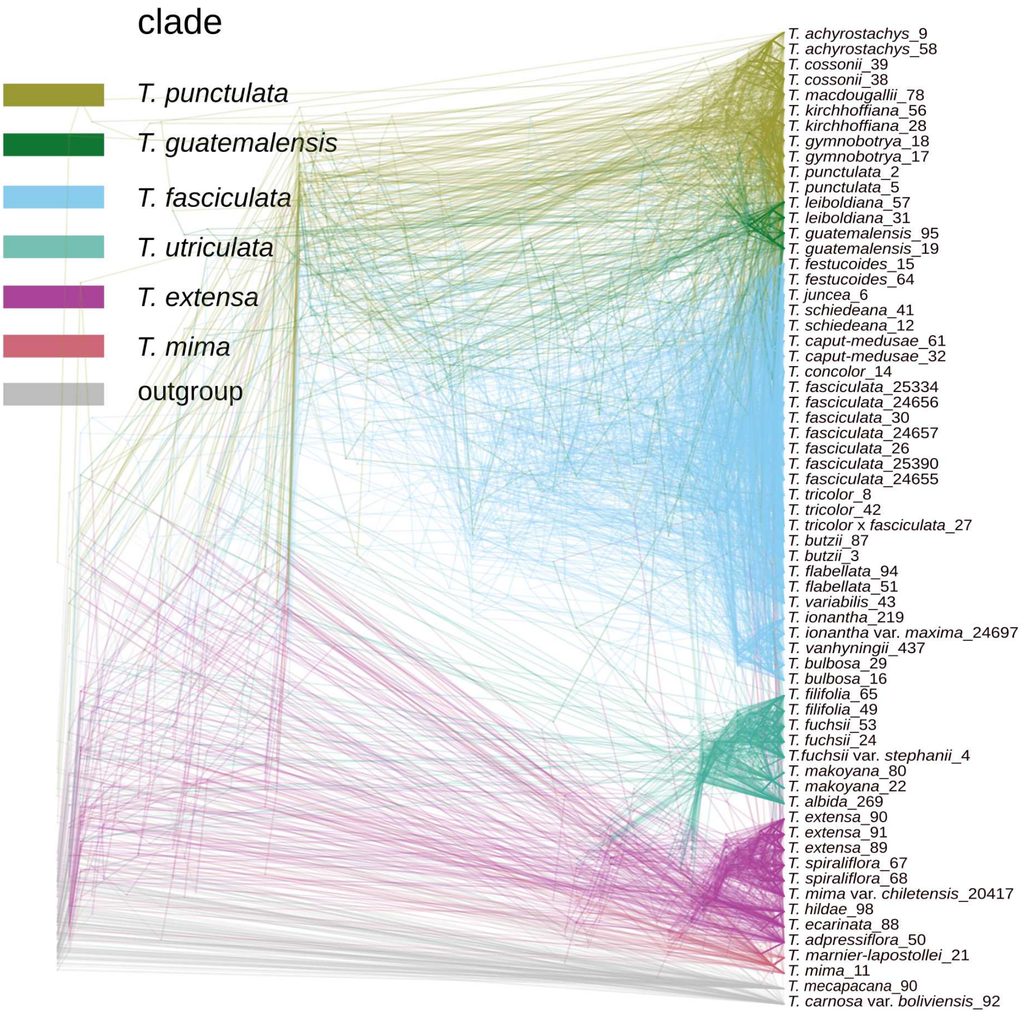

**Supporting Figure S5. a**, Derived alleles conveying deviations from tree structure in the Central American K clades. The Euler diagram represents the proportion of significantly elevated $D_{min}$ scores for comparisons in trios including species from within or between clades. Numbers indicate the total number of significantly elevated $D_{min}$ for each overlap. **b**, PCA analysis for all clades on bi-allelic, distance-pruned 16,204 SNPs, allowing maximum 10% missing data.

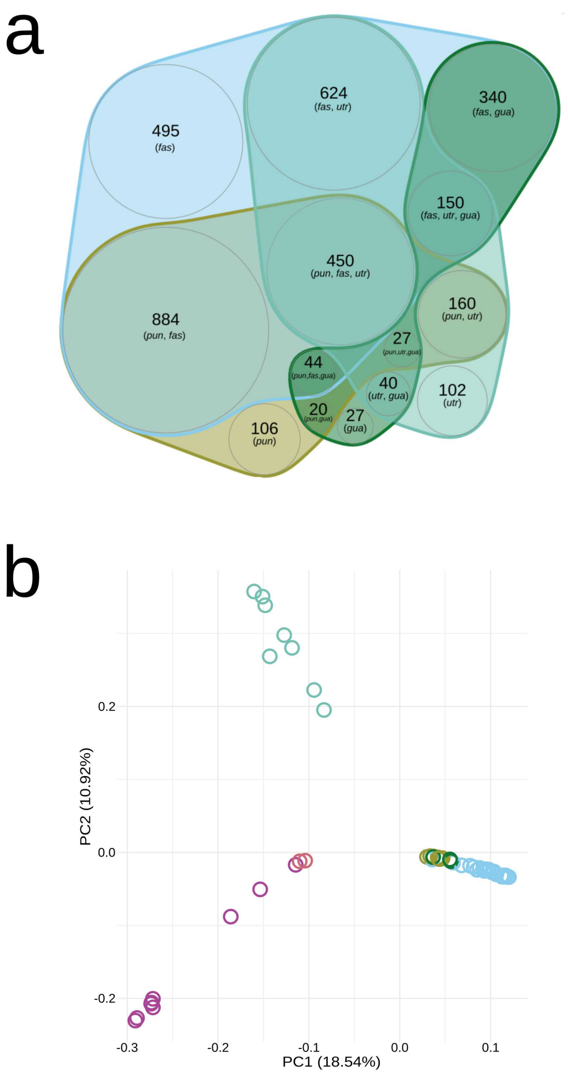

**Supporting Figure S6.** Variation along the genome in relationships between the major *Tillandsia* clades. Visualisation was produced using topology weighting by iterative sampling of subtrees in genomic windows of 50 SNPs. Colours represent the frequency of each topology along chromosomal position in each chromosome: white sectors indicate regions excluded due to high levels of missing data, including highly repetitive regions. Two chromosomes are presented: chromosome 4, to depict typical distribution of the main topologies, and chromosome 18, to contrast.

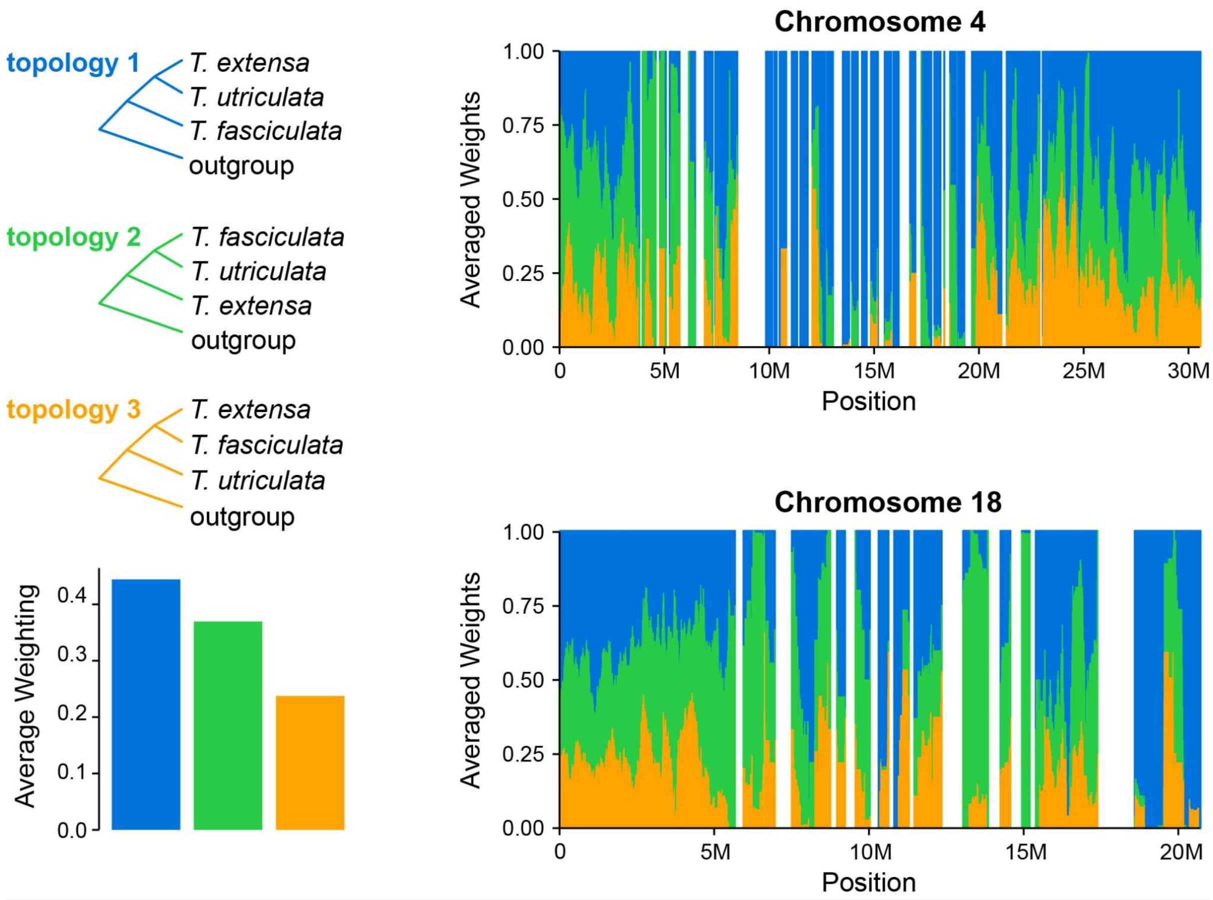

**Supporting Figure S7.** Heatmap summarizing 7,141 four-taxon D-statistic tests. *Tillandsia complanata* was used as the outgroup in all tests. The four taxa in each test have been rearranged to always obtain positive D values and P2 and P3 are shown on the axes. Colour indicates the value of D and the log value of p-value, the latter estimated using a block jackknife procedure with a 200 kb window size and corrected for family-wise error rate. Colours on the bars correspond to the clades in Figure 1.

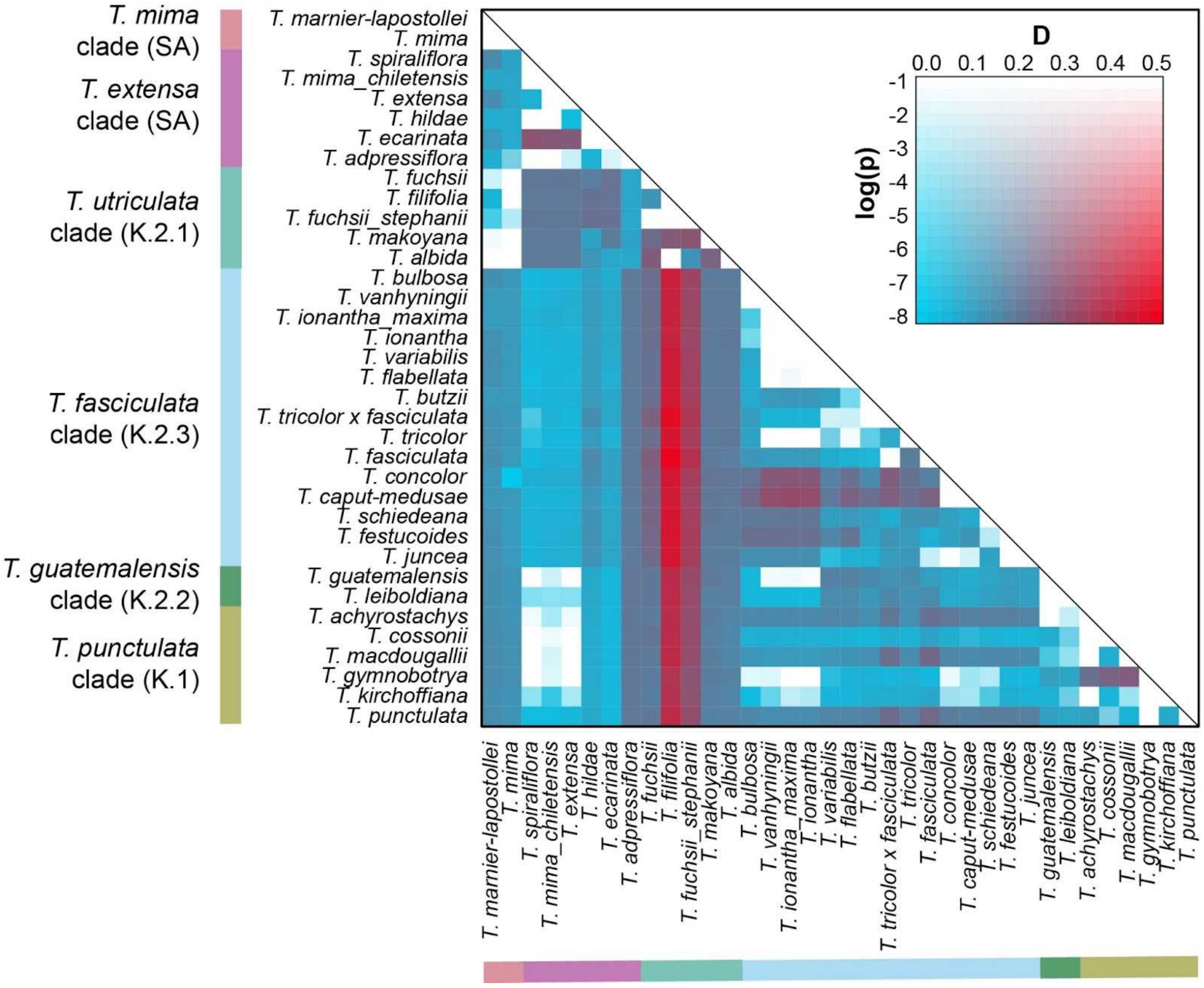

**Supporting Figure S8.** Heatmap summarising 7,141 *f*4-ratio tests. *Tillandsia complanata* was used as the outgroup for all tests. Color indicates the value of *f*4 and log value of p-value, the latter estimated using a block jackknife procedure with a 200 kb window size and corrected for family wise error rate. Colors correspond to the clades in Figure 1.

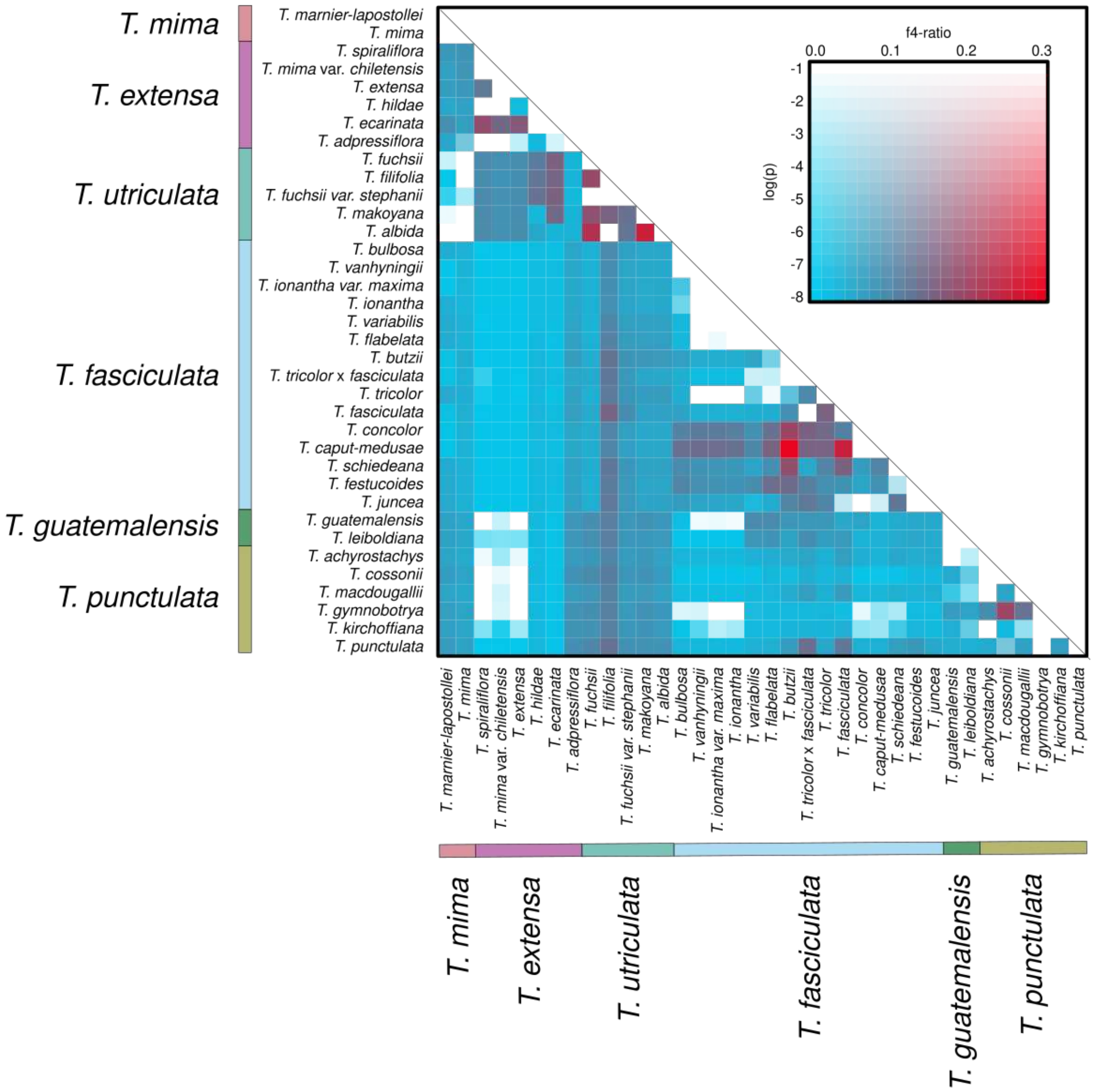
