## supporting file 1 for "The explosive radiation of the Neotropical *Tillandsia* subgenus *Tillandsia* (Bromeliaceae) has been accompanied by pervasive hybridization"

**Supporting file 1 –** maximum-likelihood trees constructred for each of the 25 chromosomes. Each tree was inferred on a dataset of concatenated SNPs with IQ-TREE, using substitution model TVMe+R2 with ascertainment bias correction. Branch lengths were calculated by number of substitutions per site and branch support was assessed using ultra-fast bootstrap estimation with 1,000 replicates.

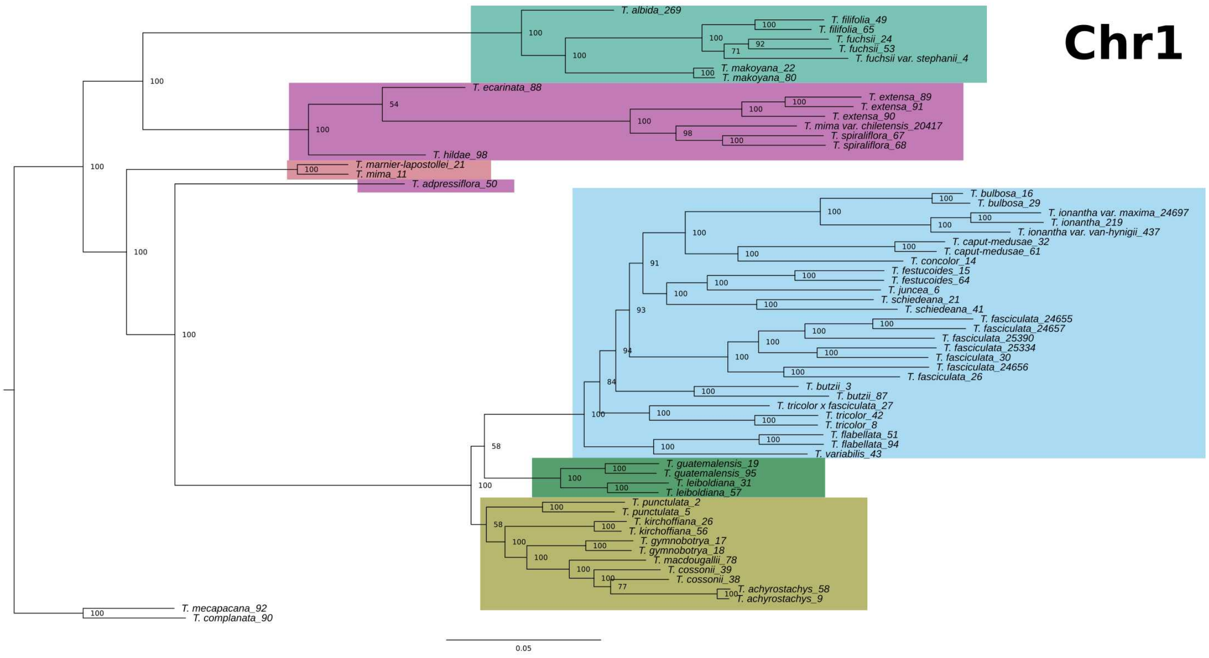
Chr1
T. albida_269
T. filifolia_49
T. filifolia_65
T. fuchsii_24
T. fuchsii_53
T. fuchsii var. stephanii_4
T. makoyana_22
T. makoyana_80
T. ecarinata_88
T. extensa_89
T. extensa_91
T. extensa_90
T. mima var. chiletensis_20417
T. spiraliflora_67
T. spiraliflora_68
T. hildae_98
T. marnier-lapostollei_21
T. mima_11
T. adpressiflora_50
T. bulbosa_16
T. bulbosa_29
T. ionantha var. maxima_24697
T. ionantha_219
T. ionantha var. van-hynigii_437
T. caput-medusae_32
T. caput-medusae_61
T. concolor_14
T. festucoides_15
T. festucoides_64
T. juncea_6
T. schiedeana_21
T. schiedeana_41
T. fasciculata_24655
T. fasciculata_24657
T. fasciculata_25390
T. fasciculata_25334
T. fasciculata_30
T. fasciculata_24656
T. fasciculata_26
T. butzii_3
T. butzii_87
T. tricolor x fasciculata_27
T. tricolor_42
T. tricolor_8
T. flabellata_51
T. flabellata_94
T. variabilis_43
T. guatemalensis_19
T. guatemalensis_95
T. leiboldiana_31
T. leiboldiana_57
T. punctulata_2
T. punctulata_5
T. kirchoffiana_26
T. kirchoffiana_56
T. gymnobotrya_17
T. gymnobotrya_18
T. macdougallii_78
T. cossonii_39
T. cossonii_38
T. achyrostachys_58
T. achyrostachys_9
T. mecapacana_92
T. complanata_90
0.05

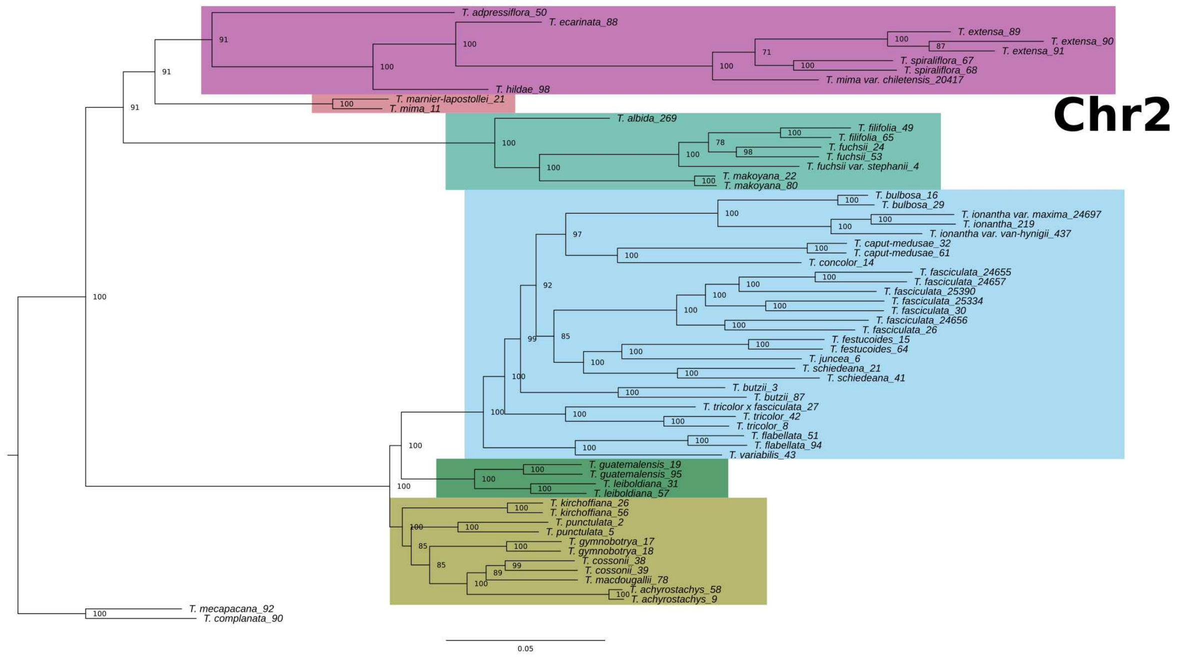
Chr2
T. adpressiflora_50
T. ecarinata_88
T. extensa_89
T. extensa_90
T. extensa_91
T. spiraliflora_67
T. spiraliflora_68
T. mima var. chiletensis_20417
T. hildae_98
T. marnier-lapostollei_21
T. mima_11
T. albida_269
T. filifolia_49
T. filifolia_65
T. fuchsii_24
T. fuchsii_53
T. fuchsii var. stephanii_4
T. makoyana_22
T. makoyana_80
T. bulbosa_16
T. bulbosa_29
T. ionantha var. maxima_24697
T. ionantha_219
T. ionantha var. van-hynigii_437
T. caput-medusae_32
T. caput-medusae_61
T. concolor_14
T. fasciculata_24655
T. fasciculata_24657
T. fasciculata_25390
T. fasciculata_25334
T. fasciculata_30
T. fasciculata_24656
T. fasciculata_26
T. festucoides_15
T. festucoides_64
T. juncea_6
T. schiedeana_21
T. schiedeana_41
T. butzii_3
T. butzii_87
T. tricolor x fasciculata_27
T. tricolor_42
T. tricolor_8
T. flabellata_51
T. flabellata_94
T. variabilis_43
T. guatemalensis_19
T. guatemalensis_95
T. leiboldiana_31
T. leiboldiana_57
T. kirchoffiana_26
T. kirchoffiana_56
T. punctulata_2
T. punctulata_5
T. gymnobotrya_17
T. gymnobotrya_18
T. cossonii_38
T. cossonii_39
T. macdougallii_78
T. achyrostachys_58
T. achyrostachys_9
T. mecapacana_92
T. complanata_90
0.05

Yardeni et al.,

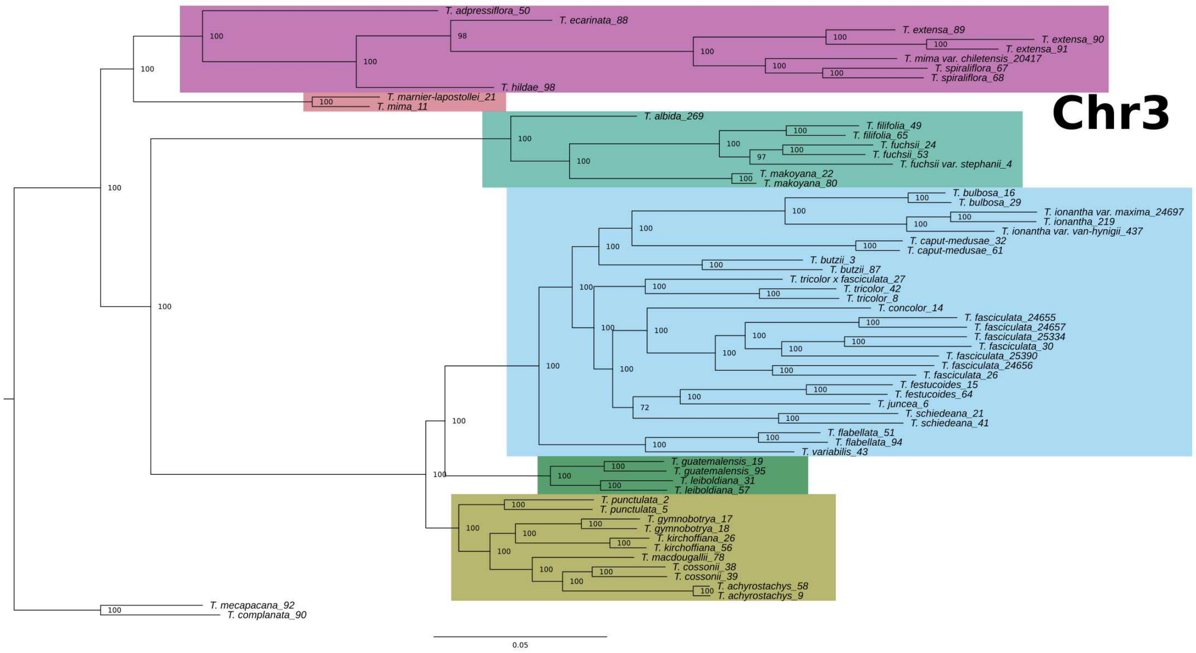


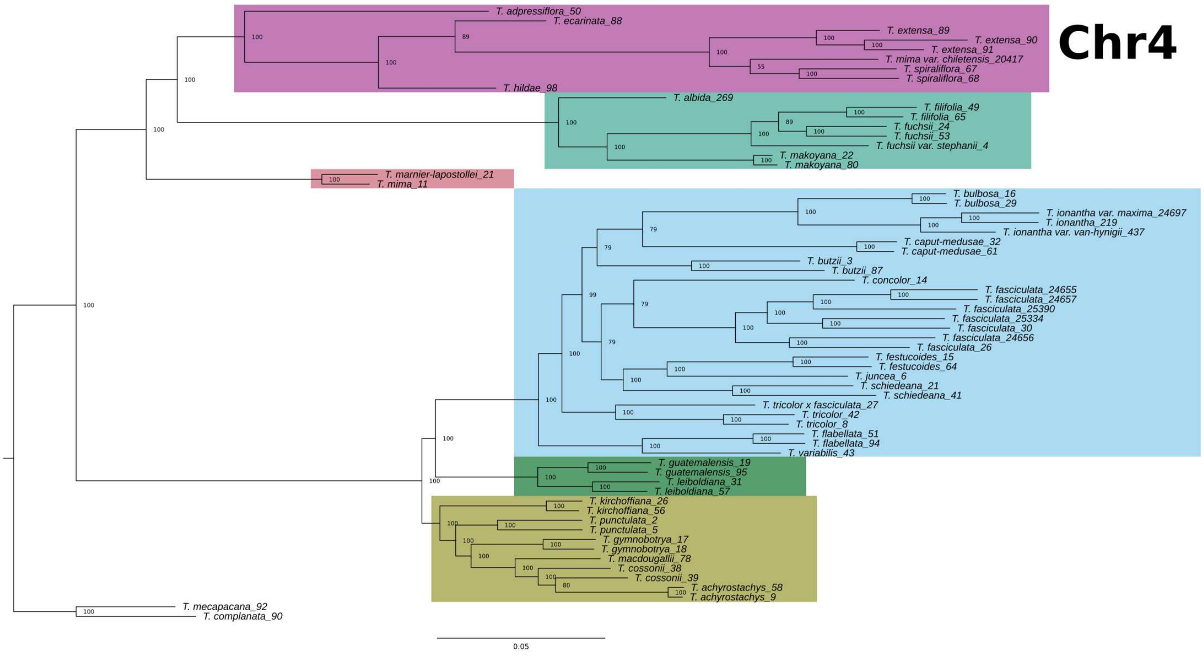

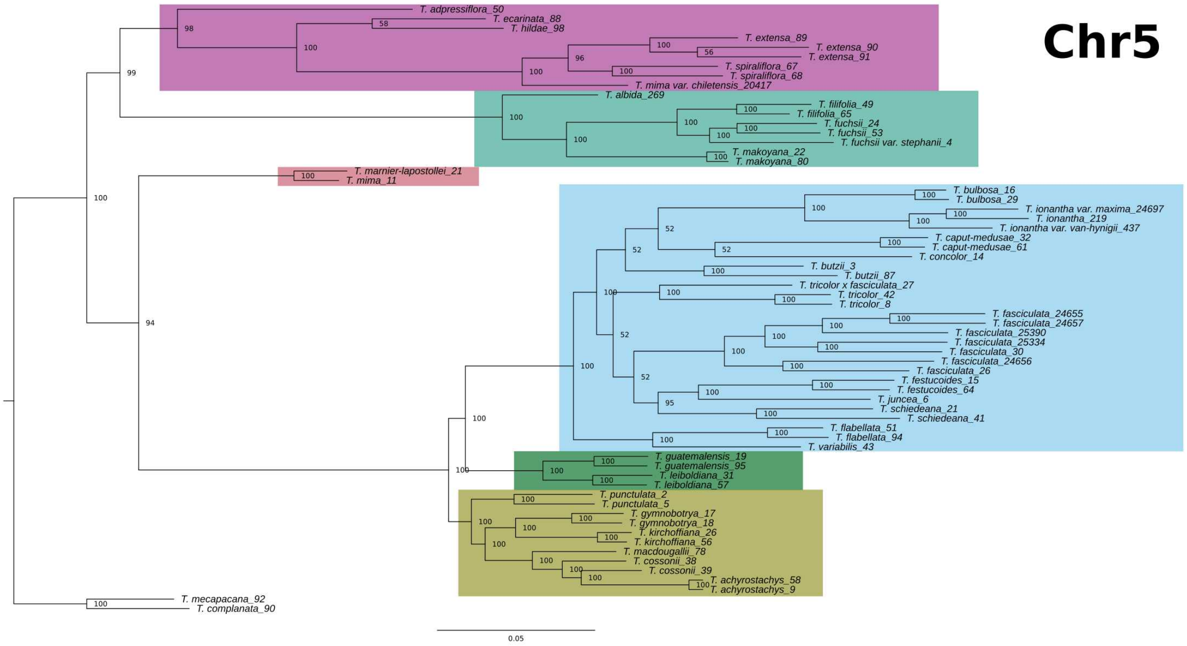
Chr5
T. adpressiflora_50
T. ecarinata_88
T. hildae_98
T. extensa_89
T. extensa_90
T. extensa_91
T. spiraliflora_67
T. spiraliflora_68
T. mima var. chiletensis_20417
T. albida_269
T. filifolia_49
T. filifolia_65
T. fuchsii_24
T. fuchsii_53
T. fuchsii var. stephanii_4
T. makoyana_22
T. makoyana_80
T. marnier-lapostollei_21
T. mima_11
T. bulbosa_16
T. bulbosa_29
T. ionantha var. maxima_24697
T. ionantha_219
T. ionantha var. van-hyngii_437
T. caput-medusae_32
T. caput-medusae_61
T. concolor_14
T. butzii_3
T. butzii_87
T. tricolor x fasciculata_27
T. tricolor_42
T. tricolor_8
T. fasciculata_24655
T. fasciculata_24657
T. fasciculata_25390
T. fasciculata_25334
T. fasciculata_30
T. fasciculata_24656
T. fasciculata_26
T. festucoides_15
T. festucoides_64
T. juncea_6
T. schiedeana_21
T. schiedeana_41
T. flabellata_51
T. flabellata_94
T. variabilis_43
T. guatemalensis_19
T. guatemalensis_95
T. leiboldiana_31
T. leiboldiana_57
T. punctulata_2
T. punctulata_5
T. gymnobotrya_17
T. gymnobotrya_18
T. kirchoffiana_26
T. kirchoffiana_56
T. macdougallii_78
T. cossonii_38
T. cossonii_39
T. achyrostachys_58
T. achyrostachys_9
T. mecapacana_92
T. complanata_90
0.05

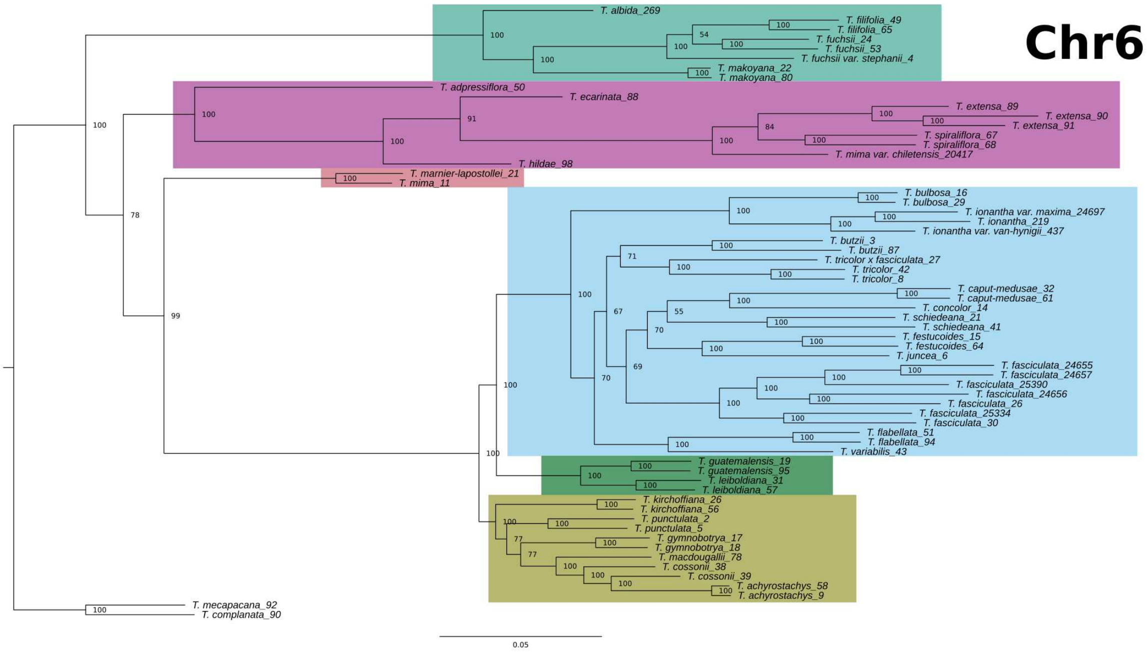
Chr6
T. albida_269
T. filifolia_49
T. filifolia_65
T. fuchsii_24
T. fuchsii_53
T. fuchsii var. stephanii_4
T. makoyana_22
T. makoyana_80
T. adpressiflora_50
T. ecarinata_88
T. extensa_89
T. extensa_90
T. extensa_91
T. spiraliflora_67
T. spiraliflora_68
T. mima var. chiletensis_20417
T. hildae_98
T. marnier-lapostollei_21
T. mima_11
T. bulbosa_16
T. bulbosa_29
T. ionantha var. maxima_24697
T. ionantha_219
T. ionantha var. van-hyngii_437
T. butzii_3
T. butzii_87
T. tricolor x fasciculata_27
T. tricolor_42
T. tricolor_8
T. caput-medusae_32
T. caput-medusae_61
T. concolor_14
T. schiedeana_21
T. schiedeana_41
T. festucoides_15
T. festucoides_64
T. juncea_6
T. fasciculata_24655
T. fasciculata_24657
T. fasciculata_25390
T. fasciculata_24656
T. fasciculata_26
T. fasciculata_25334
T. fasciculata_30
T. flabellata_51
T. flabellata_94
T. variabilis_43
T. guatemalensis_19
T. guatemalensis_95
T. leiboldiana_31
T. leiboldiana_57
T. kirchoffiana_26
T. kirchoffiana_56
T. punctulata_2
T. punctulata_5
T. gymnobotrya_17
T. gymnobotrya_18
T. macdougallii_78
T. cossonii_38
T. cossonii_39
T. achyrostachys_58
T. achyrostachys_9
T. mecapacana_92
T. complanata_90
0.05

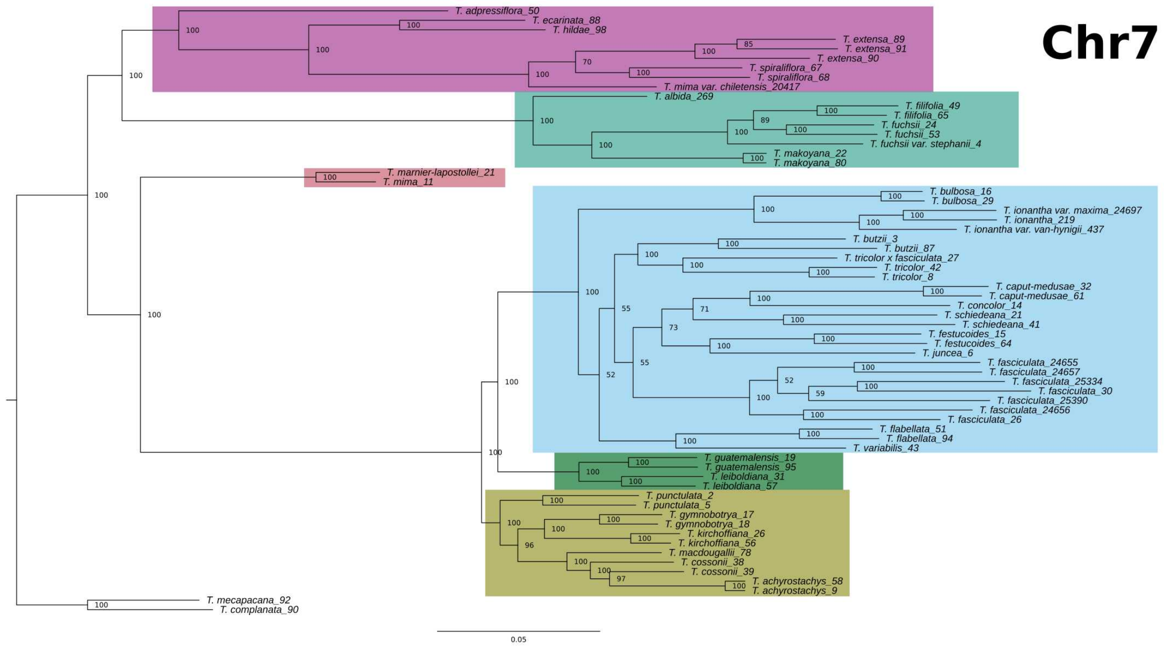
Chr7
T. adpressiflora_50
T. ecarinata_88
T. hildae_98
T. extensa_89
T. extensa_91
T. extensa_90
T. spiraliflora_67
T. spiraliflora_68
T. mima var. chiletensis_20417
T. albida_269
T. filifolia_49
T. filifolia_65
T. fuchsii_24
T. fuchsii_53
T. fuchsii var. stephanii_4
T. makoyana_22
T. makoyana_80
T. marnier-lapostollei_21
T. mima_11
T. bulbosa_16
T. bulbosa_29
T. ionantha var. maxima_24697
T. ionantha_219
T. ionantha var. van-hyningii_437
T. butzii_3
T. butzii_87
T. tricolor x fasciculata_27
T. tricolor_42
T. tricolor_8
T. caput-medusae_32
T. caput-medusae_61
T. concolor_14
T. schiedeana_21
T. schiedeana_41
T. festucoides_15
T. festucoides_64
T. juncea_6
T. fasciculata_24655
T. fasciculata_24657
T. fasciculata_25334
T. fasciculata_30
T. fasciculata_25390
T. fasciculata_24656
T. fasciculata_26
T. flabellata_51
T. flabellata_94
T. variabilis_43
T. guatemalensis_19
T. guatemalensis_95
T. leiboldiana_31
T. leiboldiana_57
T. punctulata_2
T. punctulata_5
T. gymnobotrya_17
T. gymnobotrya_18
T. kirchoffiana_26
T. kirchoffiana_56
T. macdougallii_78
T. cossonii_38
T. cossonii_39
T. achyrostachys_58
T. achyrostachys_9
T. mecapacana_92
T. complanata_90
0.05

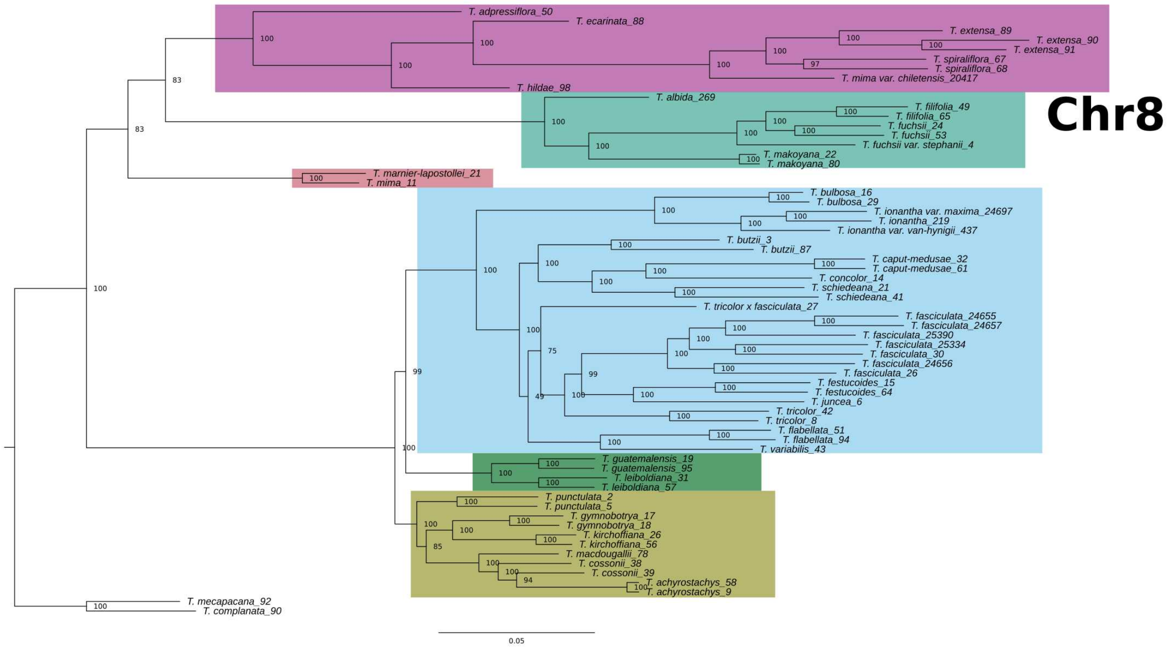
Chr8
T. adpressiflora_50
T. ecarinata_88
T. extensa_89
T. extensa_90
T. extensa_91
T. spiraliflora_67
T. spiraliflora_68
T. mima var. chiletensis_20417
T. hildae_98
T. albida_269
T. filifolia_49
T. filifolia_65
T. fuchsii_24
T. fuchsii_53
T. fuchsii var. stephanii_4
T. makoyana_22
T. makoyana_80
T. marnier-lapostollei_21
T. mima_11
T. bulbosa_16
T. bulbosa_29
T. ionantha var. maxima_24697
T. ionantha_219
T. ionantha var. van-hyningii_437
T. butzii_3
T. butzii_87
T. caput-medusae_32
T. caput-medusae_61
T. concolor_14
T. schiedeana_21
T. schiedeana_41
T. tricolor x fasciculata_27
T. fasciculata_24655
T. fasciculata_24657
T. fasciculata_25390
T. fasciculata_25334
T. fasciculata_30
T. fasciculata_24656
T. fasciculata_26
T. festucoides_15
T. festucoides_64
T. juncea_6
T. tricolor_42
T. tricolor_8
T. flabellata_51
T. flabellata_94
T. variabilis_43
T. guatemalensis_19
T. guatemalensis_95
T. leiboldiana_31
T. leiboldiana_57
T. punctulata_2
T. punctulata_5
T. gymnobotrya_17
T. gymnobotrya_18
T. kirchoffiana_26
T. kirchoffiana_56
T. macdougallii_78
T. cossonii_38
T. cossonii_39
T. achyrostachys_58
T. achyrostachys_9
T. mecapacana_92
T. complanata_90
0.05

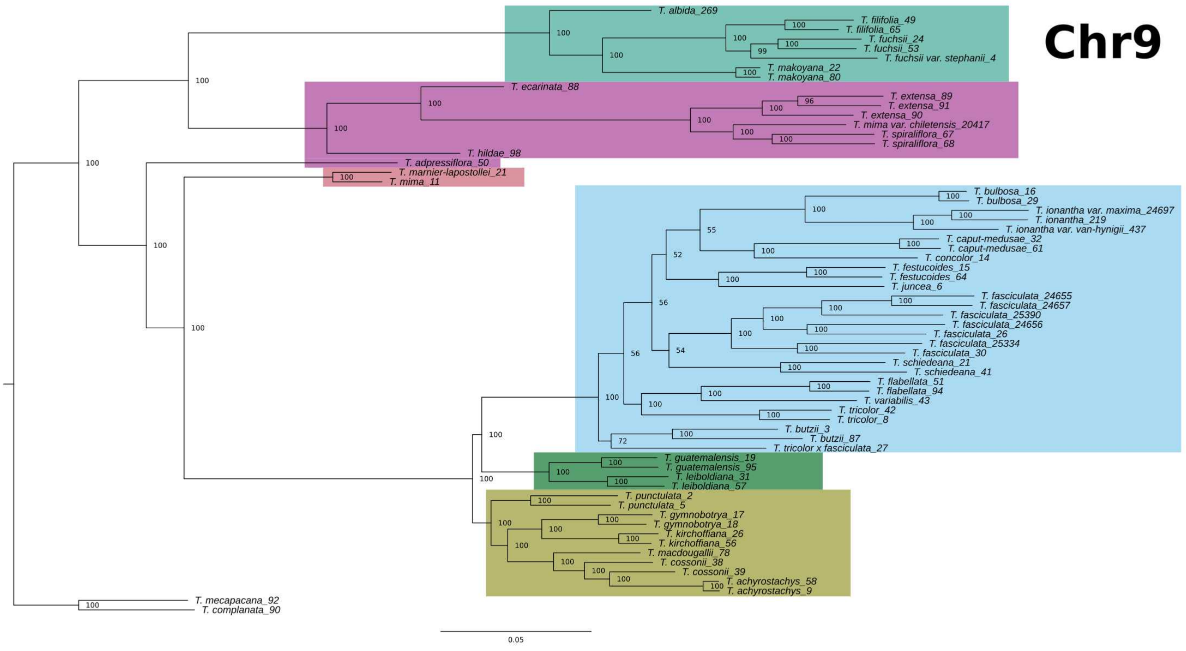
Chr9
T. albida_269
T. filifolia_49
T. filifolia_65
T. fuchsii_24
T. fuchsii_53
T. fuchsii var. stephanii_4
T. makoyana_22
T. makoyana_80
T. ecarinata_88
T. extensa_89
T. extensa_91
T. extensa_90
T. mima var. chiletensis_20417
T. spiraliflora_67
T. spiraliflora_68
T. hildae_98
T. adpressiflora_50
T. marnier-lapostollei_21
T. mima_11
T. bulbosa_16
T. bulbosa_29
T. ionantha var. maxima_24697
T. ionantha_219
T. ionantha var. van-hynigii_437
T. caput-medusae_32
T. caput-medusae_61
T. concolor_14
T. festucoides_15
T. festucoides_64
T. juncea_6
T. fasciculata_24655
T. fasciculata_24657
T. fasciculata_25390
T. fasciculata_24656
T. fasciculata_26
T. fasciculata_25334
T. fasciculata_30
T. schiedeana_21
T. schiedeana_41
T. flabellata_51
T. flabellata_94
T. variabilis_43
T. tricolor_42
T. tricolor_8
T. butzii_3
T. butzii_87
T. tricolor x fasciculata_27
T. guatemalensis_19
T. guatemalensis_95
T. leiboldiana_31
T. leiboldiana_57
T. punctulata_2
T. punctulata_5
T. gymnobotrya_17
T. gymnobotrya_18
T. kirchoffiana_26
T. kirchoffiana_56
T. macdougallii_78
T. cossonii_38
T. cossonii_39
T. achyrostachys_58
T. achyrostachys_9
T. mecapacana_92
T. complanata_90
0.05

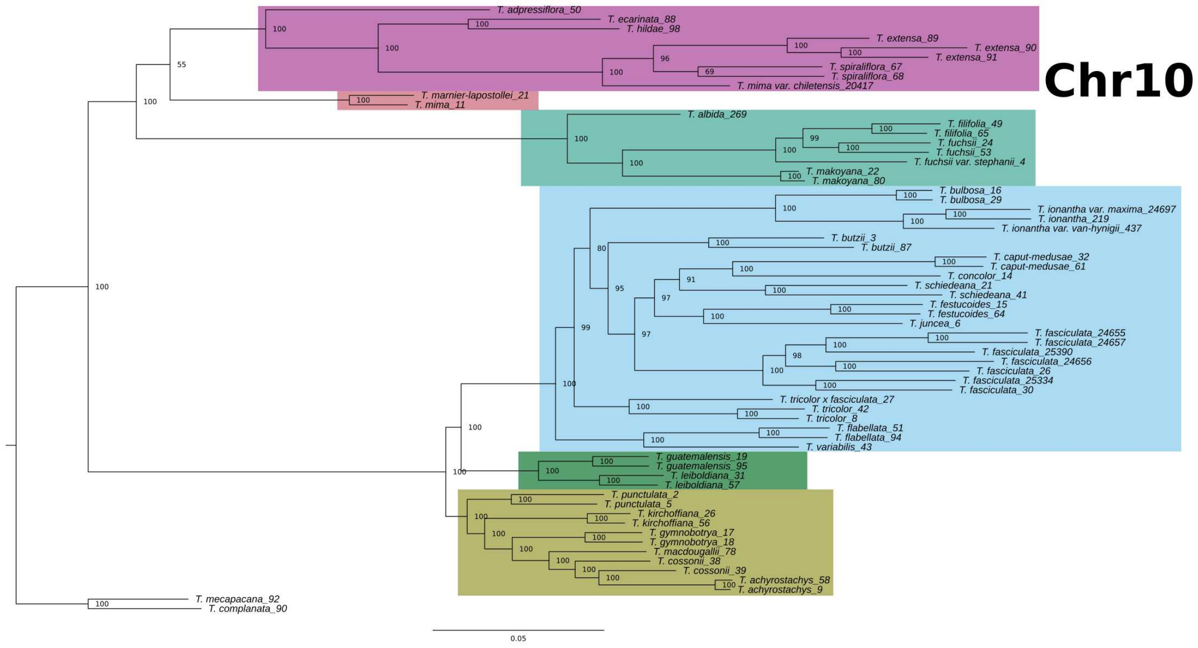
Chr10
T. adpressiflora_50
T. ecarinata_88
T. hildae_98
T. extensa_89
T. extensa_90
T. extensa_91
T. spiraliflora_67
T. spiraliflora_68
T. mima var. chiletensis_20417
T. marnier-lapostollei_21
T. mima_11
T. albida_269
T. filifolia_49
T. filifolia_65
T. fuchsii_24
T. fuchsii_53
T. fuchsii var. stephanii_4
T. makoyana_22
T. makoyana_80
T. bulbosa_16
T. bulbosa_29
T. ionantha var. maxima_24697
T. ionantha_219
T. ionantha var. van-hynigii_437
T. butzii_3
T. butzii_87
T. caput-medusae_32
T. caput-medusae_61
T. concolor_14
T. schiedeana_21
T. schiedeana_41
T. festucoides_15
T. festucoides_64
T. juncea_6
T. fasciculata_24655
T. fasciculata_24657
T. fasciculata_25390
T. fasciculata_24656
T. fasciculata_26
T. fasciculata_25334
T. fasciculata_30
T. tricolor x fasciculata_27
T. tricolor_42
T. tricolor_8
T. flabellata_51
T. flabellata_94
T. variabilis_43
T. guatemalensis_19
T. guatemalensis_95
T. leiboldiana_31
T. leiboldiana_57
T. punctulata_2
T. punctulata_5
T. kirchoffiana_26
T. kirchoffiana_56
T. gymnobotrya_17
T. gymnobotrya_18
T. macdougallii_78
T. cossonii_38
T. cossonii_39
T. achyrostachys_58
T. achyrostachys_9
T. mecapacana_92
T. complanata_90
0.05

Yardeni et al.,

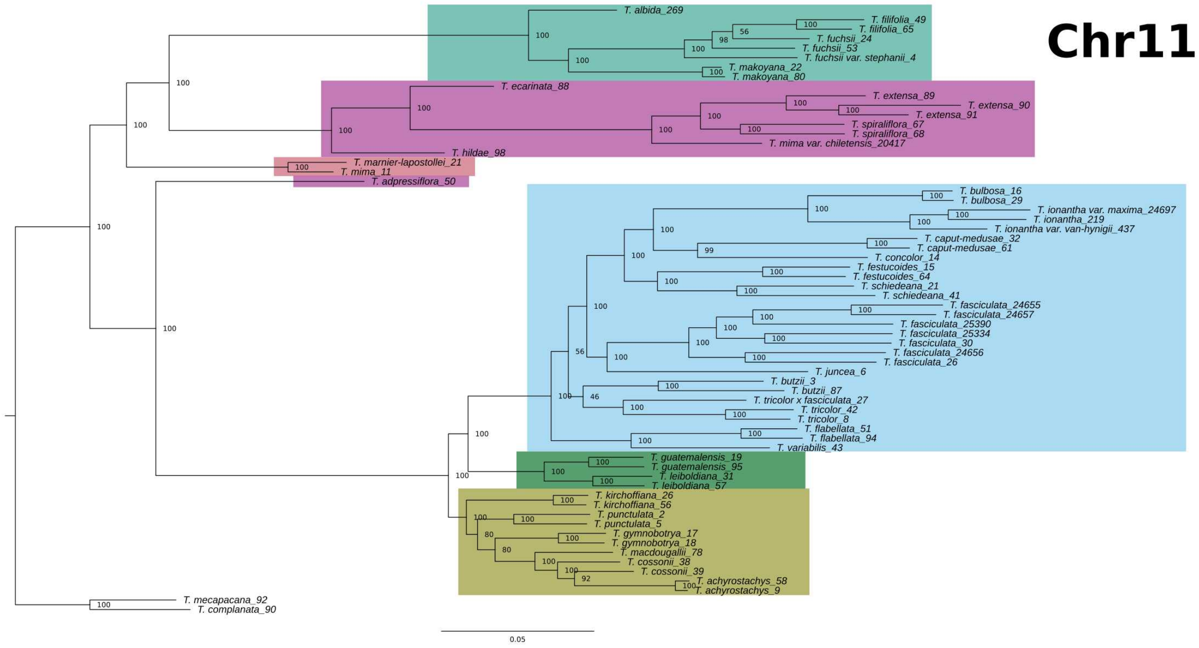


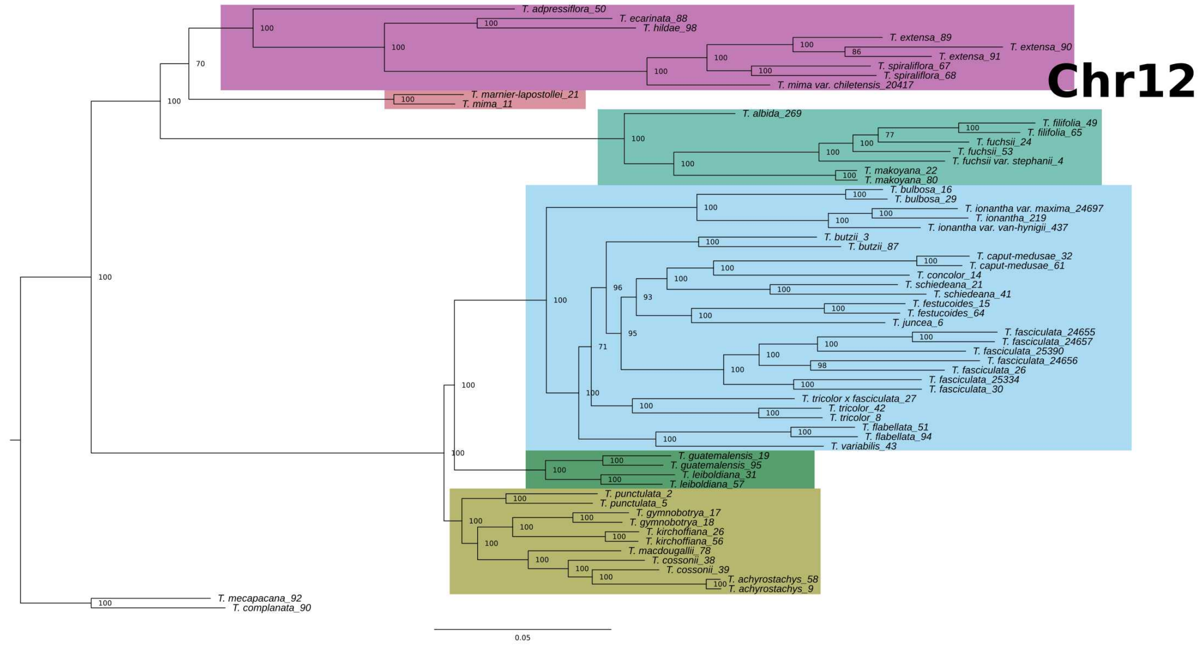

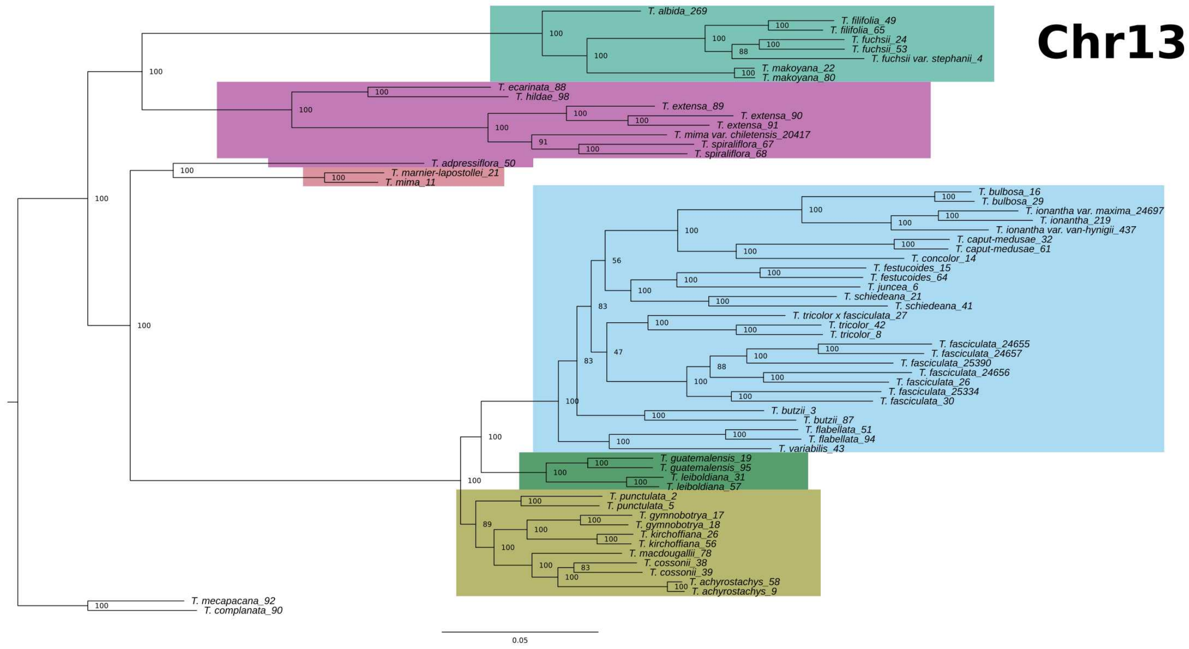
Chr13
T. albida_269
T. filifolia_49
T. filifolia_65
T. fuchsii_24
T. fuchsii_53
T. fuchsii var. stephanii_4
T. makoyana_22
T. makoyana_80
T. ecarinata_88
T. hildae_98
T. extensa_89
T. extensa_90
T. extensa_91
T. mima var. chiletensis_20417
T. spiraliflora_67
T. spiraliflora_68
T. adpressiflora_50
T. marnier-lapostollei_21
T. mima_11
T. bulbosa_16
T. bulbosa_29
T. ionantha var. maxima_24697
T. ionantha_219
T. ionantha var. van-hynigii_437
T. caput-medusae_32
T. caput-medusae_61
T. concolor_14
T. festucoides_15
T. festucoides_64
T. juncea_6
T. schiedeana_21
T. schiedeana_41
T. tricolor x fasciculata_27
T. tricolor_42
T. tricolor_8
T. fasciculata_24655
T. fasciculata_24657
T. fasciculata_25390
T. fasciculata_24656
T. fasciculata_26
T. fasciculata_25334
T. fasciculata_30
T. butzii_3
T. butzii_87
T. flabellata_51
T. flabellata_94
T. variabilis_43
T. guatemalensis_19
T. guatemalensis_95
T. leiboldiana_31
T. leiboldiana_57
T. punctulata_2
T. punctulata_5
T. gymnobotrya_17
T. gymnobotrya_18
T. kirchoffiana_26
T. kirchoffiana_56
T. macdougallii_78
T. cossonii_38
T. cossonii_39
T. achyrostachys_58
T. achyrostachys_9
T. mecapacana_92
T. complanata_90
0.05

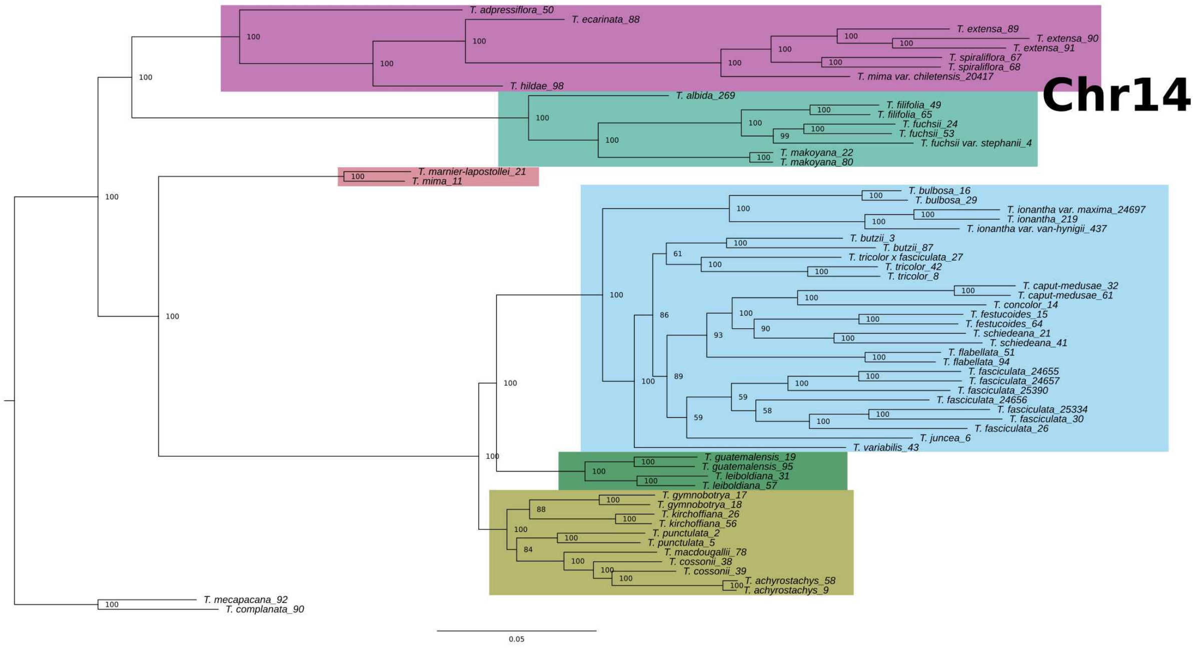
Chr14
T. adpressiflora_50
T. ecarinata_88
T. extensa_89
T. extensa_90
T. extensa_91
T. spiraliflora_67
T. spiraliflora_68
T. mima var. chiletensis_20417
T. hildae_98
T. albida_269
T. filifolia_49
T. filifolia_65
T. fuchsii_24
T. fuchsii_53
T. fuchsii var. stephanii_4
T. makoyana_22
T. makoyana_80
T. marnier-lapostollei_21
T. mima_11
T. bulbosa_16
T. bulbosa_29
T. ionantha var. maxima_24697
T. ionantha_219
T. ionantha var. van-hynigii_437
T. butzii_3
T. butzii_87
T. tricolor x fasciculata_27
T. tricolor_42
T. tricolor_8
T. caput-medusae_32
T. caput-medusae_61
T. concolor_14
T. festucoides_15
T. festucoides_64
T. schiedeana_21
T. schiedeana_41
T. flabellata_51
T. flabellata_94
T. fasciculata_24655
T. fasciculata_24657
T. fasciculata_25390
T. fasciculata_24656
T. fasciculata_25334
T. fasciculata_30
T. fasciculata_26
T. juncea_6
T. variabilis_43
T. guatemalensis_19
T. guatemalensis_95
T. leiboldiana_31
T. leiboldiana_57
T. gymnobotrya_17
T. gymnobotrya_18
T. kirchoffiana_26
T. kirchoffiana_56
T. punctulata_2
T. punctulata_5
T. macdougallii_78
T. cossonii_38
T. cossonii_39
T. achyrostachys_58
T. achyrostachys_9
T. mecapacana_92
T. complanata_90
0.05

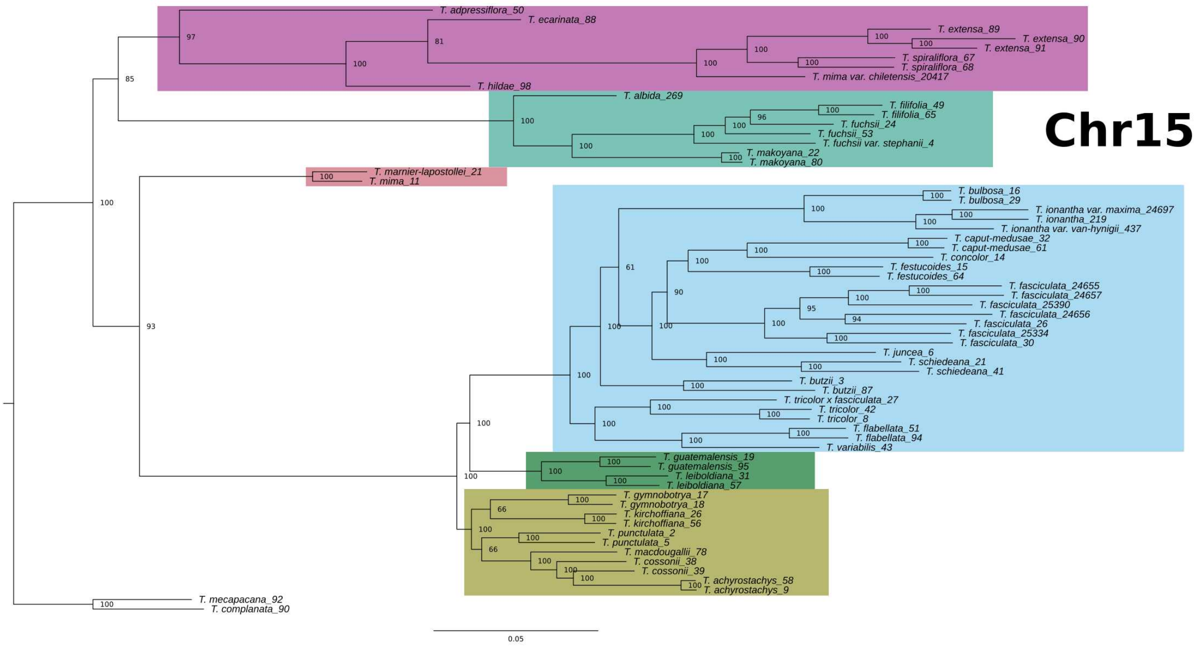
Chr15
T. adpressiflora_50
T. ecarinata_88
T. extensa_89
T. extensa_90
T. extensa_91
T. spiraliflora_67
T. spiraliflora_68
T. mima var. chiletensis_20417
T. hildae_98
T. albida_269
T. filifolia_49
T. filifolia_65
T. fuchsii_24
T. fuchsii_53
T. fuchsii var. stephanii_4
T. makoyana_22
T. makoyana_80
T. marnier-lapostollei_21
T. mima_11
T. bulbosa_16
T. bulbosa_29
T. ionantha var. maxima_24697
T. ionantha_219
T. ionantha var. van-hynigii_437
T. caput-medusae_32
T. caput-medusae_61
T. concolor_14
T. festucoides_15
T. festucoides_64
T. fasciculata_24655
T. fasciculata_24657
T. fasciculata_25390
T. fasciculata_24656
T. fasciculata_26
T. fasciculata_25334
T. fasciculata_30
T. juncea_6
T. schiedeana_21
T. schiedeana_41
T. butzii_3
T. butzii_87
T. tricolor x fasciculata_27
T. tricolor_42
T. tricolor_8
T. flabellata_51
T. flabellata_94
T. variabilis_43
T. guatemalensis_19
T. guatemalensis_95
T. leiboldiana_31
T. leiboldiana_57
T. gymnobotrya_17
T. gymnobotrya_18
T. kirchoffiana_26
T. kirchoffiana_56
T. punctulata_2
T. punctulata_5
T. macdougallii_78
T. cossonii_38
T. cossonii_39
T. achyrostachys_58
T. achyrostachys_9
T. mecapacana_92
T. complanata_90
0.05

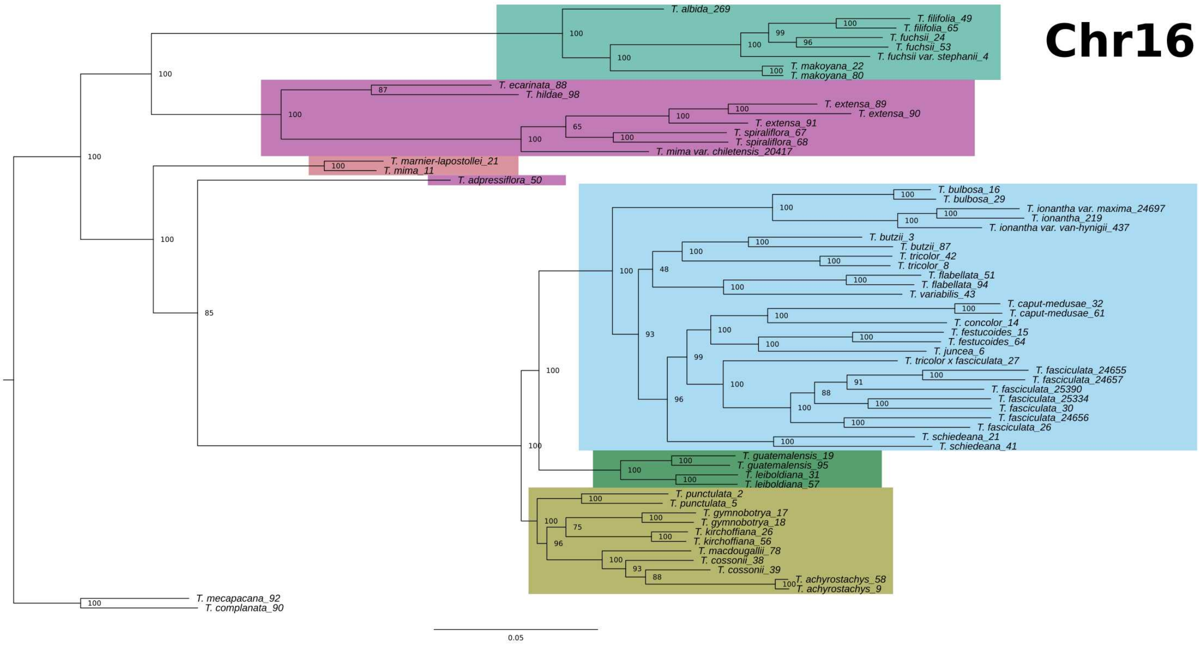
Chr16
T. albida_269
T. filifolia_49
T. filifolia_65
T. fuchsii_24
T. fuchsii_53
T. fuchsii var. stephanii_4
T. makoyana_22
T. makoyana_80
T. ecarinata_88
T. hildae_98
T. extensa_89
T. extensa_90
T. extensa_91
T. spiraliflora_67
T. spiraliflora_68
T. mima var. chiletensis_20417
T. marnier-lapostollei_21
T. mima_11
T. adpressiflora_50
T. bulbosa_16
T. bulbosa_29
T. ionantha var. maxima_24697
T. ionantha_219
T. ionantha var. van-hynigii_437
T. butzii_3
T. butzii_87
T. tricolor_42
T. tricolor_8
T. flabellata_51
T. flabellata_94
T. variabilis_43
T. caput-medusae_32
T. caput-medusae_61
T. concolor_14
T. festucoides_15
T. festucoides_64
T. juncea_6
T. tricolor x fasciculata_27
T. fasciculata_24655
T. fasciculata_24657
T. fasciculata_25390
T. fasciculata_25334
T. fasciculata_30
T. fasciculata_24656
T. fasciculata_26
T. schiedeana_21
T. schiedeana_41
T. guatemalensis_19
T. guatemalensis_95
T. leiboldiana_31
T. leiboldiana_57
T. punctulata_2
T. punctulata_5
T. gymnobotrya_17
T. gymnobotrya_18
T. kirchoffiana_26
T. kirchoffiana_56
T. macdougallii_78
T. cossonii_38
T. cossonii_39
T. achyrostachys_58
T. achyrostachys_9
T. mecapacana_92
T. complanata_90
0.05

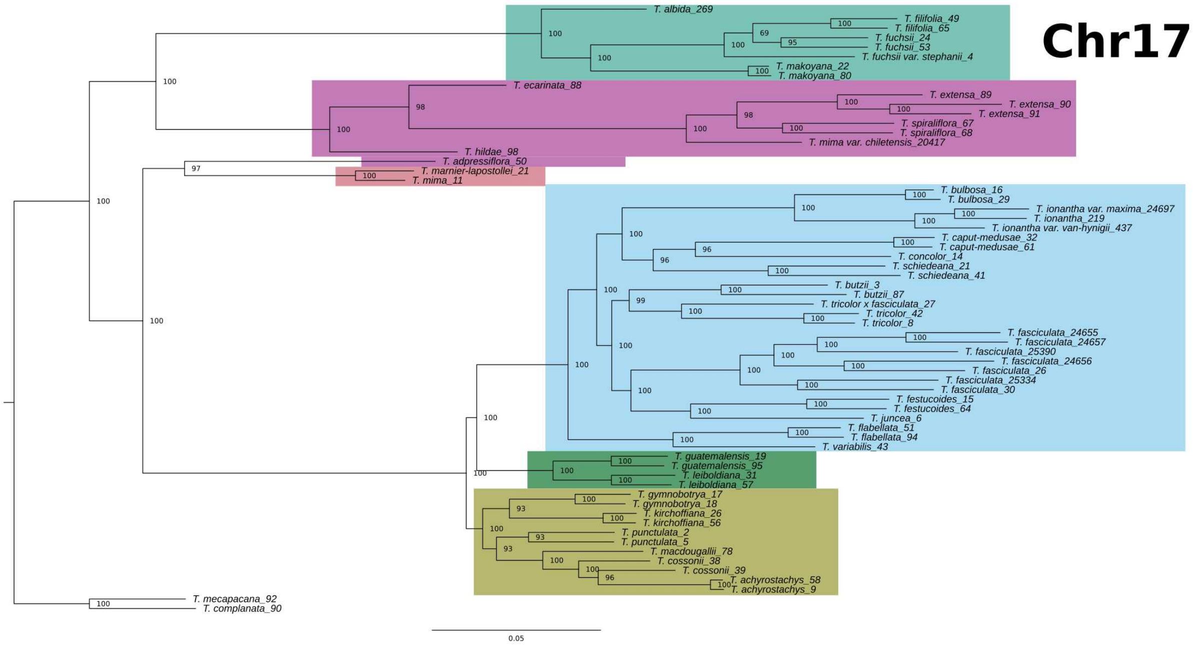
Chr17
T. albida_269
T. filifolia_49
T. filifolia_65
T. fuchsii_24
T. fuchsii_53
T. fuchsii var. stephanii_4
T. makoyana_22
T. makoyana_80
T. ecarinata_88
T. extensa_89
T. extensa_90
T. extensa_91
T. spiraliflora_67
T. spiraliflora_68
T. mima var. chiletensis_20417
T. hildae_98
T. adpressiflora_50
T. marnier-lapostollei_21
T. mima_11
T. bulbosa_16
T. bulbosa_29
T. ionantha var. maxima_24697
T. ionantha_219
T. ionantha var. van-hynigii_437
T. caput-medusae_32
T. caput-medusae_61
T. concolor_14
T. schiedeana_21
T. schiedeana_41
T. butzii_3
T. butzii_87
T. tricolor x fasciculata_27
T. tricolor_42
T. tricolor_8
T. fasciculata_24655
T. fasciculata_24657
T. fasciculata_25390
T. fasciculata_24656
T. fasciculata_26
T. fasciculata_25334
T. fasciculata_30
T. festucoides_15
T. festucoides_64
T. juncea_6
T. flabellata_51
T. flabellata_94
T. variabilis_43
T. guatemalensis_19
T. guatemalensis_95
T. leiboldiana_31
T. leiboldiana_57
T. gymnobotrya_17
T. gymnobotrya_18
T. kirchoffiana_26
T. kirchoffiana_56
T. punctulata_2
T. punctulata_5
T. macdougallii_78
T. cossonii_38
T. cossonii_39
T. achyrostachys_58
T. achyrostachys_9
T. mecapacana_92
T. complanata_90
0.05

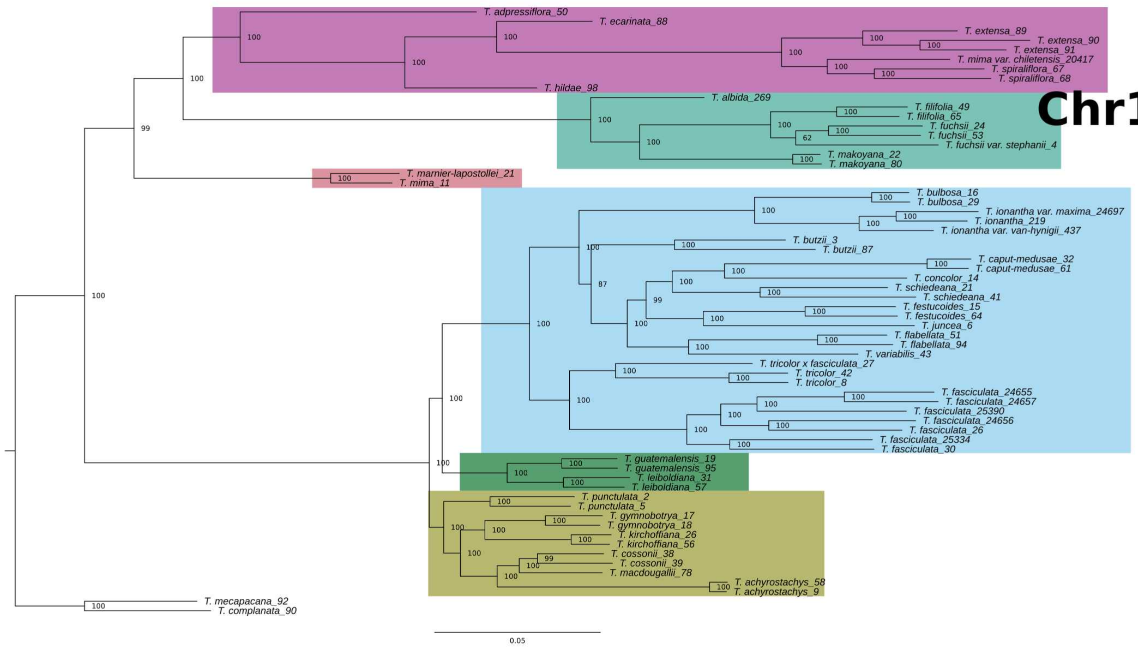
Chr18
T. adpressiflora_50
T. ecarinata_88
T. extensa_89
T. extensa_90
T. extensa_91
T. mima var. chiletensis_20417
T. spiraliflora_67
T. spiraliflora_68
T. hildae_98
T. albida_269
T. filifolia_49
T. filifolia_65
T. fuchsii_24
T. fuchsii_53
T. fuchsii var. stephanii_4
T. makoyana_22
T. makoyana_80
T. marnier-lapostollei_21
T. mima_11
T. bulbosa_16
T. bulbosa_29
T. ionantha var. maxima_24697
T. ionantha_219
T. ionantha var. van-hynigii_437
T. butzii_3
T. butzii_87
T. caput-medusae_32
T. caput-medusae_61
T. concolor_14
T. schiedeana_21
T. schiedeana_41
T. festucoides_15
T. festucoides_64
T. juncea_6
T. flabellata_51
T. flabellata_94
T. variabilis_43
T. tricolor x fasciculata_27
T. tricolor_42
T. tricolor_8
T. fasciculata_24655
T. fasciculata_24657
T. fasciculata_25390
T. fasciculata_24656
T. fasciculata_26
T. fasciculata_25334
T. fasciculata_30
T. guatemalensis_19
T. guatemalensis_95
T. leiboldiana_31
T. leiboldiana_57
T. punctulata_2
T. punctulata_5
T. gymnobotrya_17
T. gymnobotrya_18
T. kirchoffiana_26
T. kirchoffiana_56
T. cossonii_38
T. cossonii_39
T. macdougallii_78
T. achyrostachys_58
T. achyrostachys_9
T. mecapacana_92
T. complanata_90
0.05

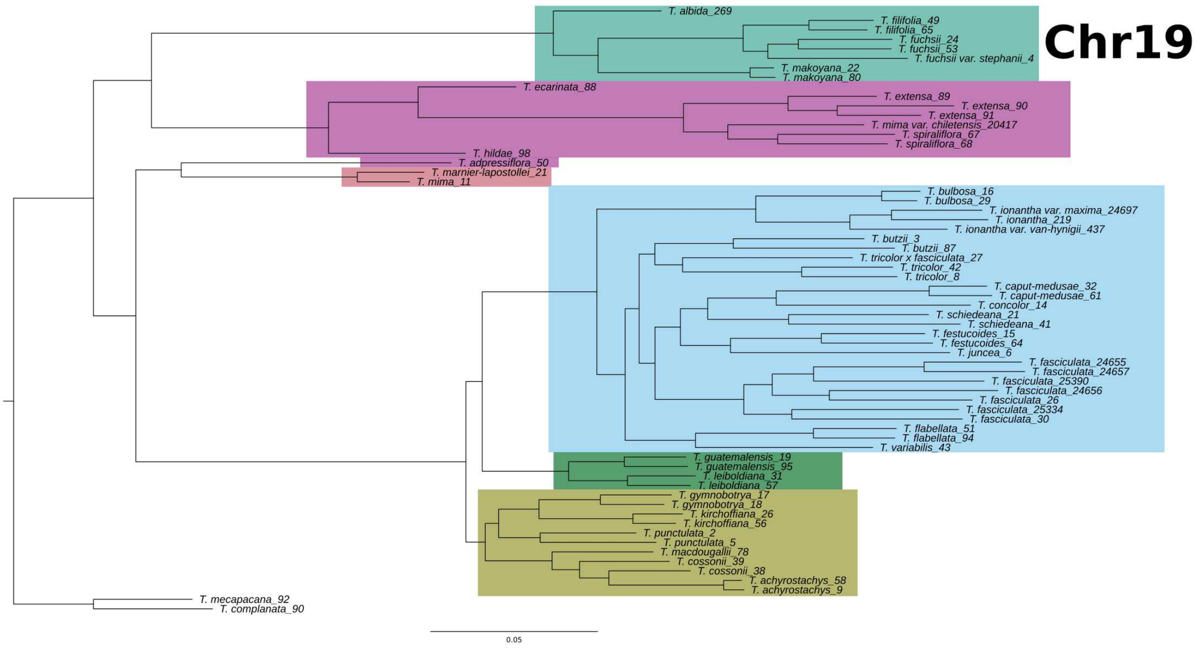
Chr19
T. albida_269
T. filifolia_49
T. filifolia_65
T. fuchsii_24
T. fuchsii_53
T. fuchsii var. stephanii_4
T. makoyana_22
T. makoyana_80
T. ecarinata_88
T. extensa_89
T. extensa_90
T. extensa_91
T. mima var. chiletensis_20417
T. spiraliflora_67
T. spiraliflora_68
T. hildae_98
T. adpressiflora_50
T. marnier-lapostollei_21
T. mima_11
T. bulbosa_16
T. bulbosa_29
T. ionantha var. maxima_24697
T. ionantha_219
T. ionantha var. van-hyningii_437
T. butzii_3
T. butzii_87
T. tricolor x fasciculata_27
T. tricolor_42
T. tricolor_8
T. caput-medusae_32
T. caput-medusae_61
T. concolor_14
T. schiedeana_21
T. schiedeana_41
T. festucoides_15
T. festucoides_64
T. juncea_6
T. fasciculata_24655
T. fasciculata_24657
T. fasciculata_25390
T. fasciculata_24656
T. fasciculata_26
T. fasciculata_25334
T. fasciculata_30
T. flabellata_51
T. flabellata_94
T. variabilis_43
T. guatemalensis_19
T. guatemalensis_95
T. leiboldiana_31
T. leiboldiana_57
T. gymnobotrya_17
T. gymnobotrya_18
T. kirchoffiana_26
T. kirchoffiana_56
T. punctulata_2
T. punctulata_5
T. macdougallii_78
T. cossonii_39
T. cossonii_38
T. achyrostachys_58
T. achyrostachys_9
T. mecapacana_92
T. complanata_90
0.05

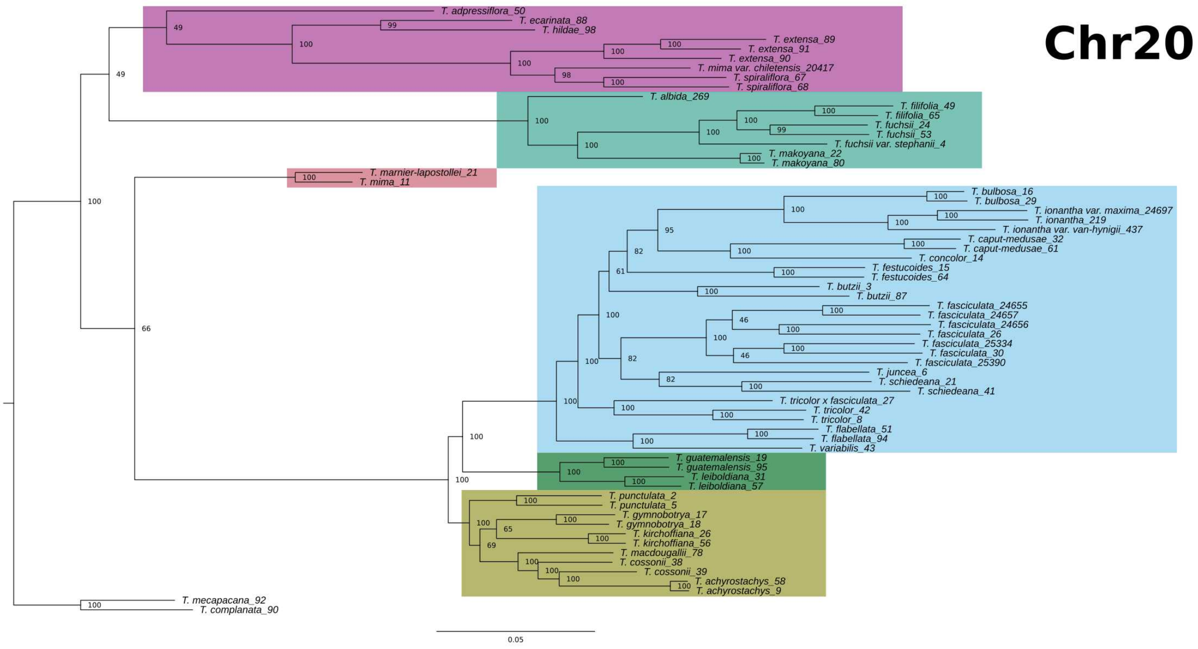
Chr20
T. adpressiflora_50
T. ecarinata_88
T. hildae_98
T. extensa_89
T. extensa_91
T. extensa_90
T. mima var. chiletensis_20417
T. spiraliflora_67
T. spiraliflora_68
T. albida_269
T. filifolia_49
T. filifolia_65
T. fuchsii_24
T. fuchsii_53
T. fuchsii var. stephanii_4
T. makoyana_22
T. makoyana_80
T. marnier-lapostollei_21
T. mima_11
T. bulbosa_16
T. bulbosa_29
T. ionantha var. maxima_24697
T. ionantha_219
T. ionantha var. van-hyningii_437
T. caput-medusae_32
T. caput-medusae_61
T. concolor_14
T. festucoides_15
T. festucoides_64
T. butzii_3
T. butzii_87
T. fasciculata_24655
T. fasciculata_24657
T. fasciculata_24656
T. fasciculata_26
T. fasciculata_25334
T. fasciculata_30
T. fasciculata_25390
T. juncea_6
T. schiedeana_21
T. schiedeana_41
T. tricolor x fasciculata_27
T. tricolor_42
T. tricolor_8
T. flabellata_51
T. flabellata_94
T. variabilis_43
T. guatemalensis_19
T. guatemalensis_95
T. leiboldiana_31
T. leiboldiana_57
T. punctulata_2
T. punctulata_5
T. gymnobotrya_17
T. gymnobotrya_18
T. kirchoffiana_26
T. kirchoffiana_56
T. macdougallii_78
T. cossonii_38
T. cossonii_39
T. achyrostachys_58
T. achyrostachys_9
T. mecapacana_92
T. complanata_90
0.05

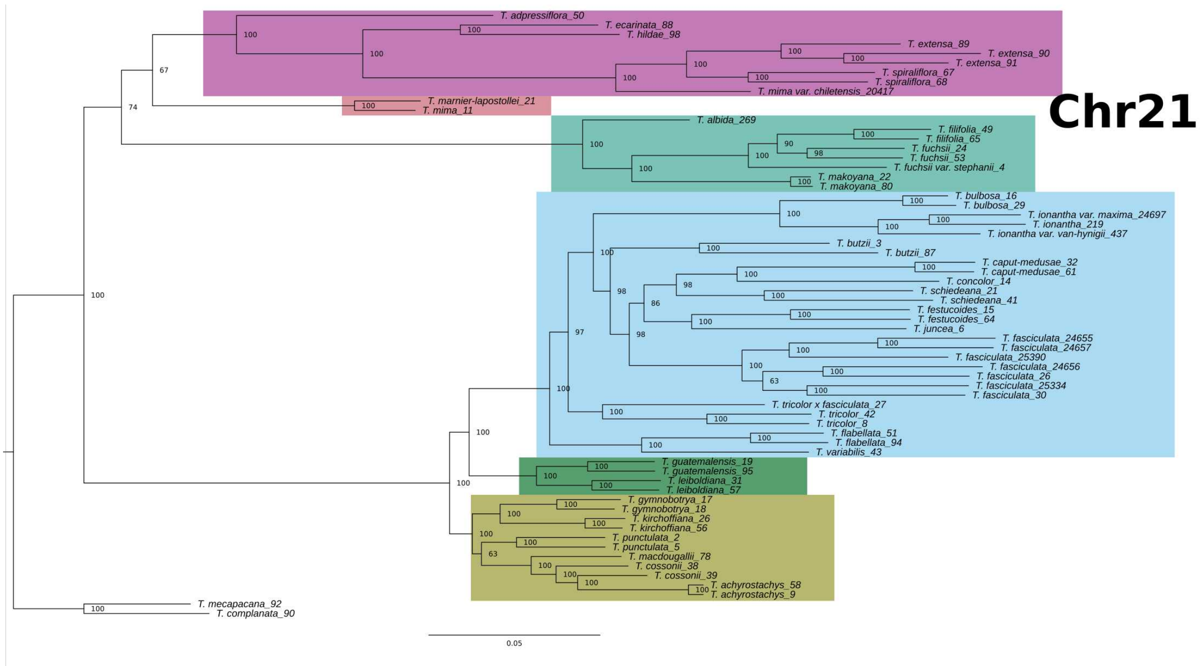
Chr21
T. adpressiflora_50
T. ecarinata_88
T. hildae_98
T. extensa_89
T. extensa_90
T. extensa_91
T. spiraliflora_67
T. spiraliflora_68
T. mima var. chiletensis_20417
T. marnier-lapostollei_21
T. mima_11
T. albida_269
T. filifolia_49
T. filifolia_65
T. fuchsii_24
T. fuchsii_53
T. fuchsii var. stephanii_4
T. makoyana_22
T. makoyana_80
T. bulbosa_16
T. bulbosa_29
T. ionantha var. maxima_24697
T. ionantha_219
T. ionantha var. van-hynigii_437
T. butzii_3
T. butzii_87
T. caput-medusae_32
T. caput-medusae_61
T. concolor_14
T. schiedeana_21
T. schiedeana_41
T. festucoides_15
T. festucoides_64
T. juncea_6
T. fasciculata_24655
T. fasciculata_24657
T. fasciculata_25390
T. fasciculata_24656
T. fasciculata_26
T. fasciculata_25334
T. fasciculata_30
T. tricolor x fasciculata_27
T. tricolor_42
T. tricolor_8
T. flabellata_51
T. flabellata_94
T. variabilis_43
T. guatemalensis_19
T. guatemalensis_95
T. leiboldiana_31
T. leiboldiana_57
T. gymnobotrya_17
T. gymnobotrya_18
T. kirchoffiana_26
T. kirchoffiana_56
T. punctulata_2
T. punctulata_5
T. macdougallii_78
T. cossonii_38
T. cossonii_39
T. achyrostachys_58
T. achyrostachys_9
T. mecapacana_92
T. complanata_90
0.05

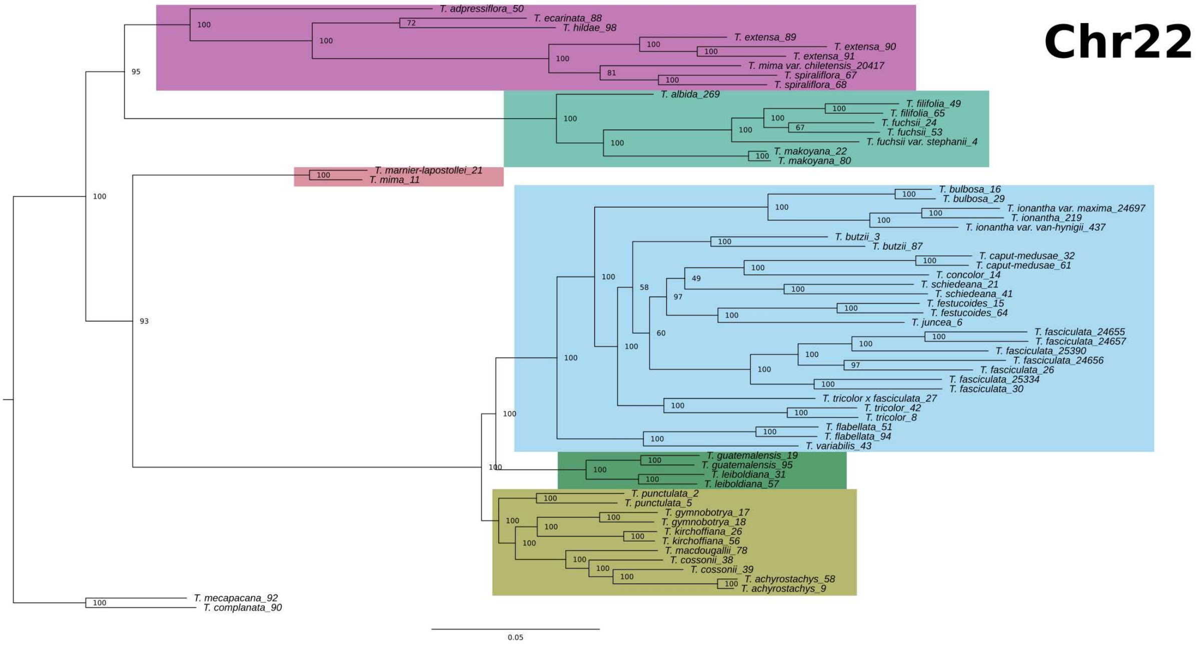
Chr22
T. adpressiflora_50
T. ecarinata_88
T. hildae_98
T. extensa_89
T. extensa_90
T. extensa_91
T. mima var. chiletensis_20417
T. spiraliflora_67
T. spiraliflora_68
T. albida_269
T. filifolia_49
T. filifolia_65
T. fuchsii_24
T. fuchsii_53
T. fuchsii var. stephanii_4
T. makoyana_22
T. makoyana_80
T. marnier-lapostollei_21
T. mima_11
T. bulbosa_16
T. bulbosa_29
T. ionantha var. maxima_24697
T. ionantha_219
T. ionantha var. van-hynigii_437
T. butzii_3
T. butzii_87
T. caput-medusae_32
T. caput-medusae_61
T. concolor_14
T. schiedeana_21
T. schiedeana_41
T. festucoides_15
T. festucoides_64
T. juncea_6
T. fasciculata_24655
T. fasciculata_24657
T. fasciculata_25390
T. fasciculata_24656
T. fasciculata_26
T. fasciculata_25334
T. fasciculata_30
T. tricolor x fasciculata_27
T. tricolor_42
T. tricolor_8
T. flabellata_51
T. flabellata_94
T. variabilis_43
T. guatemalensis_19
T. guatemalensis_95
T. leiboldiana_31
T. leiboldiana_57
T. punctulata_2
T. punctulata_5
T. gymnobotrya_17
T. gymnobotrya_18
T. kirchoffiana_26
T. kirchoffiana_56
T. macdougallii_78
T. cossonii_38
T. cossonii_39
T. achyrostachys_58
T. achyrostachys_9
T. mecapacana_92
T. complanata_90
0.05

Yardeni et al.,

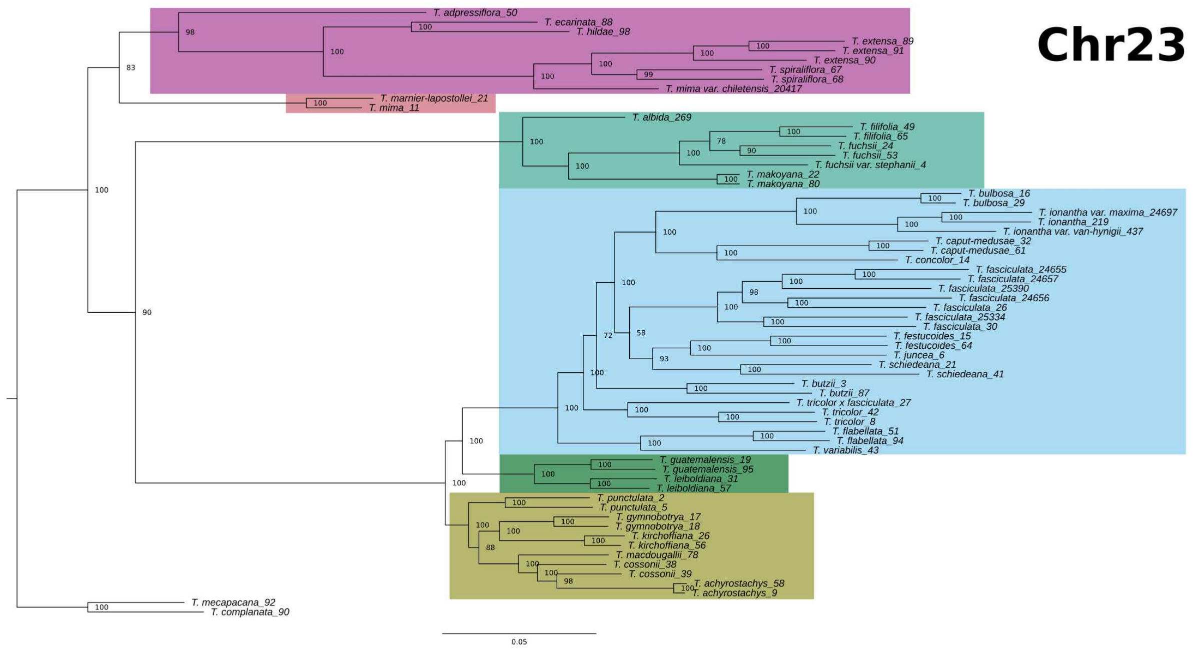


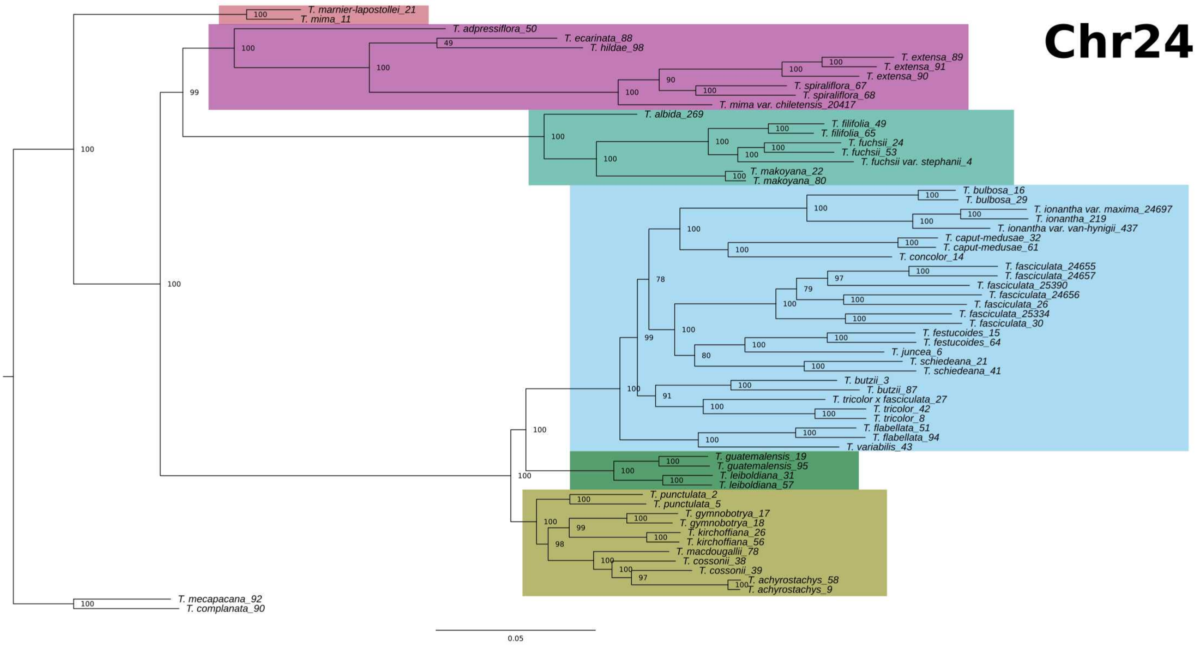

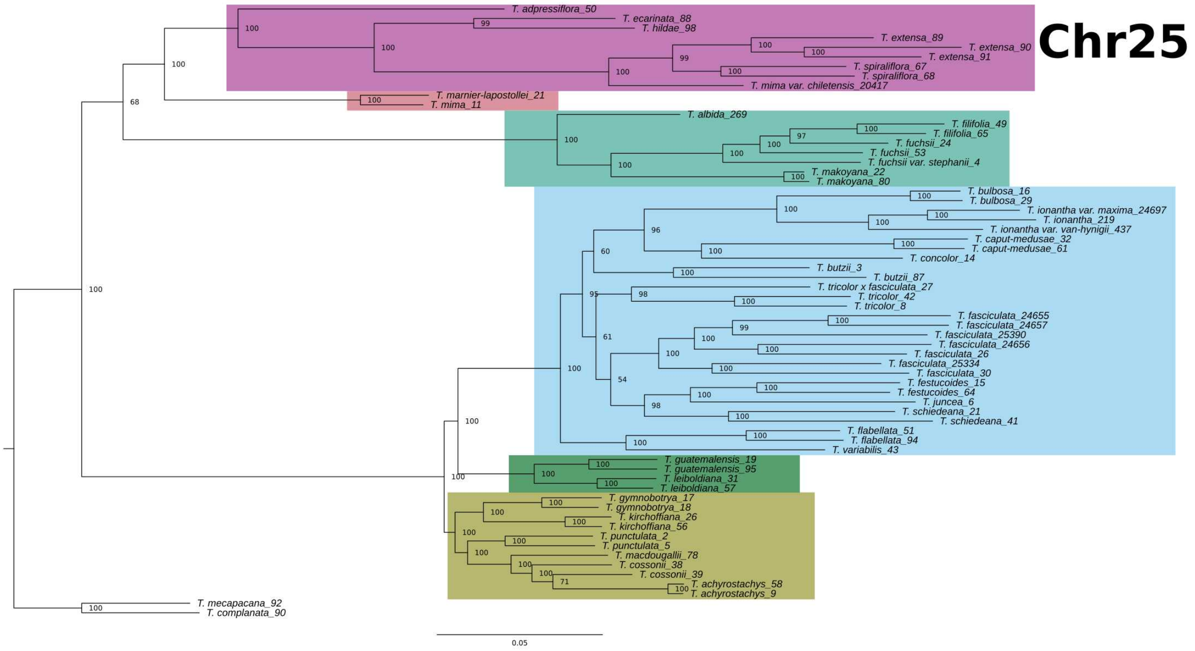
Chr25
T. adpressiflora_50
T. ecarinata_88
T. hildae_98
T. extensa_89
T. extensa_90
T. extensa_91
T. spiraliflora_67
T. spiraliflora_68
T. mima var. chiletensis_20417
T. marnier-lapostollei_21
T. mima_11
T. albida_269
T. filifolia_49
T. filifolia_65
T. fuchsii_24
T. fuchsii_53
T. fuchsii var. stephanii_4
T. makoyana_22
T. makoyana_80
T. bulbosa_16
T. bulbosa_29
T. ionantha var. maxima_24697
T. ionantha_219
T. ionantha var. van-hyningii_437
T. caput-medusae_32
T. caput-medusae_61
T. concolor_14
T. butzii_3
T. butzii_87
T. tricolor x fasciculata_27
T. tricolor_42
T. tricolor_8
T. fasciculata_24655
T. fasciculata_24657
T. fasciculata_25390
T. fasciculata_24656
T. fasciculata_26
T. fasciculata_25334
T. fasciculata_30
T. festucoides_15
T. festucoides_64
T. juncea_6
T. schiedeana_21
T. schiedeana_41
T. flabellata_51
T. flabellata_94
T. variabilis_43
T. guatemalensis_19
T. guatemalensis_95
T. leiboldiana_31
T. leiboldiana_57
T. gymnobotrya_17
T. gymnobotrya_18
T. kirchoffiana_26
T. kirchoffiana_56
T. punctulata_2
T. punctulata_5
T. macdougallii_78
T. cossonii_38
T. cossonii_39
T. achyrostachys_58
T. achyrostachys_9
T. mecapacana_92
T. complanata_90
0.05
