## supporting file 2 for "The explosive radiation of the Neotropical *Tillandsia* subgenus *Tillandsia* (Bromeliaceae) has been accompanied by pervasive hybridization"

**Supporting file 2** - Heatmaps summarizing 7,141 four-taxon D-statistic tests for each of the 25 reference chromosomes, indicated on each figure. *Tillandsia complanata* was used as the outgroup in all tests. The four taxa in each test have been rearranged to always obtain positive D values, and P2 and P3 are shown on the axes. Colour indicates the value of D and log value of p-value, as appears in legend (bottom right).

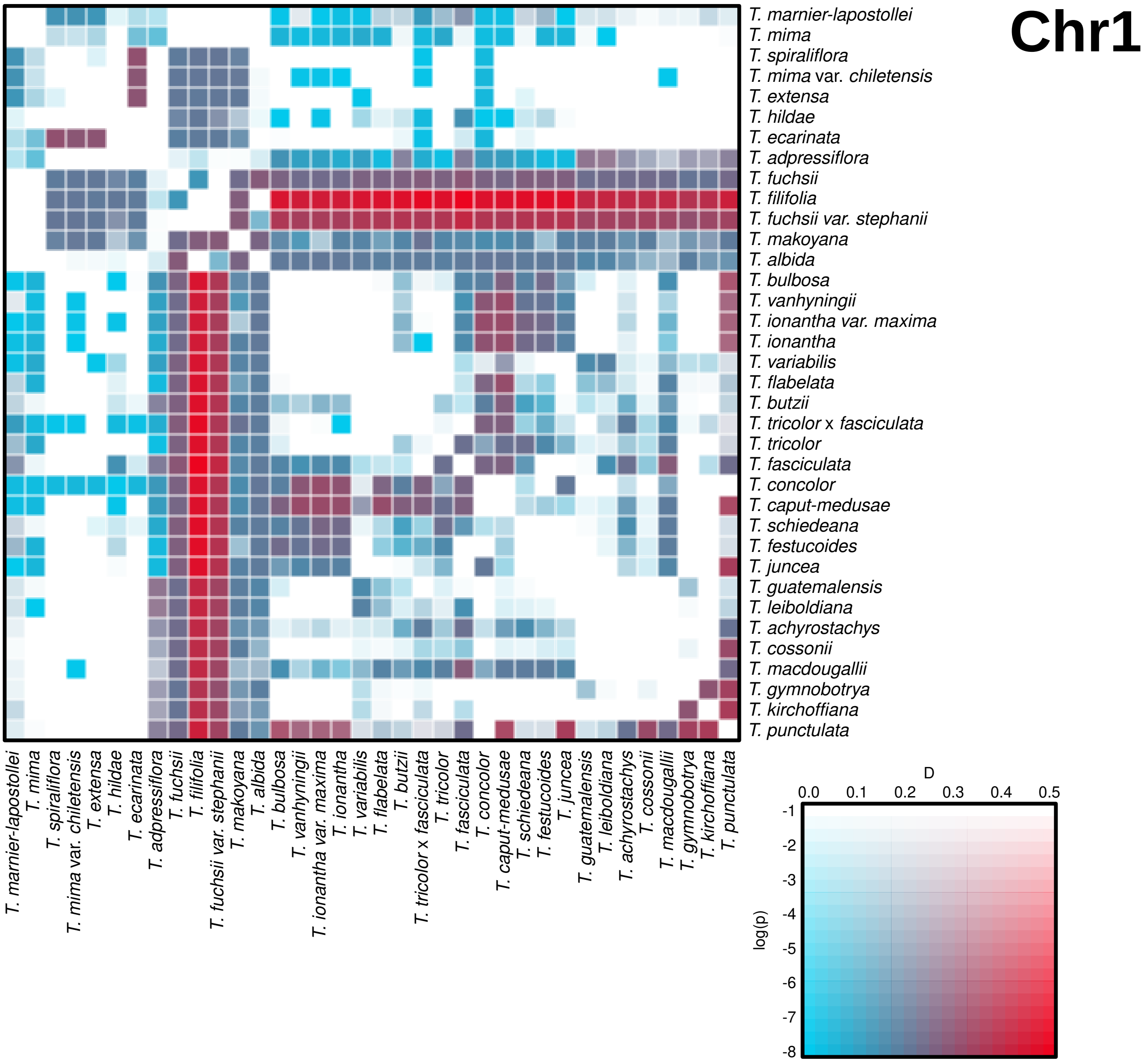
Chr1
T. marnier-lapostollei
T. mima
T. spiraliflora
T. mima var. chiletensis
T. extensa
T. hildae
T. ecarinata
T. adpressiflora
T. fuchsii
T. filifolia
T. fuchsii var. stephanii
T. makoyana
T. albida
T. bulbosa
T. vanhyningii
T. ionantha var. maxima
T. ionantha
T. variabilis
T. flabelata
T. butzii
T. tricolor x fasciculata
T. tricolor
T. fasciculata
T. concolor
T. caput-medusae
T. schiedeana
T. festucoides
T. juncea
T. guatemalensis
T. leiboldiana
T. achyrostachys
T. cossonii
T. macdougallii
T. gymnobotrya
T. kirchoffiana
T. punctulata
D
0.0
0.1
0.2
0.3
0.4
0.5
log(p)
-1
-2
-3
-4
-5
-6
-7
-8

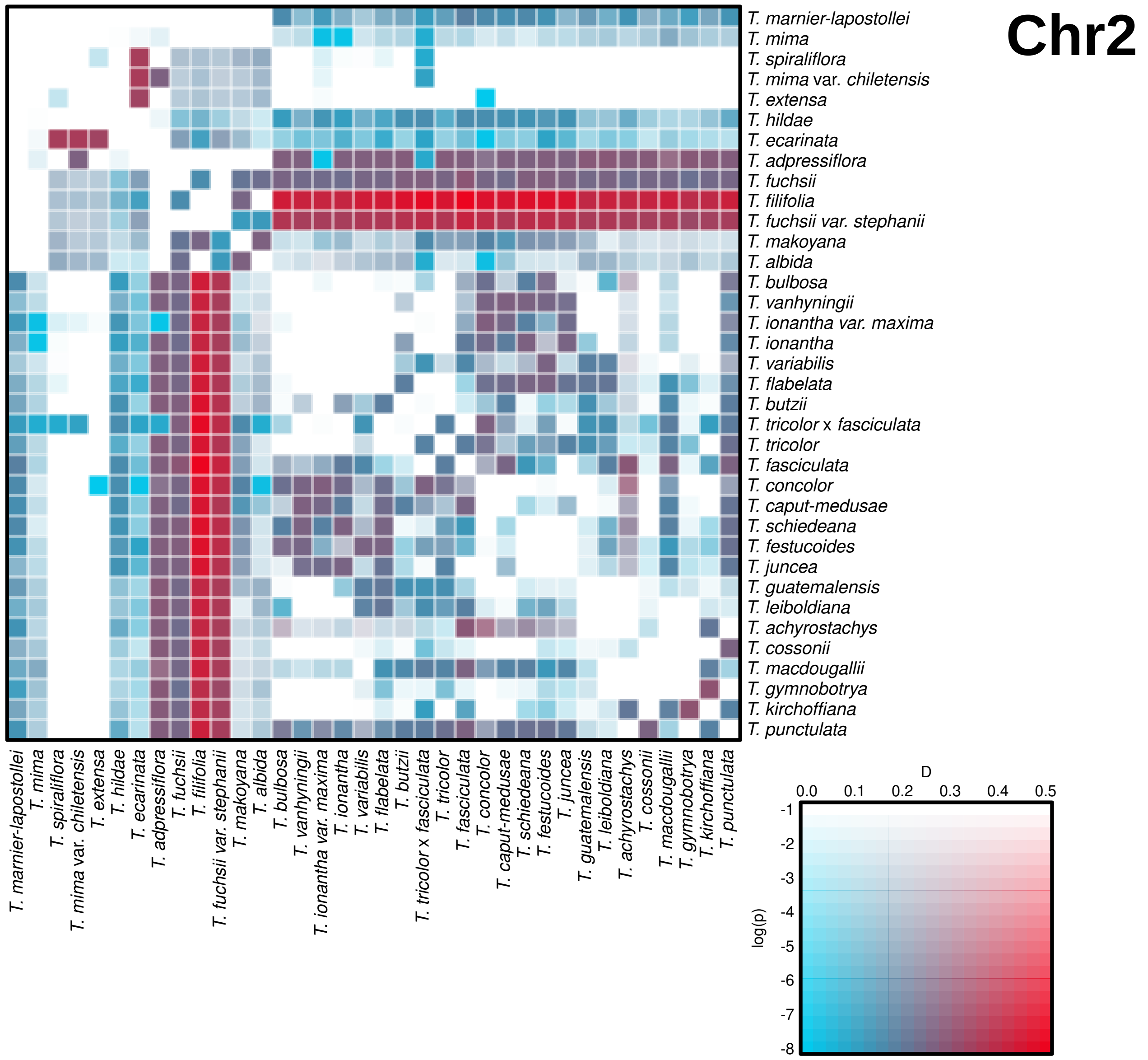
Chr2
T. marnier-lapostollei
T. mima
T. spiraliflora
T. mima var. chiletensis
T. extensa
T. hildae
T. ecarinata
T. adpressiflora
T. fuchsii
T. filifolia
T. fuchsii var. stephanii
T. makoyana
T. albida
T. bulbosa
T. vanhyningii
T. ionantha var. maxima
T. ionantha
T. variabilis
T. flabelata
T. butzii
T. tricolor x fasciculata
T. tricolor
T. fasciculata
T. concolor
T. caput-medusae
T. schiedeana
T. festucoides
T. juncea
T. guatemalensis
T. leiboldiana
T. achyrostachys
T. cossonii
T. macdougallii
T. gymnobotrya
T. kirchoffiana
T. punctulata
D
0.0 0.1 0.2 0.3 0.4 0.5
log(p)
-1 -2 -3 -4 -5 -6 -7 -8

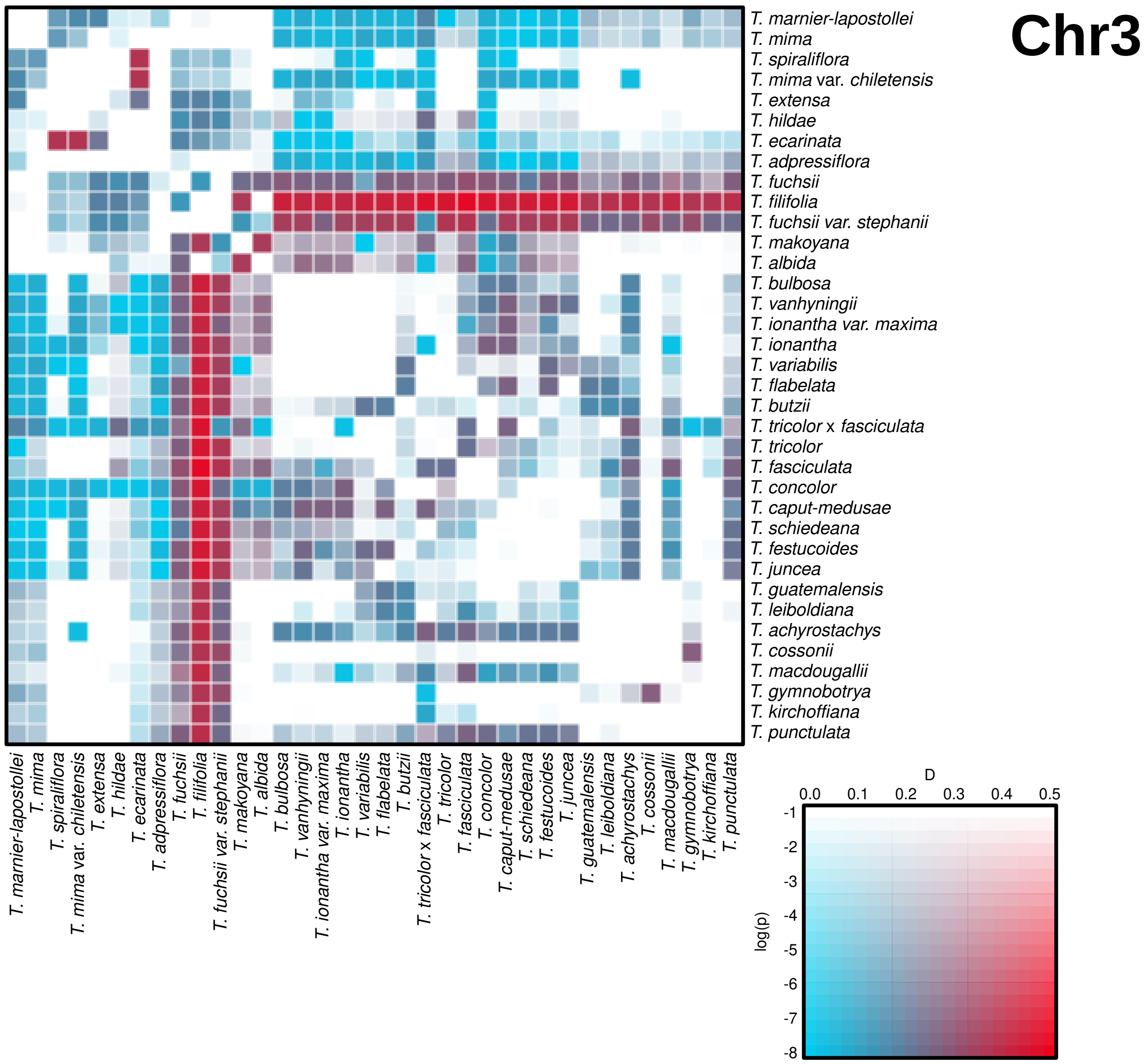
Chr3
T. marnier-lapostollei
T. mima
T. spiraliflora
T. mima var. chiletensis
T. extensa
T. hildae
T. ecarinata
T. adpressiflora
T. fuchsii
T. filifolia
T. fuchsii var. stephanii
T. makoyana
T. albida
T. bulbosa
T. vanhyningii
T. ionantha var. maxima
T. ionantha
T. variabilis
T. flabelata
T. butzii
T. tricolor x fasciculata
T. tricolor
T. fasciculata
T. concolor
T. caput-medusae
T. schiedeana
T. festucoides
T. juncea
T. guatemalensis
T. leiboldiana
T. achyrostachys
T. cossonii
T. macdougallii
T. gymnobotrya
T. kirchoffiana
T. punctulata
D
0.0
0.1
0.2
0.3
0.4
0.5
log(p)
-1
-2
-3
-4
-5
-6
-7
-8

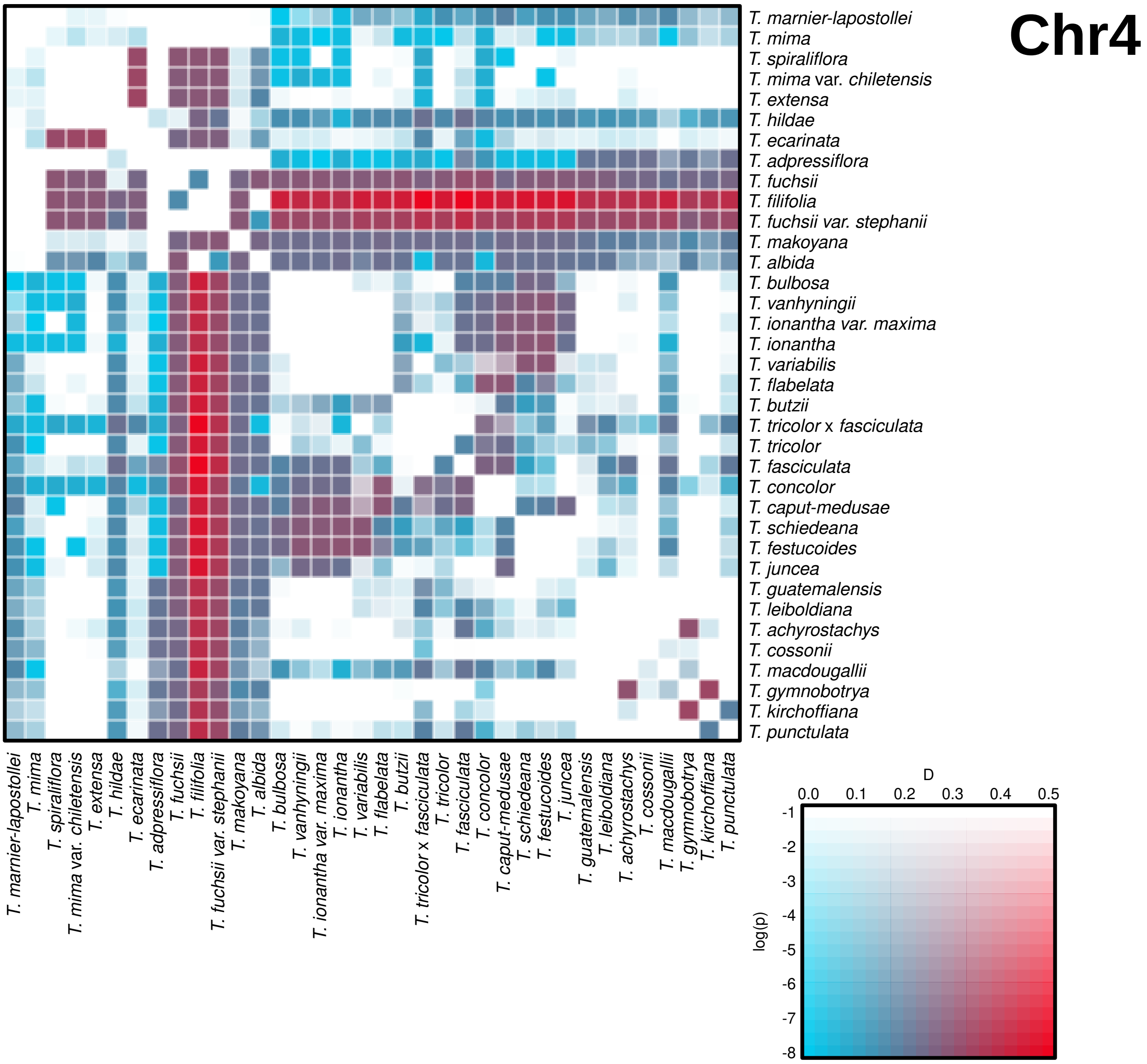
Chr4
T. marnier-lapostollei
T. mima
T. spiraliflora
T. mima var. chiletensis
T. extensa
T. hildae
T. ecarinata
T. adpressiflora
T. fuchsii
T. filifolia
T. fuchsii var. stephanii
T. makoyana
T. albida
T. bulbosa
T. vanhyningii
T. ionantha var. maxima
T. ionantha
T. variabilis
T. flabelata
T. butzii
T. tricolor x fasciculata
T. tricolor
T. fasciculata
T. concolor
T. caput-medusae
T. schiedeana
T. festucoides
T. juncea
T. guatemalensis
T. leiboldiana
T. achyrostachys
T. cossonii
T. macdougallii
T. gymnobotrya
T. kirchoffiana
T. punctulata
D
0.0
0.1
0.2
0.3
0.4
0.5
log(p)
-1
-2
-3
-4
-5
-6
-7
-8

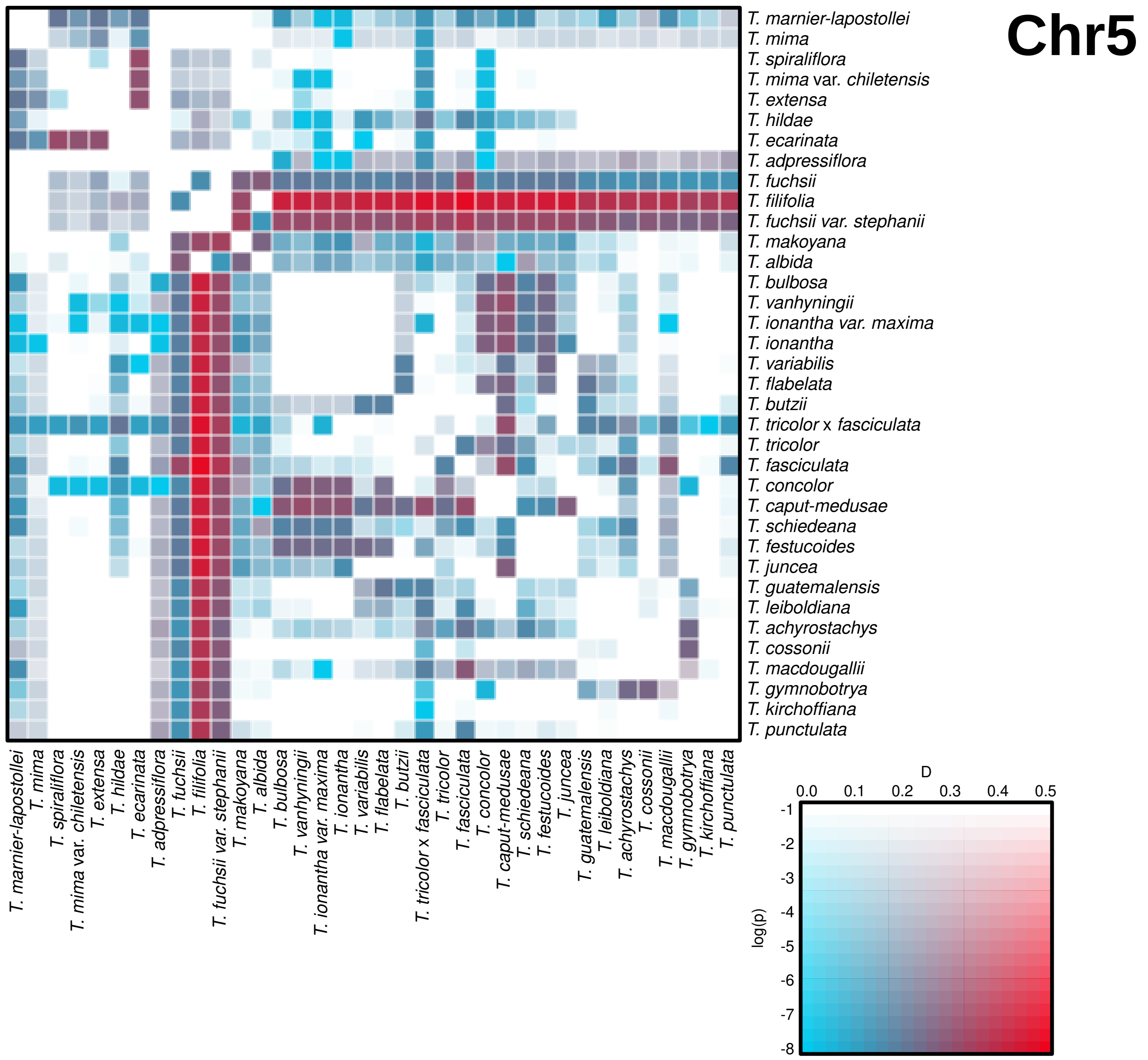

Chr5
T. marnier-lapostollei
T. mima
T. spiraliflora
T. mima var. chiletensis
T. extensa
T. hildae
T. ecarinata
T. adpressiflora
T. fuchsii
T. filifolia
T. fuchsii var. stephanii
T. makoyana
T. albida
T. bulbosa
T. vanhyningii
T. ionantha var. maxima
T. ionantha
T. variabilis
T. flabelata
T. butzii
T. tricolor x fasciculata
T. tricolor
T. fasciculata
T. concolor
T. caput-medusae
T. schiedeana
T. festucoides
T. juncea
T. guatemalensis
T. leiboldiana
T. achyrostachys
T. cossonii
T. macdougallii
T. gymnobotrya
T. kirchoffiana
T. punctulata
D
0.0
0.1
0.2
0.3
0.4
0.5
log(p)
-1
-2
-3
-4
-5
-6
-7
-8

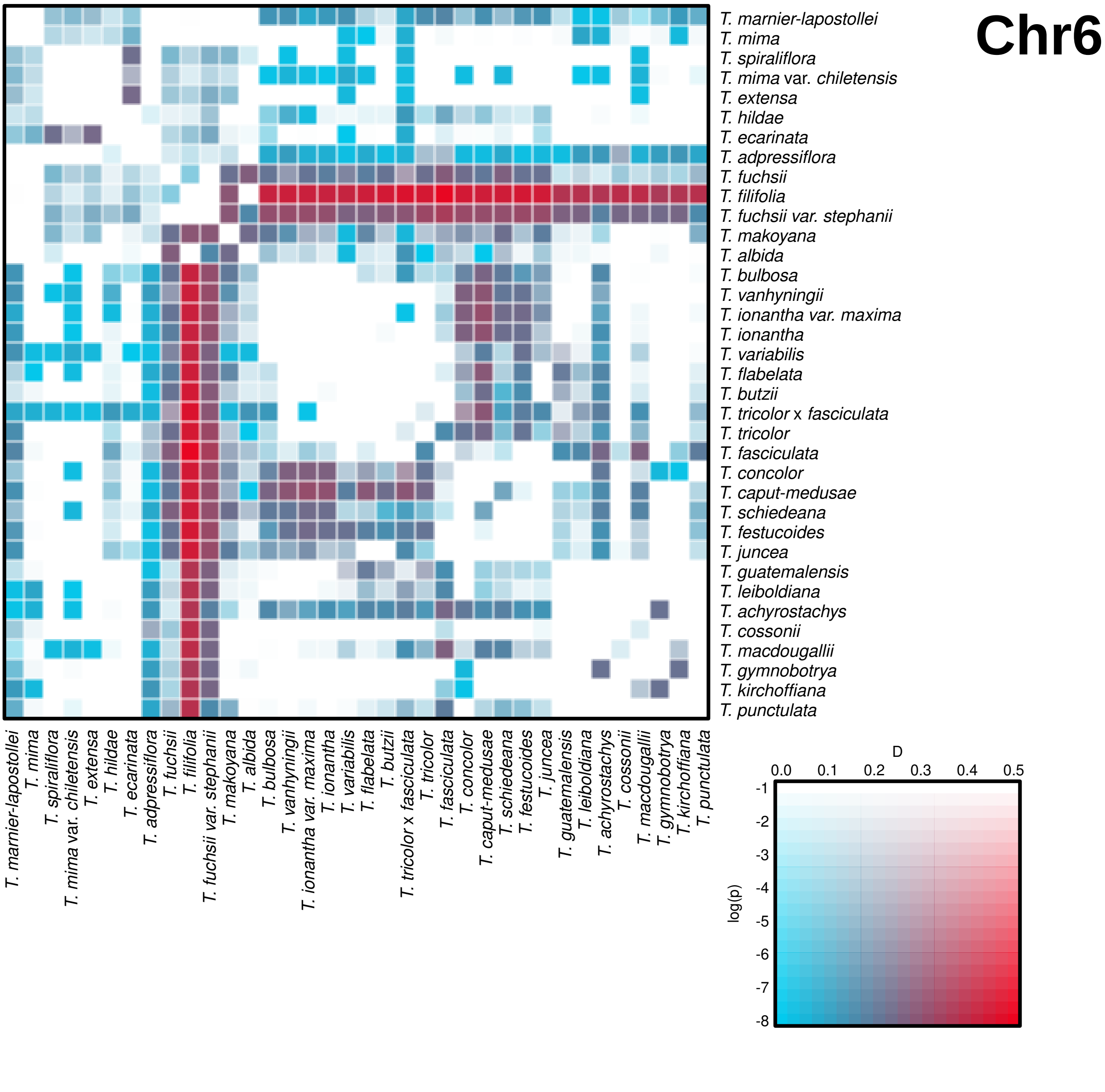
Chr6
T. marnier-lapostollei
T. mima
T. spiraliflora
T. mima var. chiletensis
T. extensa
T. hildae
T. ecarinata
T. adpressiflora
T. fuchsii
T. filifolia
T. fuchsii var. stephanii
T. makoyana
T. albida
T. bulbosa
T. vanhyningii
T. ionantha var. maxima
T. ionantha
T. variabilis
T. flabelata
T. butzii
T. tricolor x fasciculata
T. tricolor
T. fasciculata
T. concolor
T. caput-medusae
T. schiedeana
T. festucoides
T. juncea
T. guatemalensis
T. leiboldiana
T. achyrostachys
T. cossonii
T. macdougallii
T. gymnobotrya
T. kirchoffiana
T. punctulata
D
0.0
0.1
0.2
0.3
0.4
0.5
log(p)
-1
-2
-3
-4
-5
-6
-7
-8

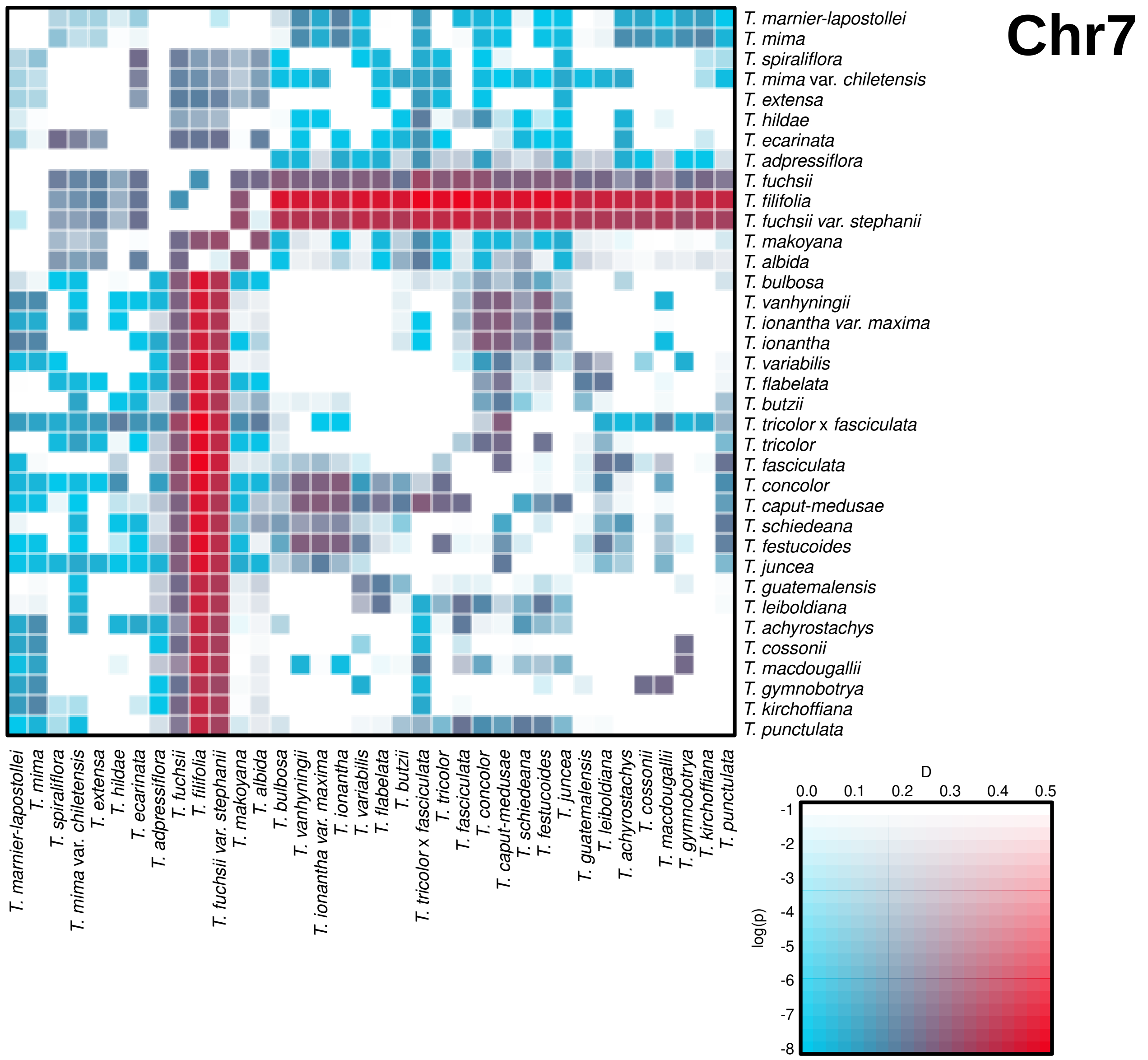
Chr7
T. marnier-lapostollei
T. mima
T. spiraliflora
T. mima var. chiletensis
T. extensa
T. hildae
T. ecarinata
T. adpressiflora
T. fuchsii
T. filifolia
T. fuchsii var. stephanii
T. makoyana
T. albida
T. bulbosa
T. vanhyningii
T. ionantha var. maxima
T. ionantha
T. variabilis
T. flabelata
T. butzii
T. tricolor x fasciculata
T. tricolor
T. fasciculata
T. concolor
T. caput-medusae
T. schiedeana
T. festucoides
T. juncea
T. guatemalensis
T. leiboldiana
T. achyrostachys
T. cossonii
T. macdougallii
T. gymnobotrya
T. kirchoffiana
T. punctulata
D
0.0
0.1
0.2
0.3
0.4
0.5
log(p)
-1
-2
-3
-4
-5
-6
-7
-8

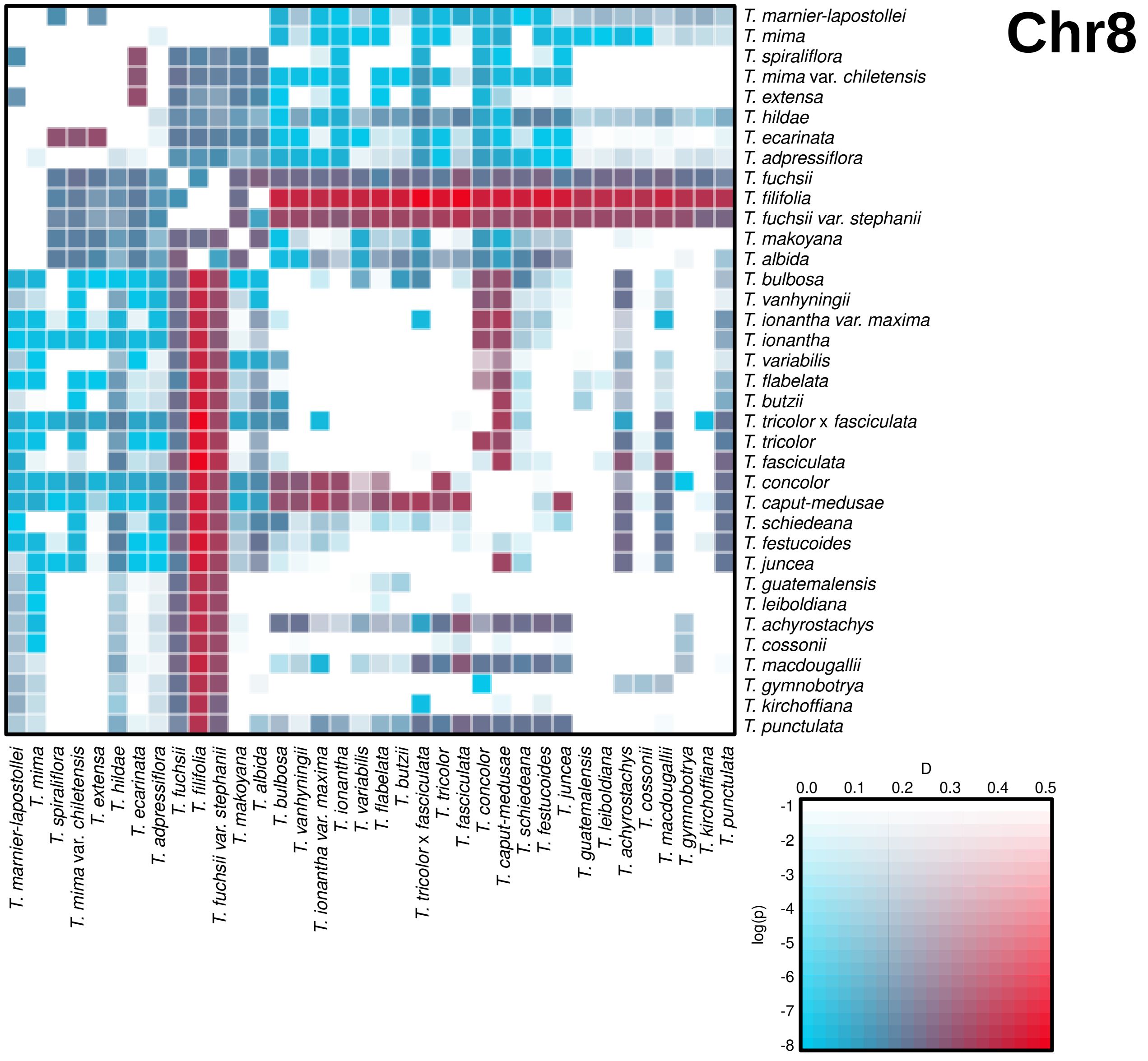
Chr8
T. marnier-lapostollei
T. mima
T. spiraliflora
T. mima var. chiletensis
T. extensa
T. hildae
T. ecarinata
T. adpressiflora
T. fuchsii
T. filifolia
T. fuchsii var. stephanii
T. makoyana
T. albida
T. bulbosa
T. vanhyningii
T. ionantha var. maxima
T. ionantha
T. variabilis
T. flabelata
T. butzii
T. tricolor x fasciculata
T. tricolor
T. fasciculata
T. concolor
T. caput-medusae
T. schiedeana
T. festucoides
T. juncea
T. guatemalensis
T. leiboldiana
T. achyrostachys
T. cossonii
T. macdougallii
T. gymnobotrya
T. kirchoffiana
T. punctulata
D
0.0
0.1
0.2
0.3
0.4
0.5
log(p)
-1
-2
-3
-4
-5
-6
-7
-8

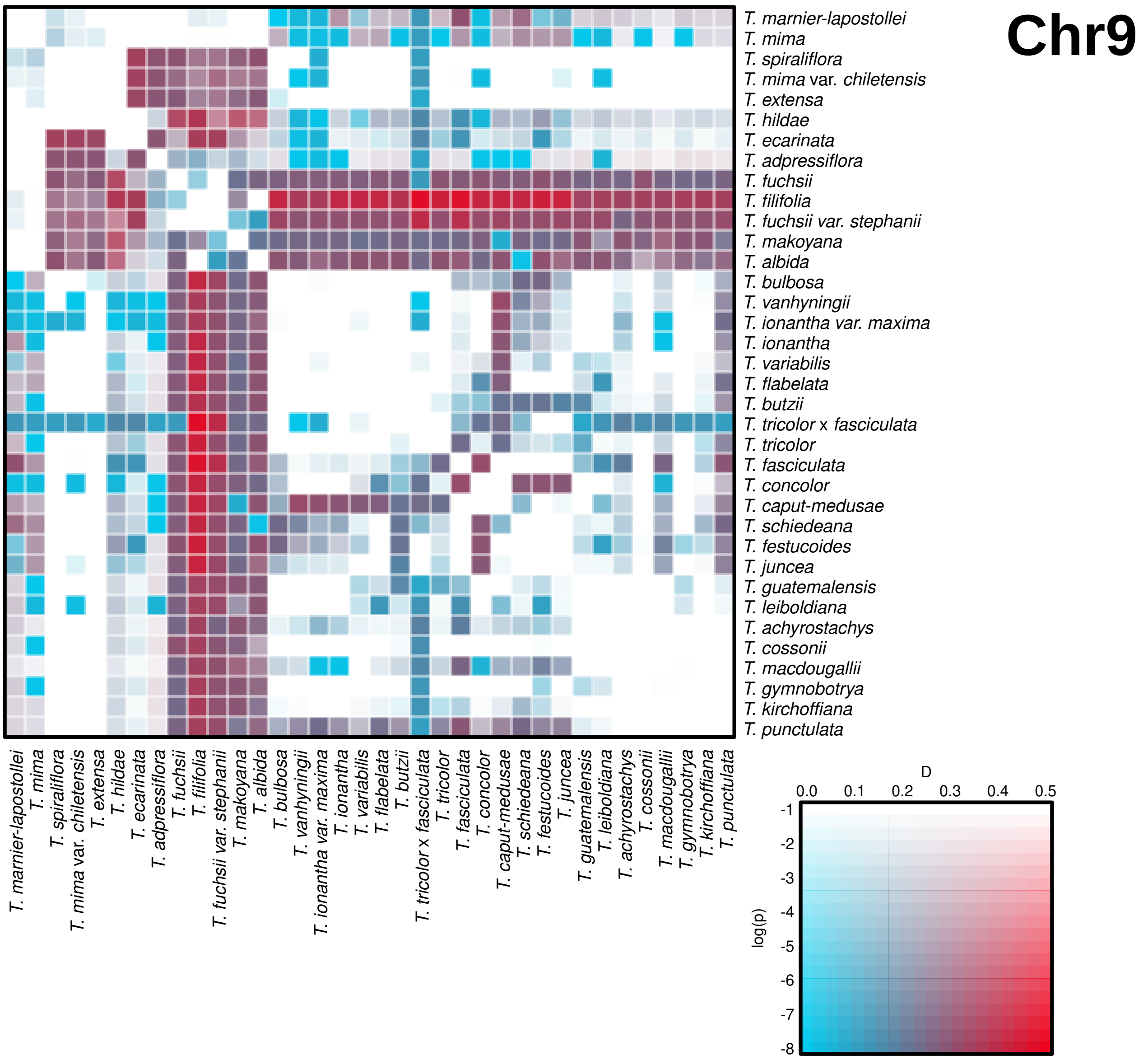
Chr9
T. marnier-lapostollei
T. mima
T. spiraliflora
T. mima var. chiletensis
T. extensa
T. hildae
T. ecarinata
T. adpressiflora
T. fuchsii
T. filifolia
T. fuchsii var. stephanii
T. makoyana
T. albida
T. bulbosa
T. vanhyningii
T. ionantha var. maxima
T. ionantha
T. variabilis
T. flabelata
T. butzii
T. tricolor x fasciculata
T. tricolor
T. fasciculata
T. concolor
T. caput-medusae
T. schiedeana
T. festucoides
T. juncea
T. guatemalensis
T. leiboldiana
T. achyrostachys
T. cossonii
T. macdougallii
T. gymnobotrya
T. kirchoffiana
T. punctulata
D
0.0
0.1
0.2
0.3
0.4
0.5
log(p)
-1
-2
-3
-4
-5
-6
-7
-8

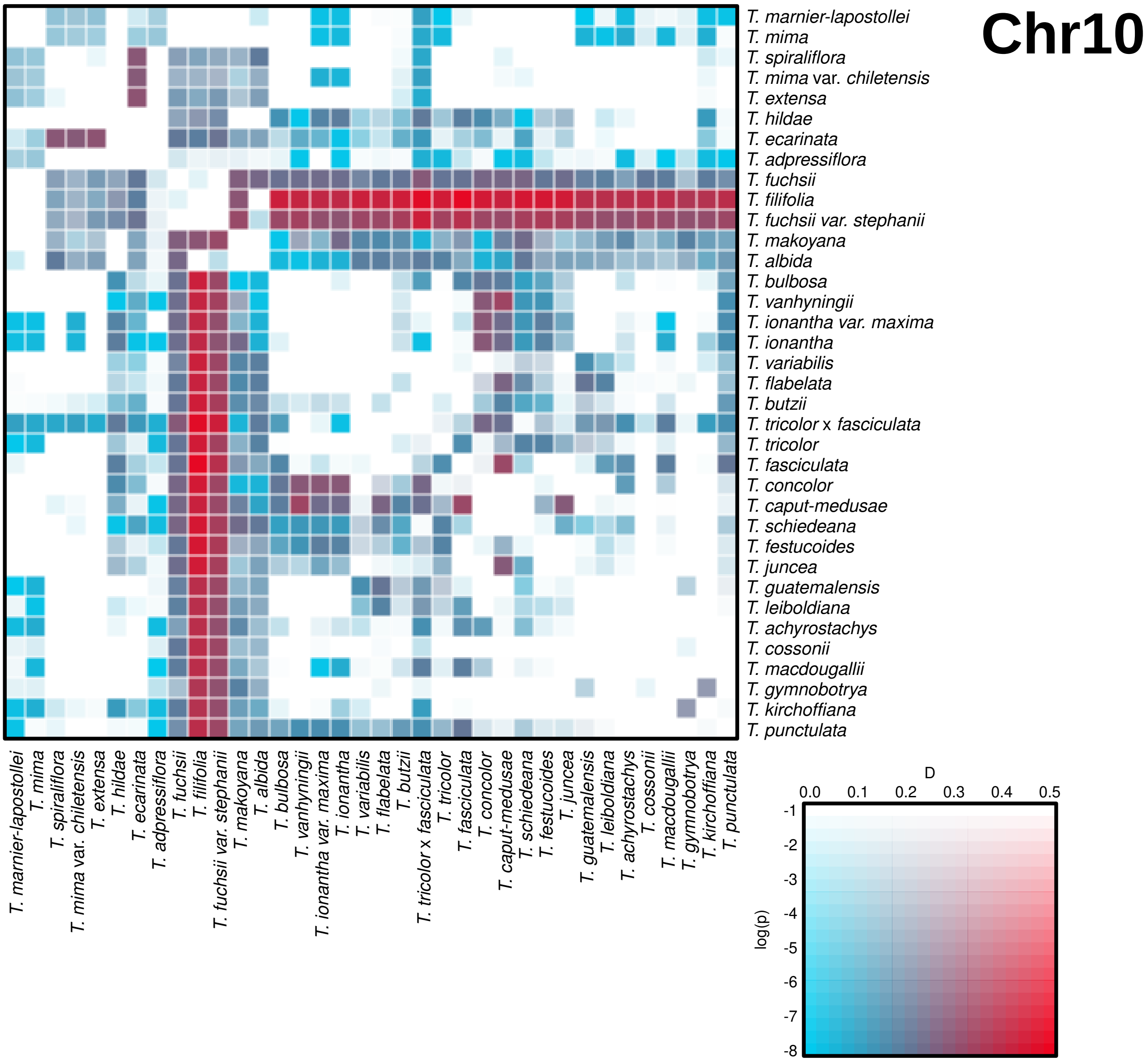
Chr10
T. marnier-lapostollei
T. mima
T. spiraliflora
T. mima var. chiletensis
T. extensa
T. hildae
T. ecarinata
T. adpressiflora
T. fuchsii
T. filifolia
T. fuchsii var. stephanii
T. makoyana
T. albida
T. bulbosa
T. vanhyningii
T. ionantha var. maxima
T. ionantha
T. variabilis
T. flabelata
T. butzii
T. tricolor x fasciculata
T. tricolor
T. fasciculata
T. concolor
T. caput-medusae
T. schiedeana
T. festucoides
T. juncea
T. guatemalensis
T. leiboldiana
T. achyrostachys
T. cossonii
T. macdougallii
T. gymnobotrya
T. kirchoffiana
T. punctulata
T. marnier-lapostollei
T. mima
T. spiraliflora
T. mima var. chiletensis
T. extensa
T. hildae
T. ecarinata
T. adpressiflora
T. fuchsii
T. filifolia
T. fuchsii var. stephanii
T. makoyana
T. albida
T. bulbosa
T. vanhyningii
T. ionantha var. maxima
T. ionantha
T. variabilis
T. flabelata
T. butzii
T. tricolor x fasciculata
T. tricolor
T. fasciculata
T. concolor
T. caput-medusae
T. schiedeana
T. festucoides
T. juncea
T. guatemalensis
T. leiboldiana
T. achyrostachys
T. cossonii
T. macdougallii
T. gymnobotrya
T. kirchoffiana
T. punctulata
D
0.0
0.1
0.2
0.3
0.4
0.5
log(p)
-1
-2
-3
-4
-5
-6
-7
-8

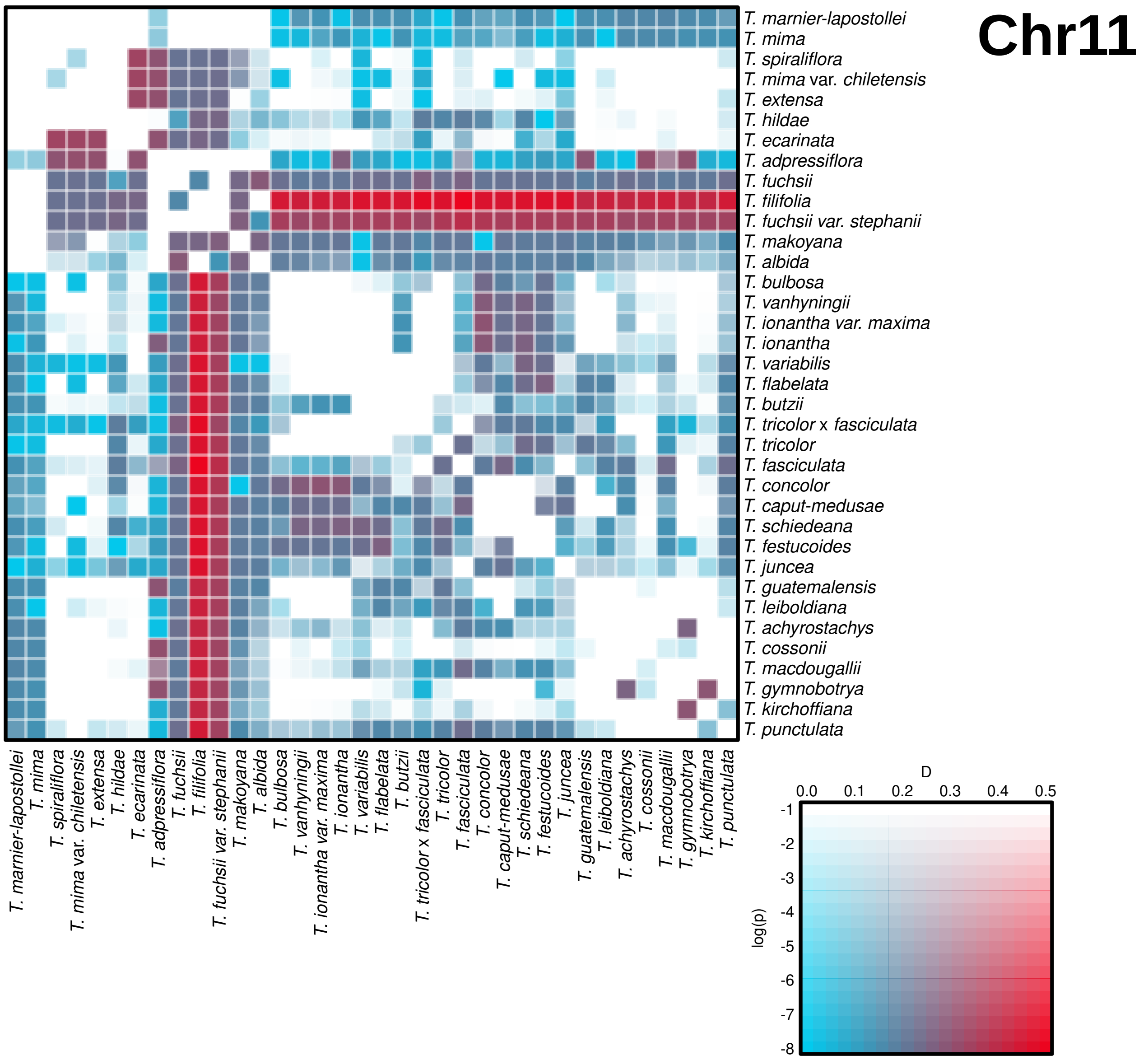

Chr11
T. marnier-lapostollei
T. mima
T. spiraliflora
T. mima var. chiletensis
T. extensa
T. hildae
T. ecarinata
T. adpressiflora
T. fuchsii
T. filifolia
T. fuchsii var. stephanii
T. makoyana
T. albida
T. bulbosa
T. vanhyningii
T. ionantha var. maxima
T. ionantha
T. variabilis
T. flabelata
T. butzii
T. tricolor x fasciculata
T. tricolor
T. fasciculata
T. concolor
T. caput-medusae
T. schiedeana
T. festucoides
T. juncea
T. guatemalensis
T. leiboldiana
T. achyrostachys
T. cossonii
T. macdougallii
T. gymnobotrya
T. kirchoffiana
T. punctulata
D
0.0
0.1
0.2
0.3
0.4
0.5
log(p)
-1
-2
-3
-4
-5
-6
-7
-8

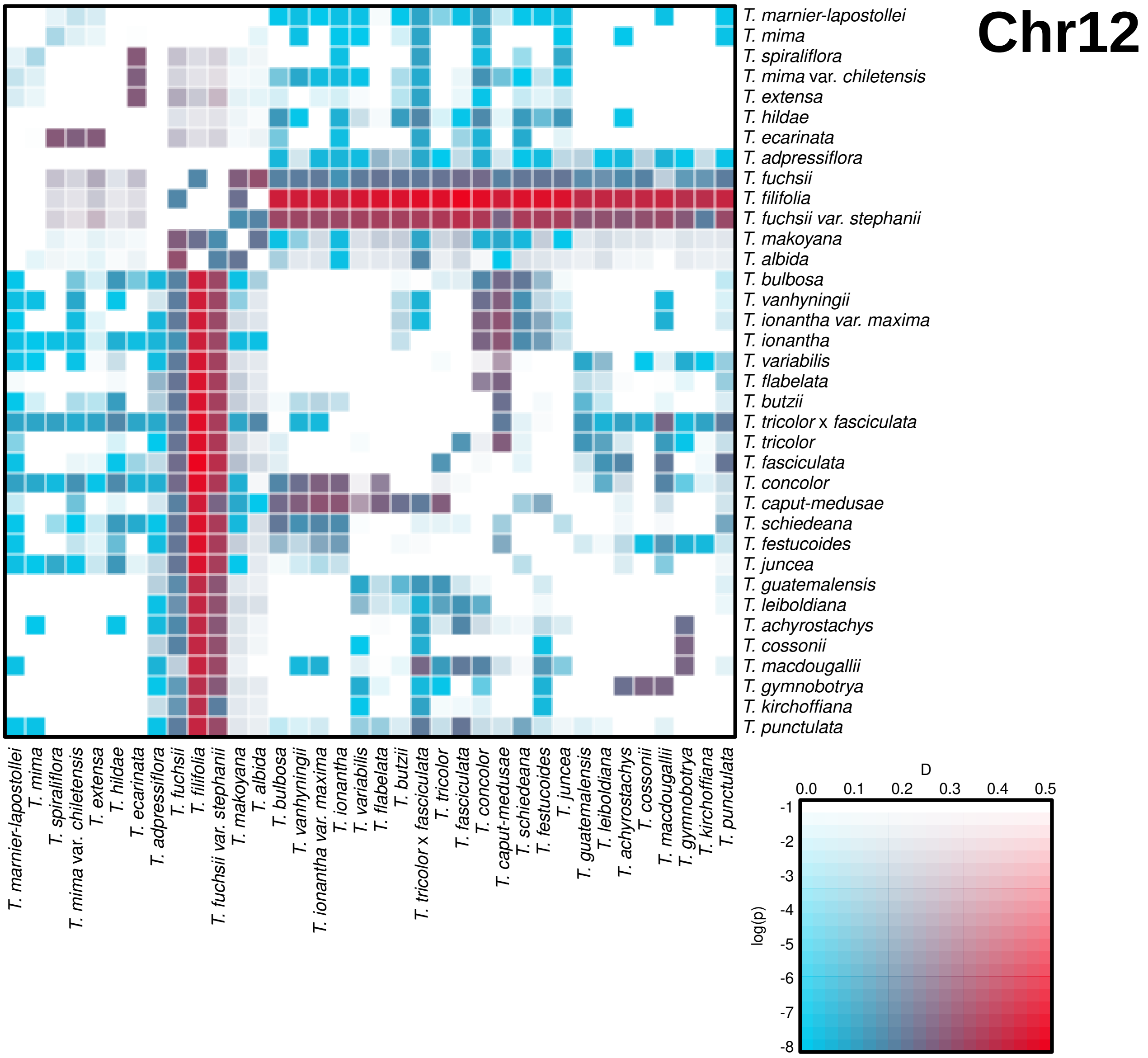
Chr12
T. marnier-lapostollei
T. mima
T. spiraliflora
T. mima var. chiletensis
T. extensa
T. hildae
T. ecarinata
T. adpressiflora
T. fuchsii
T. filifolia
T. fuchsii var. stephanii
T. makoyana
T. albida
T. bulbosa
T. vanhyningii
T. ionantha var. maxima
T. ionantha
T. variabilis
T. flabelata
T. butzii
T. tricolor x fasciculata
T. tricolor
T. fasciculata
T. concolor
T. caput-medusae
T. schiedeana
T. festucoides
T. juncea
T. guatemalensis
T. leiboldiana
T. achyrostachys
T. cossonii
T. macdougallii
T. gymnobotrya
T. kirchoffiana
T. punctulata
D
0.0
0.1
0.2
0.3
0.4
0.5
log(p)
-1
-2
-3
-4
-5
-6
-7
-8

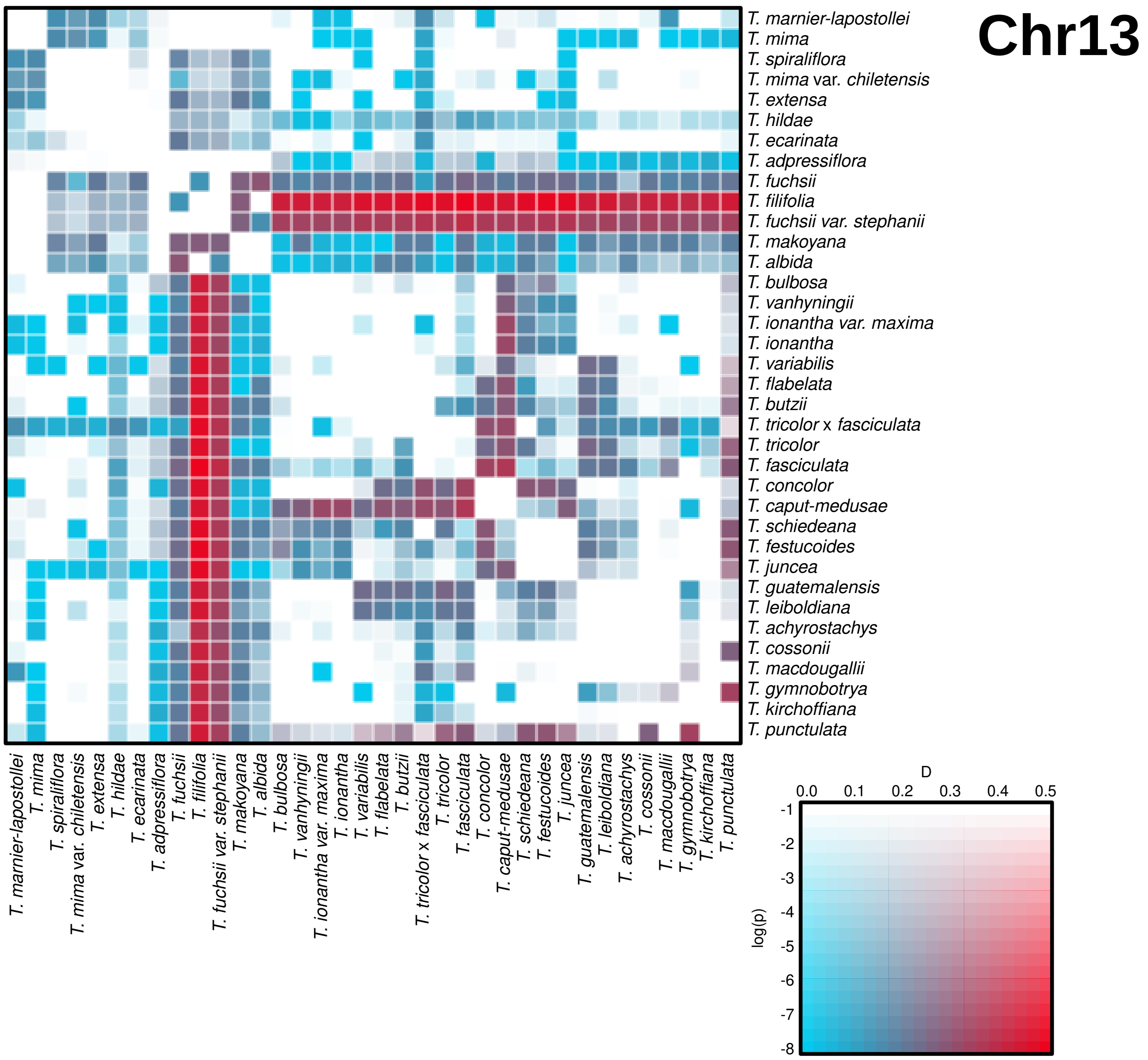
Chr13
T. marnier-lapostollei
T. mima
T. spiraliflora
T. mima var. chiletensis
T. extensa
T. hildae
T. ecarinata
T. adpressiflora
T. fuchsii
T. filifolia
T. fuchsii var. stephanii
T. makoyana
T. albida
T. bulbosa
T. vanhyningii
T. ionantha var. maxima
T. ionantha
T. variabilis
T. flabelata
T. butzii
T. tricolor x fasciculata
T. tricolor
T. fasciculata
T. concolor
T. caput-medusae
T. schiedeana
T. festucoides
T. juncea
T. guatemalensis
T. leiboldiana
T. achyrostachys
T. cossonii
T. macdougallii
T. gymnobotrya
T. kirchoffiana
T. punctulata
D
0.0
0.1
0.2
0.3
0.4
0.5
log(p)
-1
-2
-3
-4
-5
-6
-7
-8

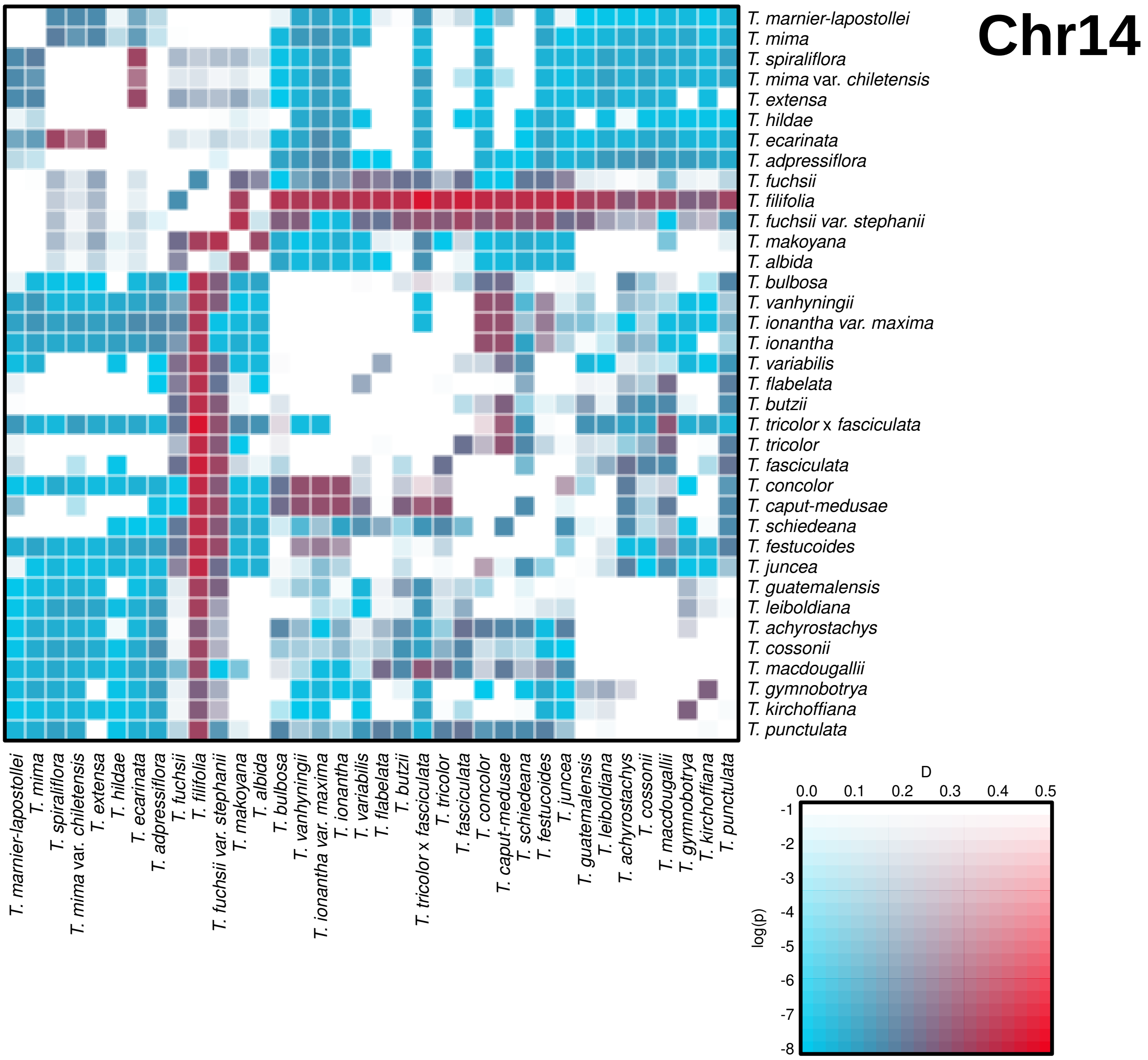

Chr14
T. marnier-lapostollei
T. mima
T. spiraliflora
T. mima var. chiletensis
T. extensa
T. hildae
T. ecarinata
T. adpressiflora
T. fuchsii
T. filifolia
T. fuchsii var. stephanii
T. makoyana
T. albida
T. bulbosa
T. vanhyningii
T. ionantha var. maxima
T. ionantha
T. variabilis
T. flabelata
T. butzii
T. tricolor x fasciculata
T. tricolor
T. fasciculata
T. concolor
T. caput-medusae
T. schiedeana
T. festucoides
T. juncea
T. guatemalensis
T. leiboldiana
T. achyrostachys
T. cossonii
T. macdougallii
T. gymnobotrya
T. kirchoffiana
T. punctulata
D
0.0
0.1
0.2
0.3
0.4
0.5
log(p)
-1
-2
-3
-4
-5
-6
-7
-8

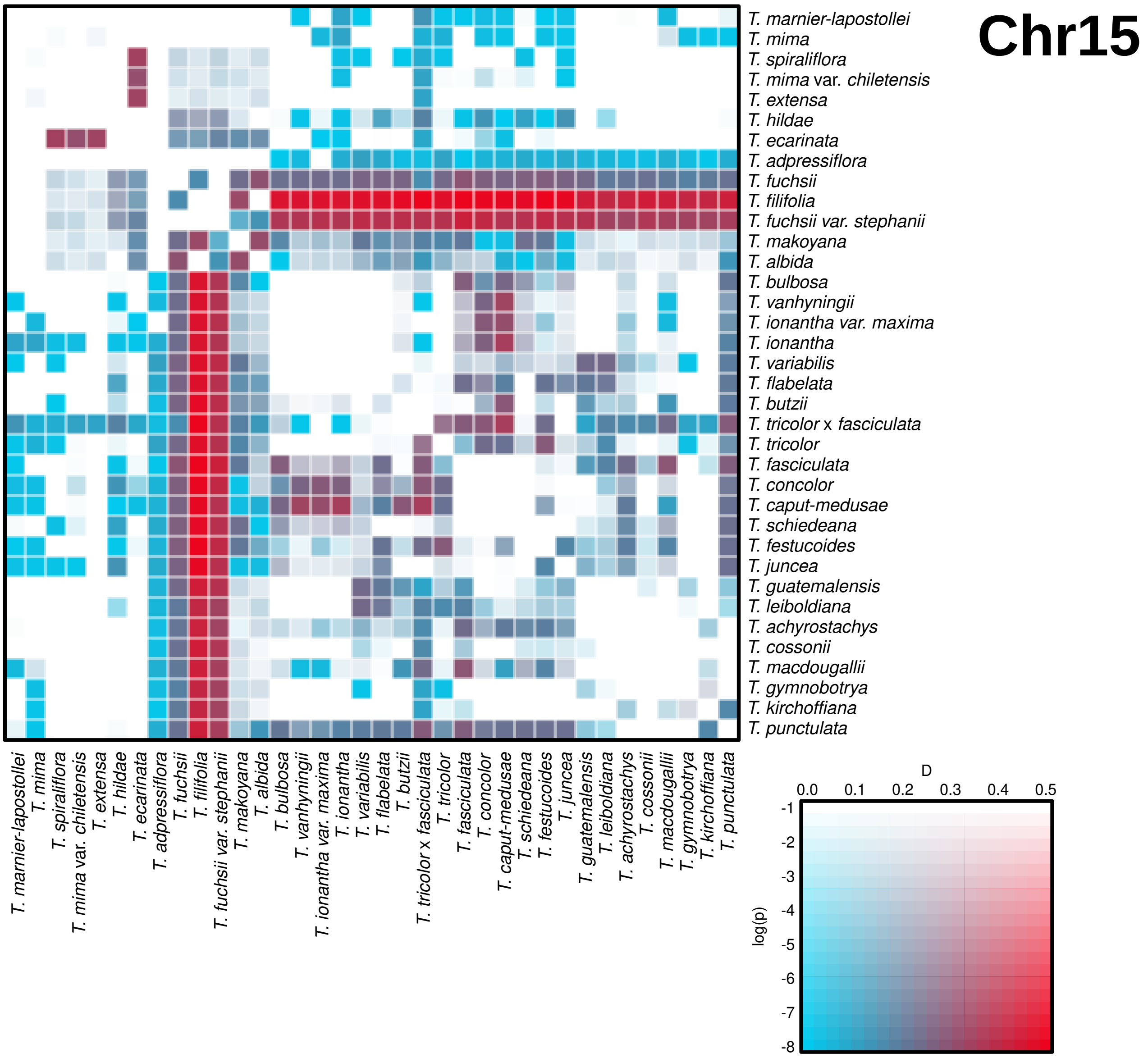
Chr15
T. marnier-lapostollei
T. mima
T. spiraliflora
T. mima var. chiletensis
T. extensa
T. hildae
T. ecarinata
T. adpressiflora
T. fuchsii
T. filifolia
T. fuchsii var. stephanii
T. makoyana
T. albida
T. bulbosa
T. vanhyningii
T. ionantha var. maxima
T. ionantha
T. variabilis
T. flabelata
T. butzii
T. tricolor x fasciculata
T. tricolor
T. fasciculata
T. concolor
T. caput-medusae
T. schiedeana
T. festucoides
T. juncea
T. guatemalensis
T. leiboldiana
T. achyrostachys
T. cossonii
T. macdougallii
T. gymnobotrya
T. kirchoffiana
T. punctulata
D
0.0
0.1
0.2
0.3
0.4
0.5
log(p)
-1
-2
-3
-4
-5
-6
-7
-8

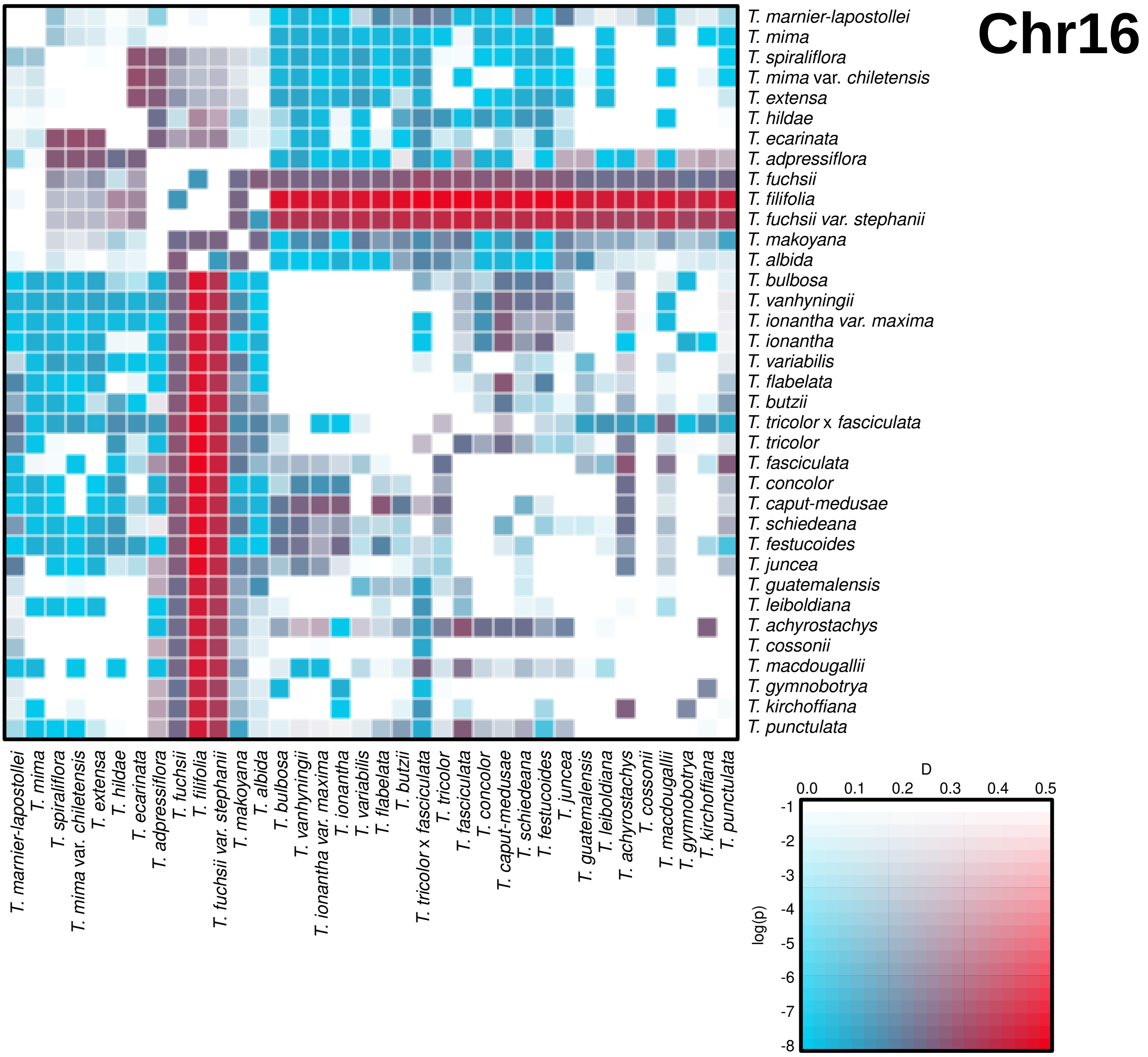
Chr16
T. marnier-lapostollei
T. mima
T. spiraliflora
T. mima var. chiletensis
T. extensa
T. hildae
T. ecarinata
T. adpressiflora
T. fuchsii
T. filifolia
T. fuchsii var. stephanii
T. makoyana
T. albida
T. bulbosa
T. vanhyningii
T. ionantha var. maxima
T. ionantha
T. variabilis
T. flabelata
T. butzii
T. tricolor x fasciculata
T. tricolor
T. fasciculata
T. concolor
T. caput-medusae
T. schiedeana
T. festucoides
T. juncea
T. guatemalensis
T. leiboldiana
T. achyrostachys
T. cossonii
T. macdougallii
T. gymnobotrya
T. kirchoffiana
T. punctulata
D
0.0
0.1
0.2
0.3
0.4
0.5
log(p)
-1
-2
-3
-4
-5
-6
-7
-8

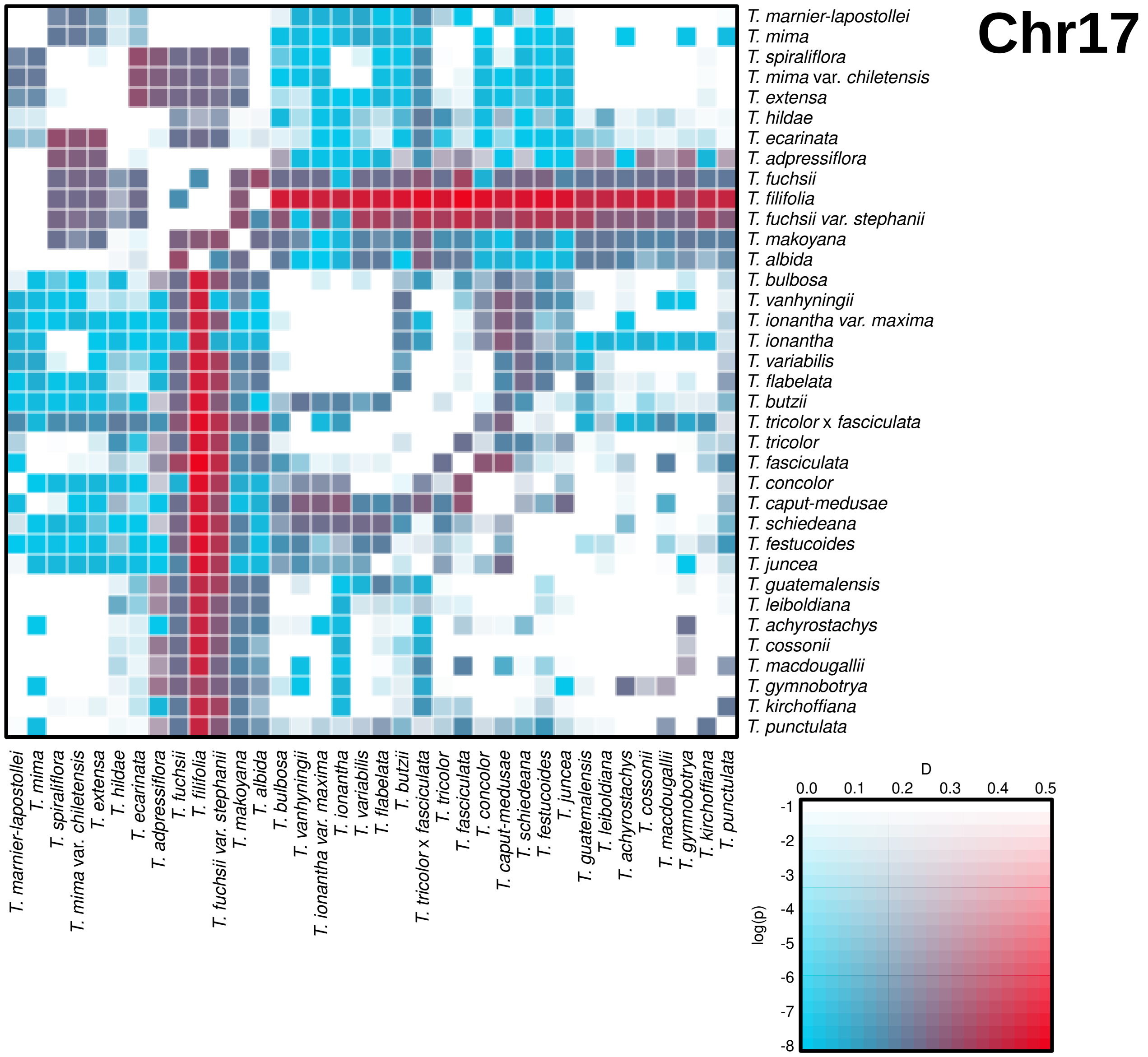
Chr17
T. marnier-lapostollei
T. mima
T. spiraliflora
T. mima var. chiletensis
T. extensa
T. hildae
T. ecarinata
T. adpressiflora
T. fuchsii
T. filifolia
T. fuchsii var. stephanii
T. makoyana
T. albida
T. bulbosa
T. vanhyningii
T. ionantha var. maxima
T. ionantha
T. variabilis
T. flabelata
T. butzii
T. tricolor x fasciculata
T. tricolor
T. fasciculata
T. concolor
T. caput-medusae
T. schiedeana
T. festucoides
T. juncea
T. guatemalensis
T. leiboldiana
T. achyrostachys
T. cossonii
T. macdougallii
T. gymnobotrya
T. kirchoffiana
T. punctulata
D
0.0 0.1 0.2 0.3 0.4 0.5
log(p)
-1 -2 -3 -4 -5 -6 -7 -8

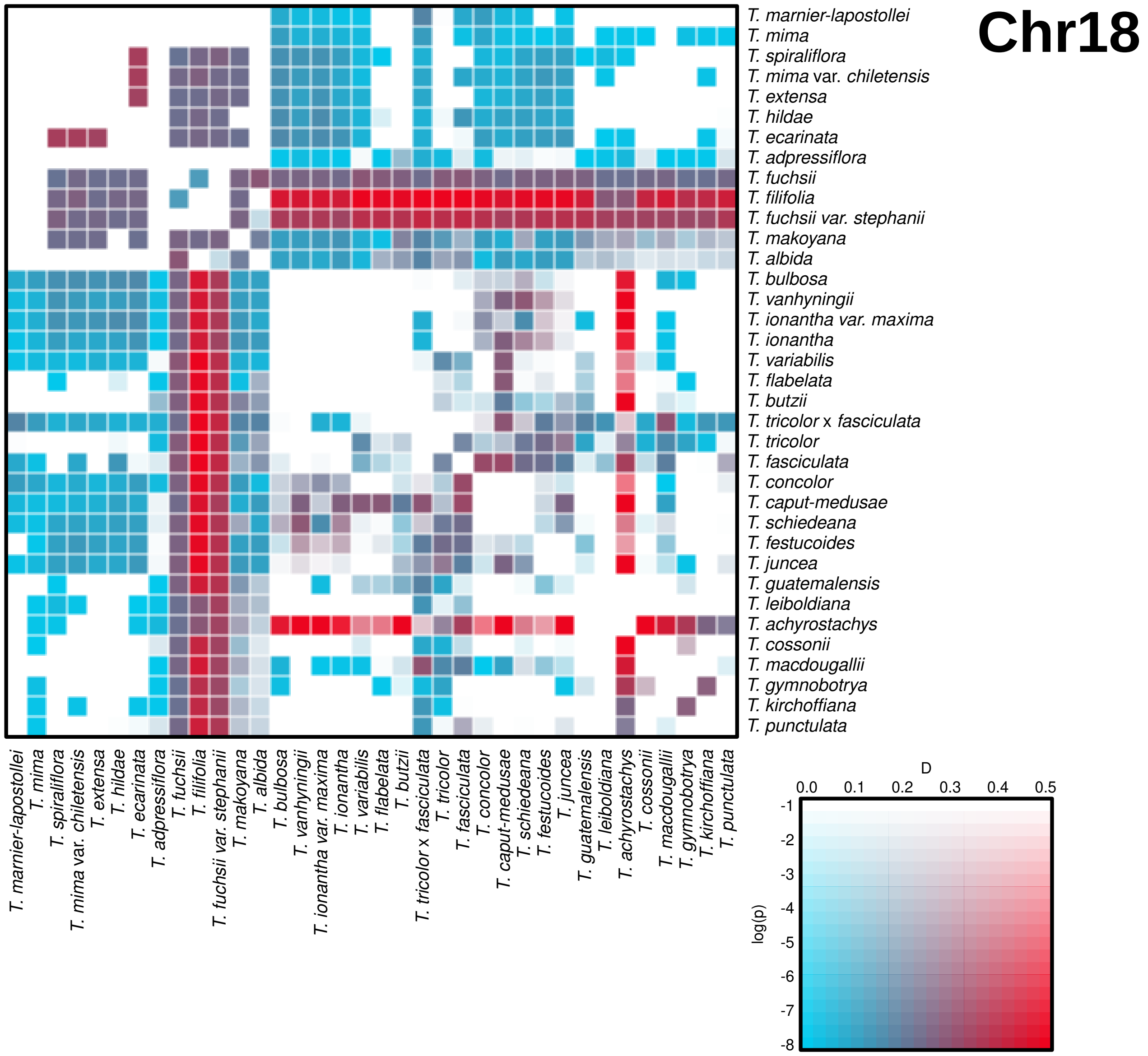

Chr18
T. marnier-lapostollei
T. mima
T. spiraliflora
T. mima var. chiletensis
T. extensa
T. hildae
T. ecarinata
T. adpressiflora
T. fuchsii
T. filifolia
T. fuchsii var. stephanii
T. makoyana
T. albida
T. bulbosa
T. vanhyningii
T. ionantha var. maxima
T. ionantha
T. variabilis
T. flabelata
T. butzii
T. tricolor x fasciculata
T. tricolor
T. fasciculata
T. concolor
T. caput-medusae
T. schiedeana
T. festucoides
T. juncea
T. guatemalensis
T. leiboldiana
T. achyrostachys
T. cossonii
T. macdougallii
T. gymnobotrya
T. kirchoffiana
T. punctulata
D
0.0
0.1
0.2
0.3
0.4
0.5
log(p)
-1
-2
-3
-4
-5
-6
-7
-8

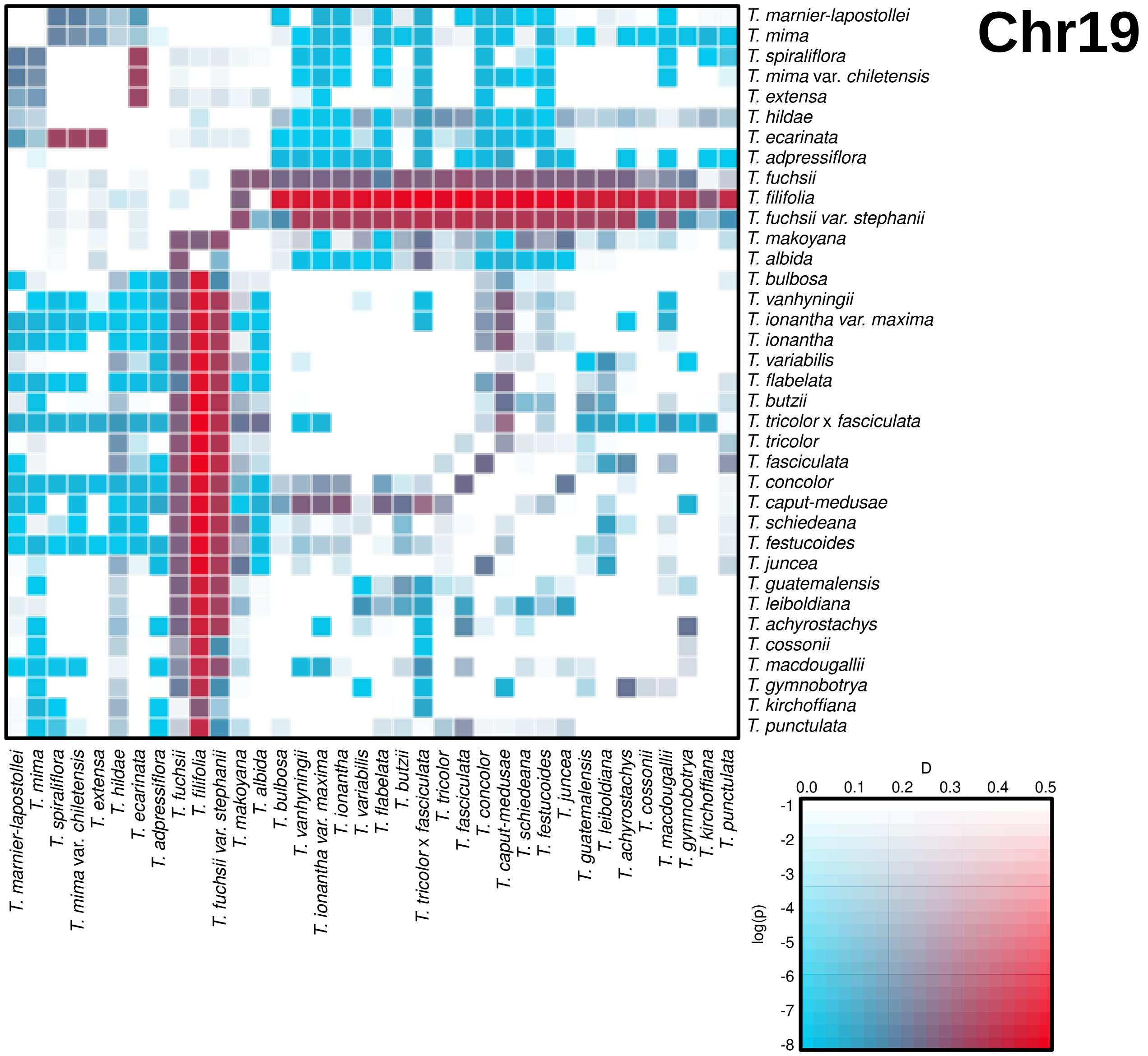
Chr19
T. marnier-lapostollei
T. mima
T. spiraliflora
T. mima var. chiletensis
T. extensa
T. hildae
T. ecarinata
T. adpressiflora
T. fuchsii
T. filifolia
T. fuchsii var. stephanii
T. makoyana
T. albida
T. bulbosa
T. vanhyningii
T. ionantha var. maxima
T. ionantha
T. variabilis
T. flabelata
T. butzii
T. tricolor x fasciculata
T. tricolor
T. fasciculata
T. concolor
T. caput-medusae
T. schiedeana
T. festucoides
T. juncea
T. guatemalensis
T. leiboldiana
T. achyrostachys
T. cossonii
T. macdougallii
T. gymnobotrya
T. kirchoffiana
T. punctulata
D
0.0
0.1
0.2
0.3
0.4
0.5
log(p)
-1
-2
-3
-4
-5
-6
-7
-8

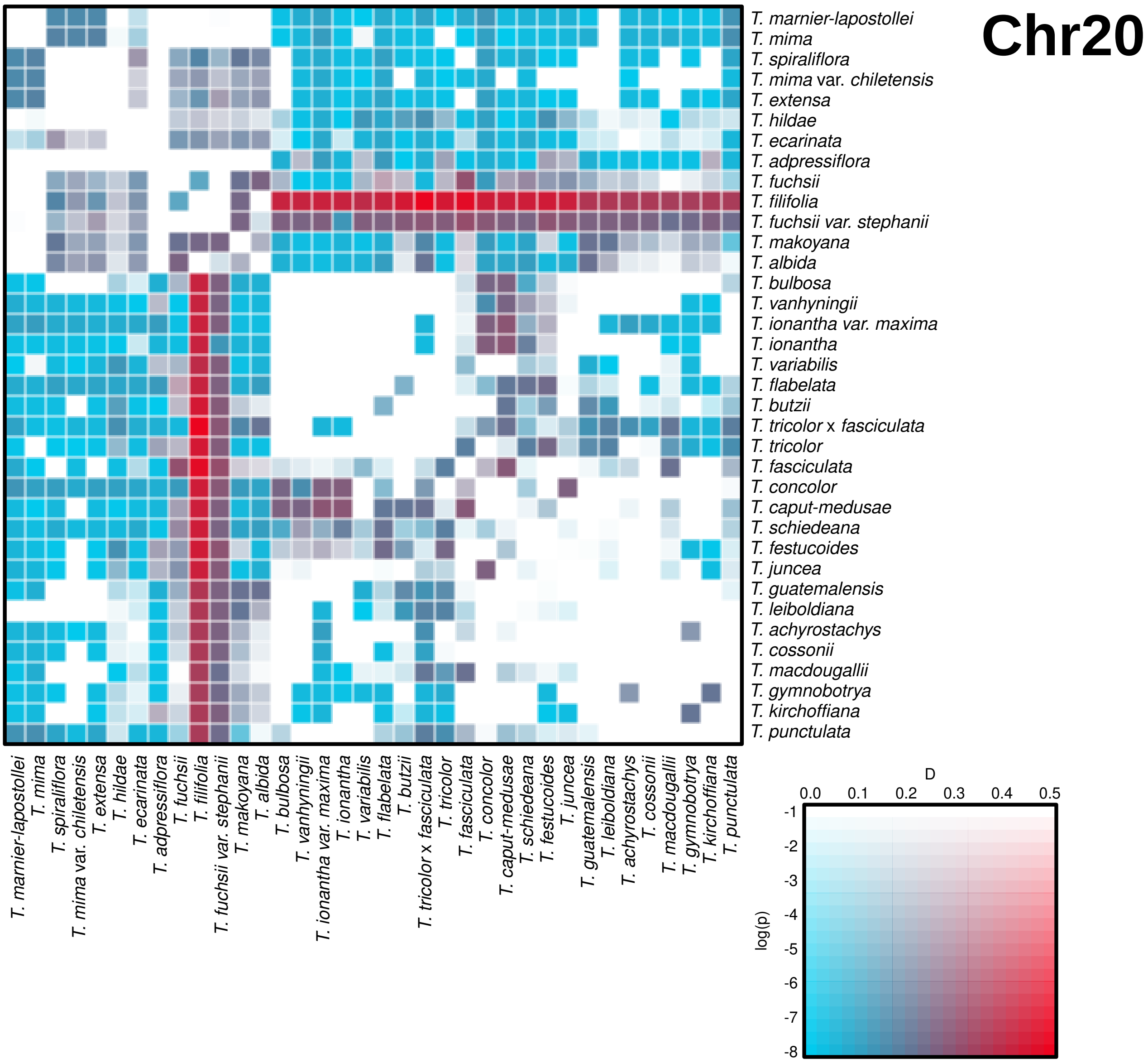
Chr20
T. marnier-lapostollei
T. mima
T. spiraliflora
T. mima var. chiletensis
T. extensa
T. hildae
T. ecarinata
T. adpressiflora
T. fuchsii
T. filifolia
T. fuchsii var. stephanii
T. makoyana
T. albida
T. bulbosa
T. vanhyningii
T. ionantha var. maxima
T. ionantha
T. variabilis
T. flabelata
T. butzii
T. tricolor x fasciculata
T. tricolor
T. fasciculata
T. concolor
T. caput-medusae
T. schiedeana
T. festucoides
T. juncea
T. guatemalensis
T. leiboldiana
T. achyrostachys
T. cossonii
T. macdougallii
T. gymnobotrya
T. kirchoffiana
T. punctulata
D
0.0
0.1
0.2
0.3
0.4
0.5
log(p)
-1
-2
-3
-4
-5
-6
-7
-8

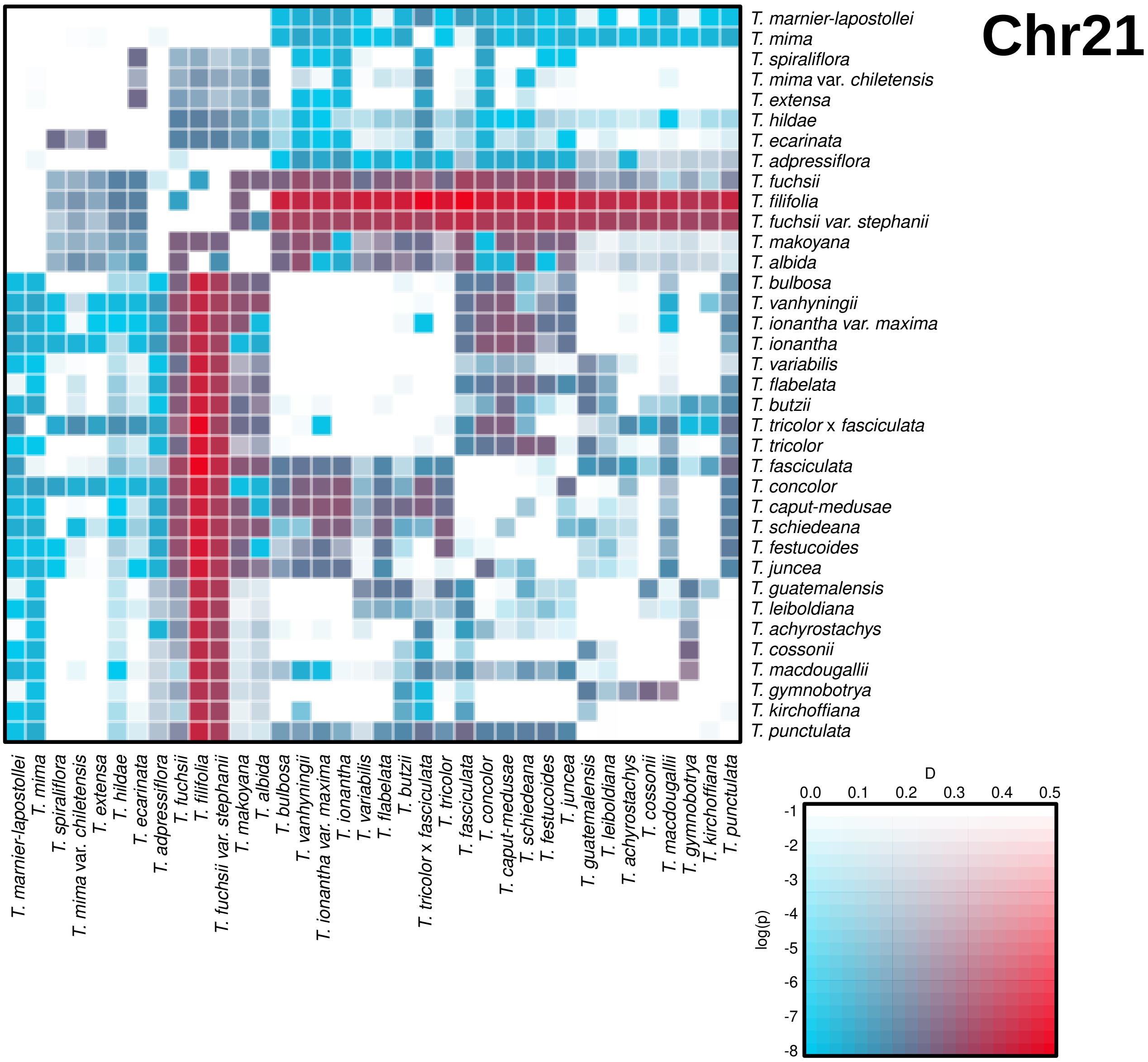
Chr21
T. marnier-lapostollei
T. mima
T. spiraliflora
T. mima var. chiletensis
T. extensa
T. hildae
T. ecarinata
T. adpressiflora
T. fuchsii
T. filifolia
T. fuchsii var. stephanii
T. makoyana
T. albida
T. bulbosa
T. vanhyningii
T. ionantha var. maxima
T. ionantha
T. variabilis
T. flabelata
T. butzii
T. tricolor x fasciculata
T. tricolor
T. fasciculata
T. concolor
T. caput-medusae
T. schiedeana
T. festucoides
T. juncea
T. guatemalensis
T. leiboldiana
T. achyrostachys
T. cossonii
T. macdougallii
T. gymnobotrya
T. kirchoffiana
T. punctulata
D
0.0
0.1
0.2
0.3
0.4
0.5
log(p)
-1
-2
-3
-4
-5
-6
-7
-8

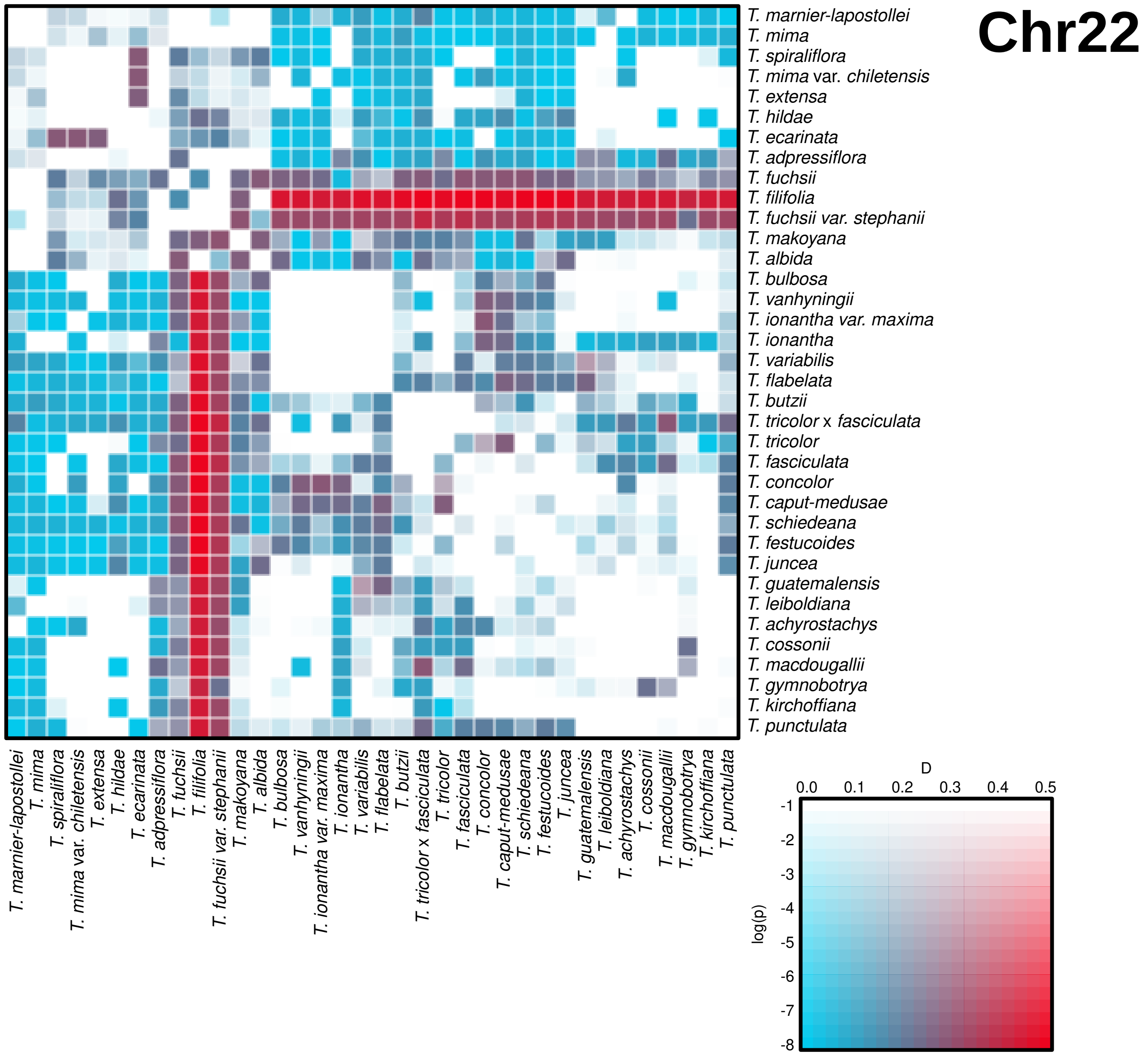
Chr22
T. marnier-lapostollei
T. mima
T. spiraliflora
T. mima var. chiletensis
T. extensa
T. hildae
T. ecarinata
T. adpressiflora
T. fuchsii
T. filifolia
T. fuchsii var. stephanii
T. makoyana
T. albida
T. bulbosa
T. vanhyningii
T. ionantha var. maxima
T. ionantha
T. variabilis
T. flabelata
T. butzii
T. tricolor x fasciculata
T. tricolor
T. fasciculata
T. concolor
T. caput-medusae
T. schiedeana
T. festucoides
T. juncea
T. guatemalensis
T. leiboldiana
T. achyrostachys
T. cossonii
T. macdougallii
T. gymnobotrya
T. kirchoffiana
T. punctulata
D
0.0
0.1
0.2
0.3
0.4
0.5
log(p)
-1
-2
-3
-4
-5
-6
-7
-8

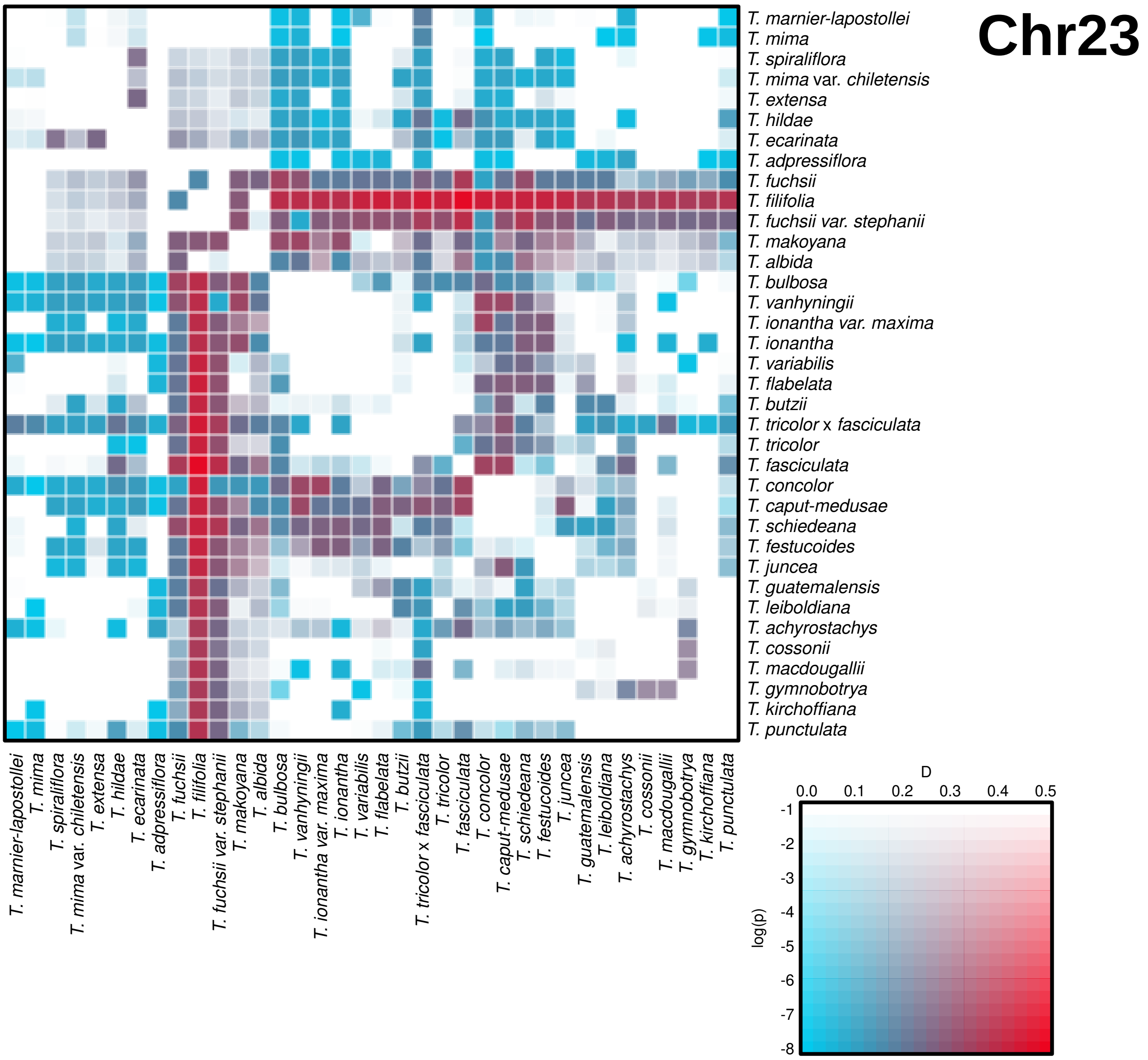
Chr23
T. marnier-lapostollei
T. mima
T. spiraliflora
T. mima var. chiletensis
T. extensa
T. hildae
T. ecarinata
T. adpressiflora
T. fuchsii
T. filifolia
T. fuchsii var. stephanii
T. makoyana
T. albida
T. bulbosa
T. vanhyningii
T. ionantha var. maxima
T. ionantha
T. variabilis
T. flabelata
T. butzii
T. tricolor x fasciculata
T. tricolor
T. fasciculata
T. concolor
T. caput-medusae
T. schiedeana
T. festucoides
T. juncea
T. guatemalensis
T. leiboldiana
T. achyrostachys
T. cossonii
T. macdougallii
T. gymnobotrya
T. kirchoffiana
T. punctulata
D
0.0
0.1
0.2
0.3
0.4
0.5
log(p)
-1
-2
-3
-4
-5
-6
-7
-8

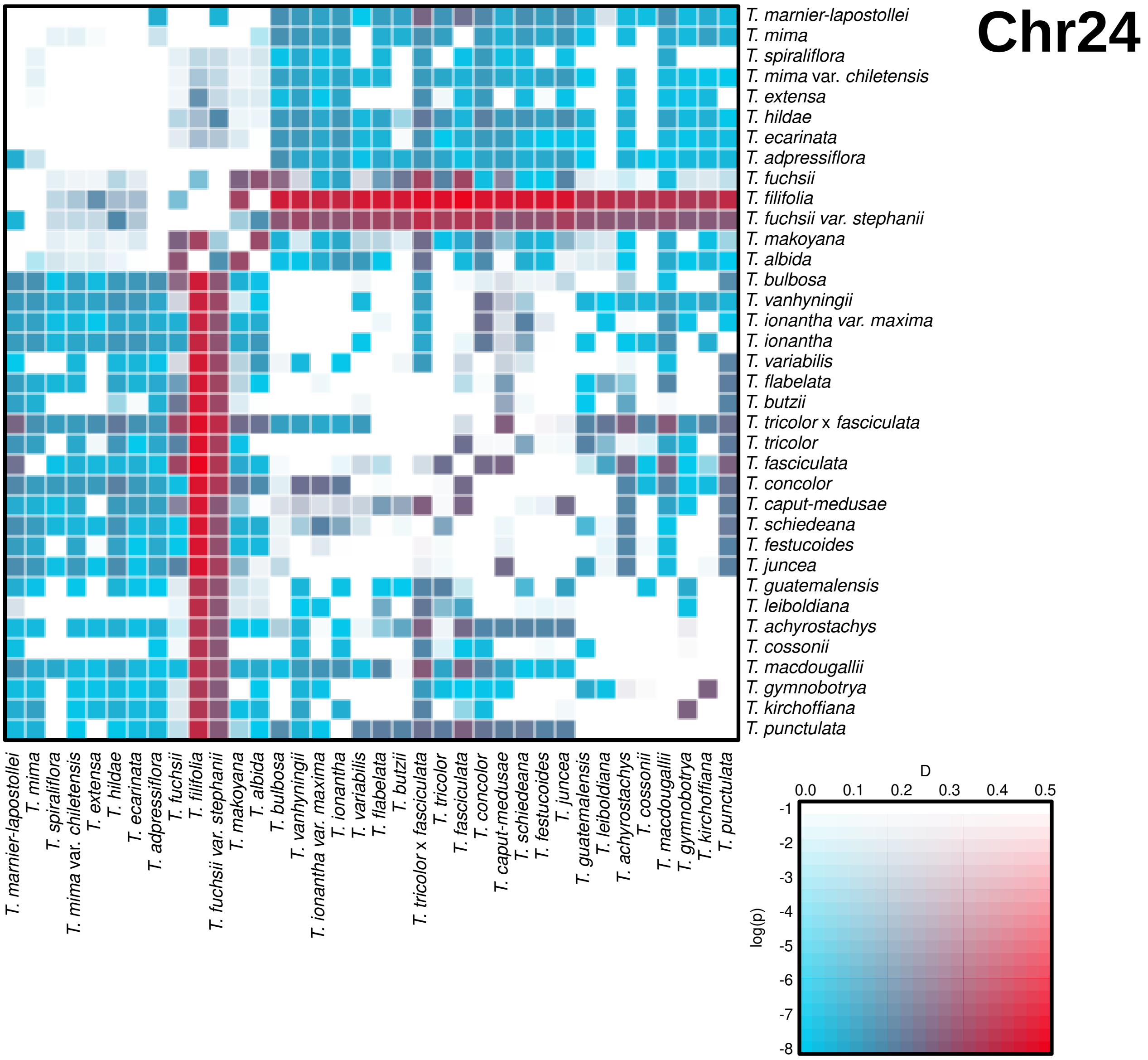
Chr24
T. marnier-lapostollei
T. mima
T. spiraliflora
T. mima var. chiletensis
T. extensa
T. hildae
T. ecarinata
T. adpressiflora
T. fuchsii
T. filifolia
T. fuchsii var. stephanii
T. makoyana
T. albida
T. bulbosa
T. vanhyningii
T. ionantha var. maxima
T. ionantha
T. variabilis
T. flabelata
T. butzii
T. tricolor x fasciculata
T. tricolor
T. fasciculata
T. concolor
T. caput-medusae
T. schiedeana
T. festucoides
T. juncea
T. guatemalensis
T. leiboldiana
T. achyrostachys
T. cossonii
T. macdougallii
T. gymnobotrya
T. kirchoffiana
T. punctulata
D
0.0
0.1
0.2
0.3
0.4
0.5
log(p)
-1
-2
-3
-4
-5
-6
-7
-8

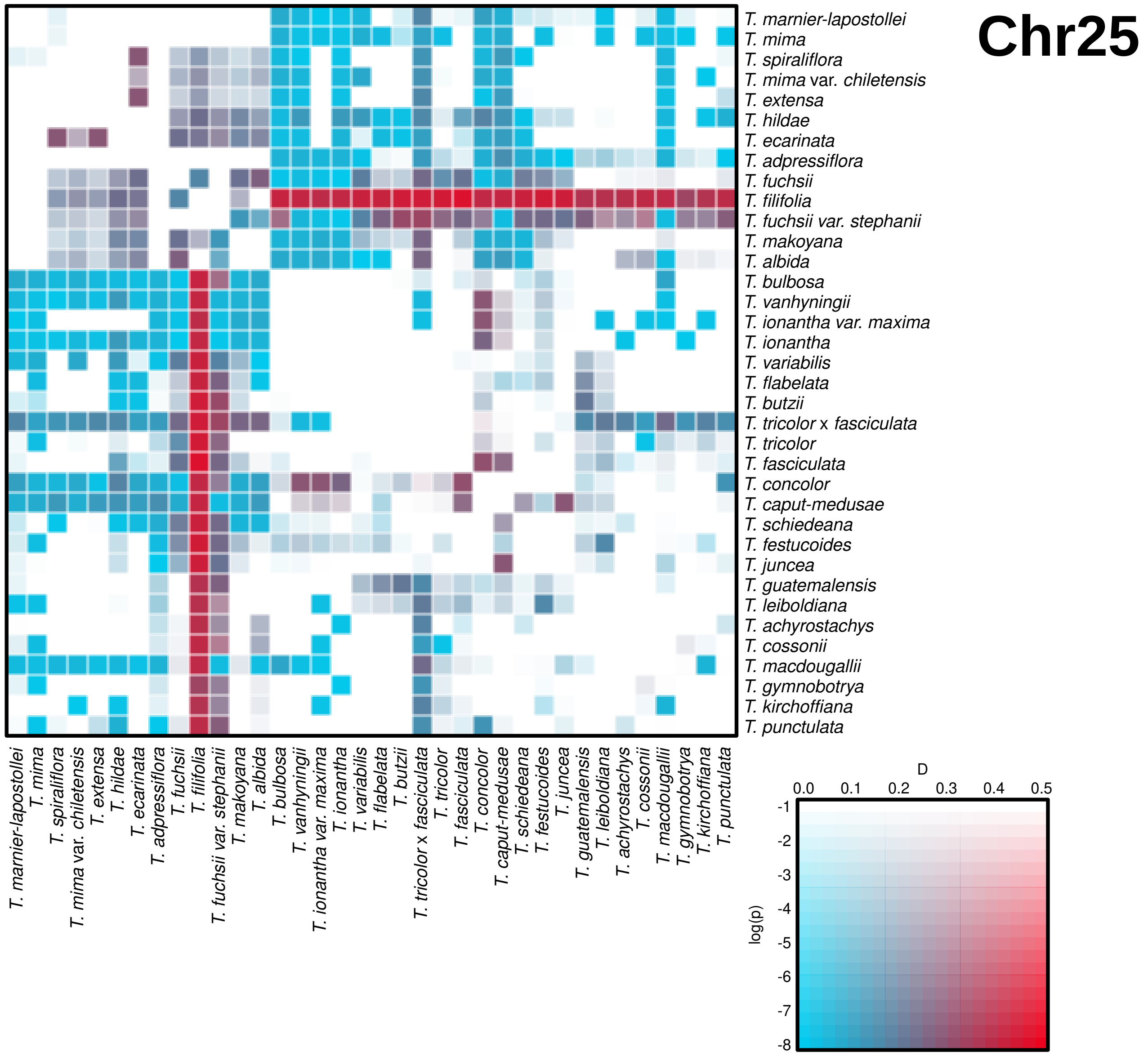
Chr25
T. marnier-lapostollei
T. mima
T. spiraliflora
T. mima var. chiletensis
T. extensa
T. hildae
T. ecarinata
T. adpressiflora
T. fuchsii
T. filifolia
T. fuchsii var. stephanii
T. makoyana
T. albida
T. bulbosa
T. vanhyningii
T. ionantha var. maxima
T. ionantha
T. variabilis
T. flabelata
T. butzii
T. tricolor x fasciculata
T. tricolor
T. fasciculata
T. concolor
T. caput-medusae
T. schiedeana
T. festucoides
T. juncea
T. guatemalensis
T. leiboldiana
T. achyrostachys
T. cossonii
T. macdougallii
T. gymnobotrya
T. kirchoffiana
T. punctulata
D
0.0
0.1
0.2
0.3
0.4
0.5
log(p)
-1
-2
-3
-4
-5
-6
-7
-8
