## supporting file 3 for "The explosive radiation of the Neotropical *Tillandsia* subgenus *Tillandsia* (Bromeliaceae) has been accompanied by pervasive hybridization"

### Supporting file 3 – reanalysis with different minor allele values

**General notes**

The filtering threshold chosen for this work was MAF < 0.045; corresponding to a minor allele present in at least 6 chromosomal complements in a data-set of 69 individuals (see main text). Any filter for minor allele frequency may bias branch lengths in phylogenetic analysis. To examine the possibility MAF filtering biasing the results, we repeated the analysis with different thresholds of filtering for minor alleles. To increase clarity, we use an explicit value of MAC (minor allele count) with values between 0-5.

**SNP numbers**

| MAC filter | SNP numbers |
|---|---|
| 0 | 12,170,779 |
| 1 | 7,954,068 |
| 2 | 4,970,792 |
| 3 | 3,869,478 |
| 4 | 2,600,839 |
| 5 | 2,256,452 |

**Maximum likelihood trees**

for simplification, maximum likelihood trees were ran on a concatenated SNP matrix – otherwise as described in the main text, using iqtree2 with model inference. Bootstrap values were not calculated. Clade colors correspond to main text.
In the results below, differences in branch length can be observed. As expected, long branches are diminished as minor alleles are filtered. This mostly affects single species, rather than whole clades – notably, species like *T. makoyana* and *T. achyrostachys*. The tree topology is not affected.

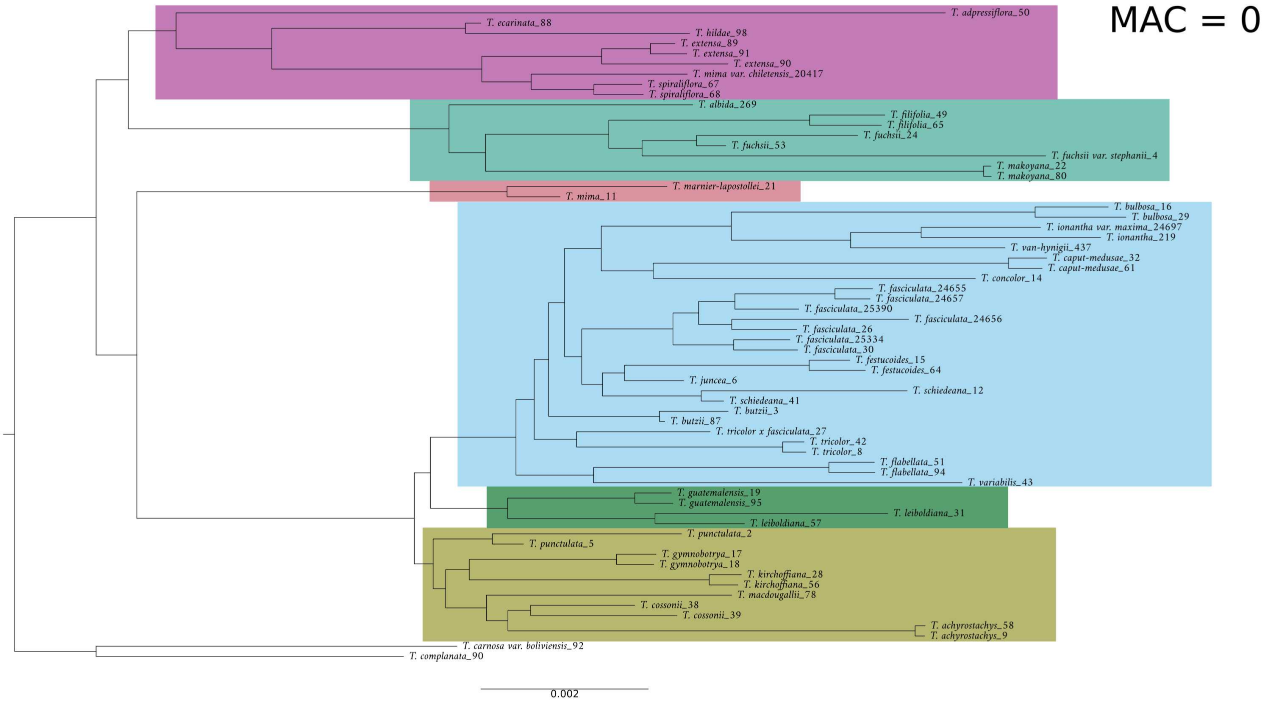
MAC = 0
T. adpressiflora_50
T. ecarinata_88
T. hildae_98
T. extensa_89
T. extensa_91
T. extensa_90
T. mima var. chiletensis_20417
T. spiraliflora_67
T. spiraliflora_68
T. albida_269
T. filifolia_49
T. filifolia_65
T. fuchsii_24
T. fuchsii_53
T. fuchsii var. stephanii_4
T. makoyana_22
T. makoyana_80
T. marnier-lapostollei_21
T. mima_11
T. bulbosa_16
T. bulbosa_29
T. ionantha var. maxima_24697
T. ionantha_219
T. van-hynigii_437
T. caput-medusae_32
T. caput-medusae_61
T. concolor_14
T. fasciculata_24655
T. fasciculata_24657
T. fasciculata_25390
T. fasciculata_24656
T. fasciculata_26
T. fasciculata_25334
T. fasciculata_30
T. festucoides_15
T. festucoides_64
T. juncea_6
T. schiedeana_12
T. schiedeana_41
T. butzii_3
T. butzii_87
T. tricolor x fasciculata_27
T. tricolor_42
T. tricolor_8
T. flabellata_51
T. flabellata_94
T. variabilis_43
T. guatemalensis_19
T. guatemalensis_95
T. leiboldiana_31
T. leiboldiana_57
T. punctulata_2
T. punctulata_5
T. gymnobotrya_17
T. gymnobotrya_18
T. kirchoffiana_28
T. kirchoffiana_56
T. macdougallii_78
T. cossonii_38
T. cossonii_39
T. achyrostachys_58
T. achyrostachys_9
T. carnosa var. boliviensis_92
T. complanata_90
0.002

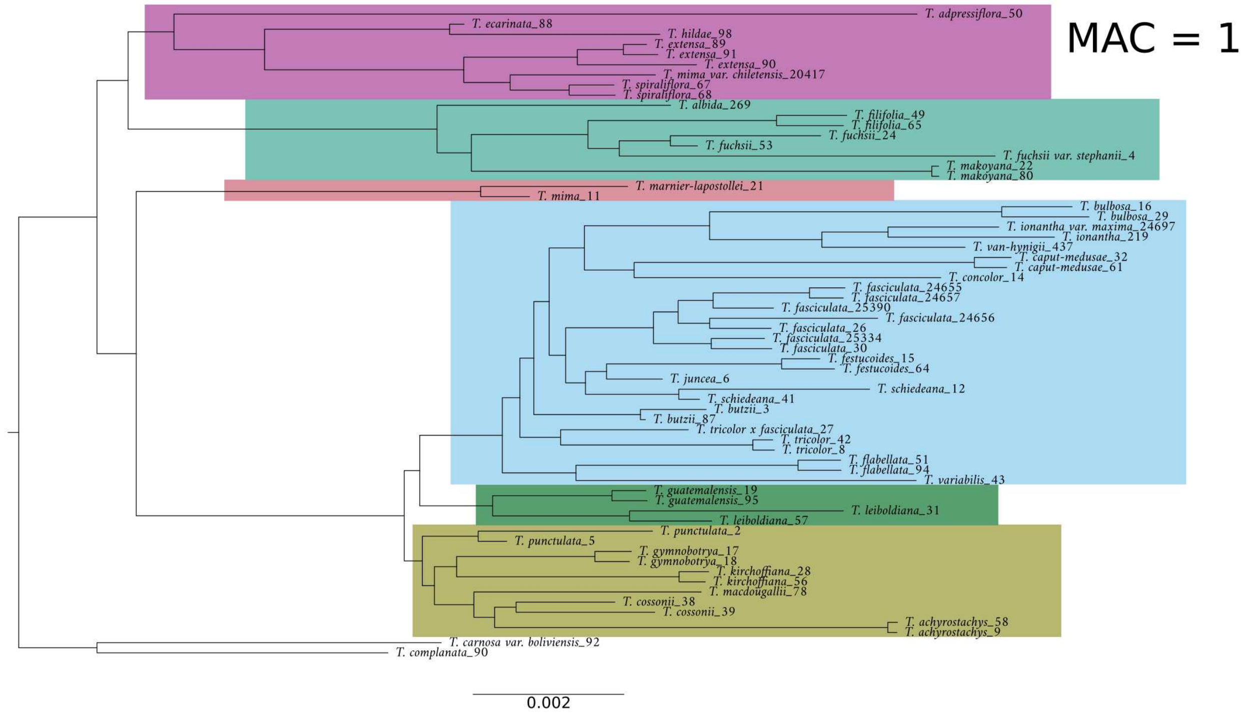
MAC = 1
T. adpressiflora_50
T. ecarinata_88
T. hildae_98
T. extensa_89
T. extensa_91
T. extensa_90
T. mima var. chiletensis_20417
T. spiraliflora_67
T. spiraliflora_68
T. albida_269
T. filifolia_49
T. filifolia_65
T. fuchsii_24
T. fuchsii_53
T. fuchsii var. stephanii_4
T. makoyana_22
T. makoyana_80
T. marnier-lapostollei_21
T. mima_11
T. bulbosa_16
T. bulbosa_29
T. ionantha var. maxima_24697
T. ionantha_219
T. van-hynigii_437
T. caput-medusae_32
T. caput-medusae_61
T. concolor_14
T. fasciculata_24655
T. fasciculata_24657
T. fasciculata_25390
T. fasciculata_24656
T. fasciculata_26
T. fasciculata_25334
T. fasciculata_30
T. festucoides_15
T. festucoides_64
T. juncea_6
T. schiedeana_12
T. schiedeana_41
T. butzii_3
T. butzii_87
T. tricolor x fasciculata_27
T. tricolor_42
T. tricolor_8
T. flabellata_51
T. flabellata_94
T. variabilis_43
T. guatemalensis_19
T. guatemalensis_95
T. leiboldiana_31
T. leiboldiana_57
T. punctulata_2
T. punctulata_5
T. gymnobotrya_17
T. gymnobotrya_18
T. kirchoffiana_28
T. kirchoffiana_56
T. macdougallii_78
T. cossonii_38
T. cossonii_39
T. achyrostachys_58
T. achyrostachys_9
T. carnosa var. boliviensis_92
T. complanata_90
0.002

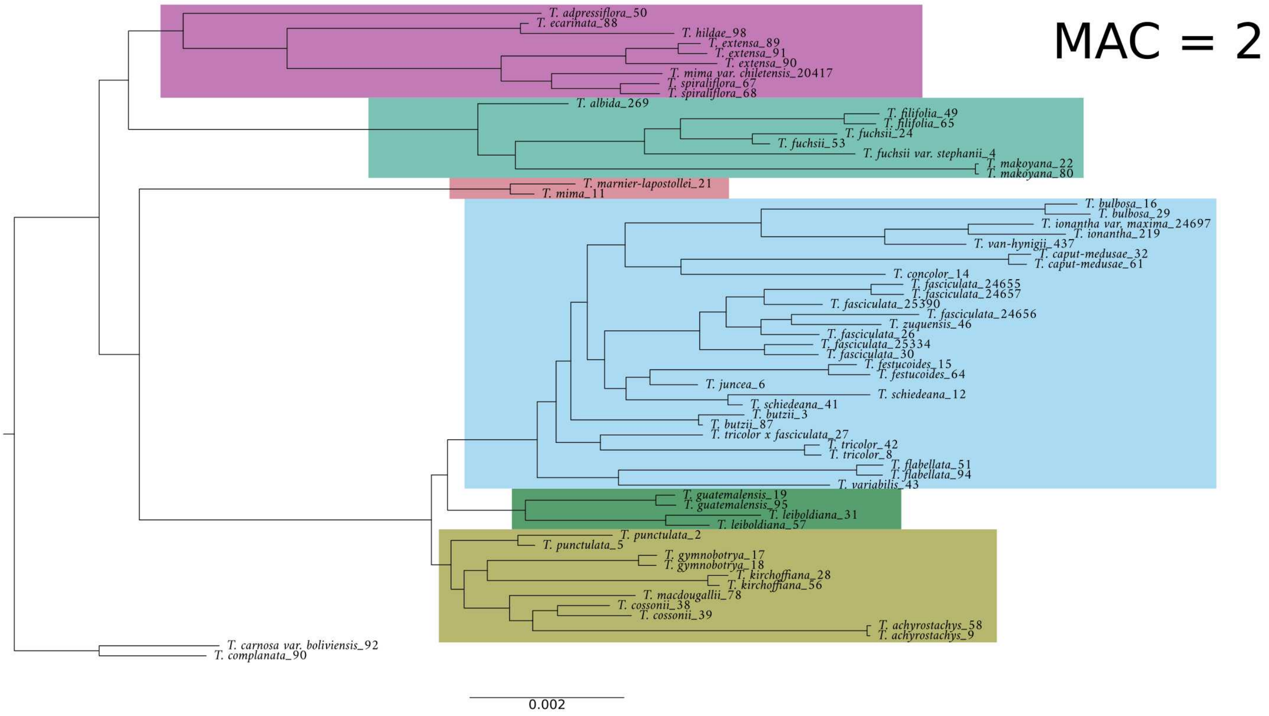
MAC = 2
T. adpressiflora_50
T. ecarinata_88
T. hildae_98
T. extensa_89
T. extensa_91
T. extensa_90
T. mima var. chiletensis_20417
T. spiraliflora_67
T. spiraliflora_68
T. albida_269
T. filifolia_49
T. filifolia_65
T. fuchsii_24
T. fuchsii_53
T. fuchsii var. stephanii_4
T. makoyana_22
T. makoyana_80
T. marnier-lapostollei_21
T. mima_11
T. bulbosa_16
T. bulbosa_29
T. ionantha var. maxima_24697
T. ionantha_219
T. van-hynigii_437
T. caput-medusae_32
T. caput-medusae_61
T. concolor_14
T. fasciculata_24655
T. fasciculata_24657
T. fasciculata_25390
T. fasciculata_24656
T. zuquensis_46
T. fasciculata_26
T. fasciculata_25334
T. fasciculata_30
T. festucoides_15
T. festucoides_64
T. juncea_6
T. schiedeana_12
T. schiedeana_41
T. butzii_3
T. butzii_87
T. tricolor x fasciculata_27
T. tricolor_42
T. tricolor_8
T. flabellata_51
T. flabellata_94
T. variabilis_43
T. guatemalensis_19
T. guatemalensis_95
T. leiboldiana_31
T. leiboldiana_57
T. punctulata_2
T. punctulata_5
T. gymnobotrya_17
T. gymnobotrya_18
T. kirchoffiana_28
T. kirchoffiana_56
T. macdougallii_78
T. cossonii_38
T. cossonii_39
T. achyrostachys_58
T. achyrostachys_9
T. carnosa var. boliviensis_92
T. complanata_90
0.002

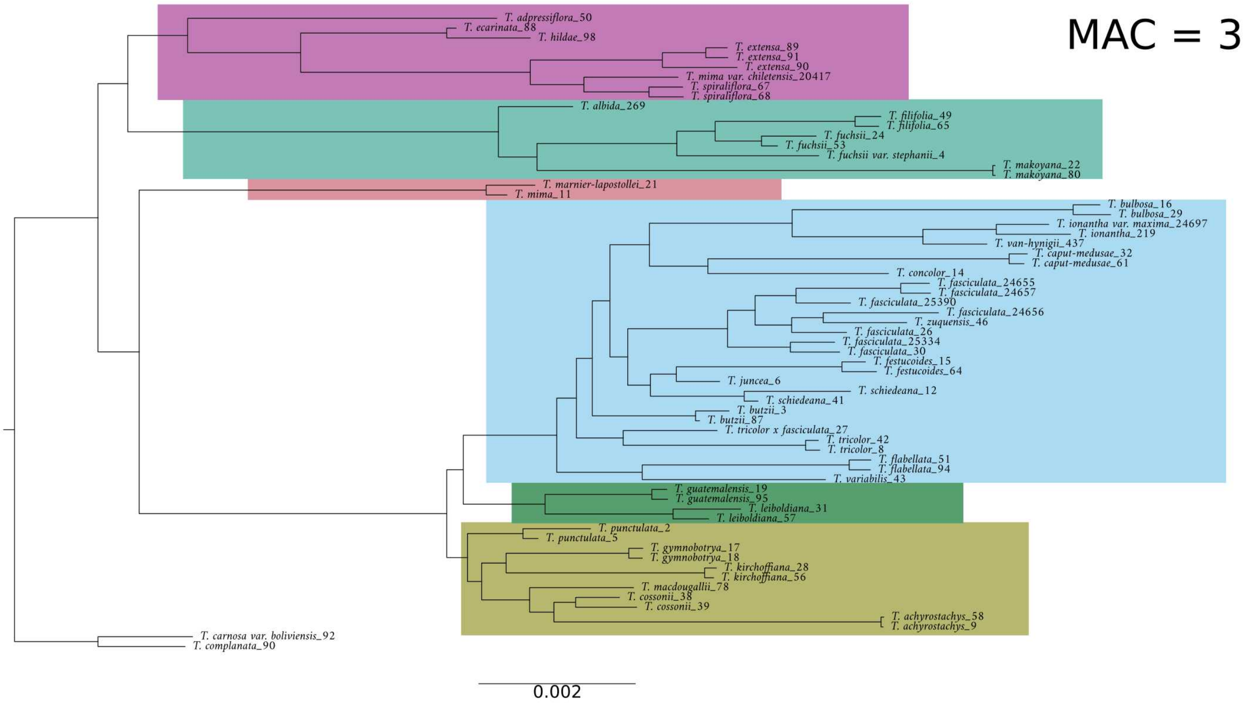
MAC = 3
T. adpressiflora_50
T. ecarinata_88
T. hildae_98
T. extensa_89
T. extensa_91
T. extensa_90
T. mima var. chiletensis_20417
T. spiraliflora_67
T. spiraliflora_68
T. albida_269
T. filifolia_49
T. filifolia_65
T. fuchsii_24
T. fuchsii_53
T. fuchsii var. stephanii_4
T. makoyana_22
T. makoyana_80
T. marnier-lapostollei_21
T. mima_11
T. bulbosa_16
T. bulbosa_29
T. ionantha var. maxima_24697
T. ionantha_219
T. van-hynigii_437
T. caput-medusae_32
T. caput-medusae_61
T. concolor_14
T. fasciculata_24655
T. fasciculata_24657
T. fasciculata_25390
T. fasciculata_24656
T. zuquensis_46
T. fasciculata_26
T. fasciculata_25334
T. fasciculata_30
T. festucoides_15
T. festucoides_64
T. juncea_6
T. schiedeana_12
T. schiedeana_41
T. butzii_3
T. butzii_87
T. tricolor x fasciculata_27
T. tricolor_42
T. tricolor_8
T. flabellata_51
T. flabellata_94
T. variabilis_43
T. guatemalensis_19
T. guatemalensis_95
T. leiboldiana_31
T. leiboldiana_57
T. punctulata_2
T. punctulata_5
T. gymnobotrya_17
T. gymnobotrya_18
T. kirchoffiana_28
T. kirchoffiana_56
T. macdougallii_78
T. cossonii_38
T. cossonii_39
T. achyrostachys_58
T. achyrostachys_9
T. carnosa var. boliviensis_92
T. complanata_90
0.002

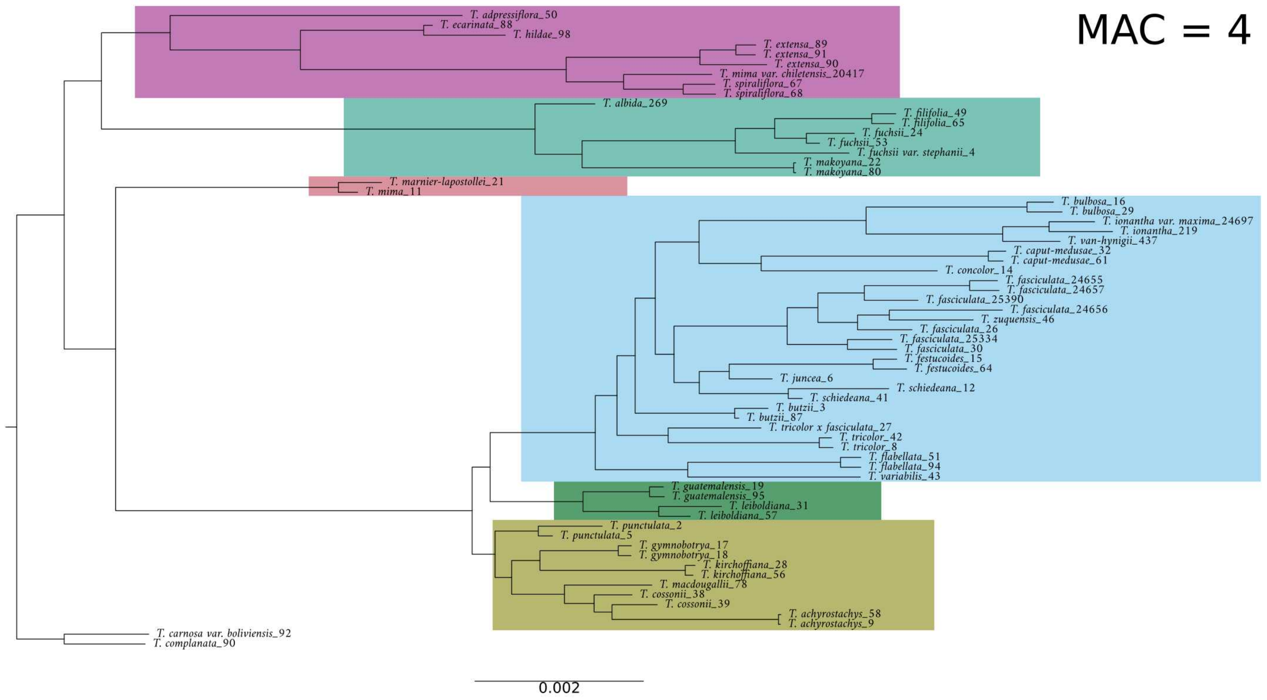
MAC = 4
T. adpressiflora_50
T. ecarinata_88
T. hildae_98
T. extensa_89
T. extensa_91
T. extensa_90
T. mima var. chiletensis_20417
T. spiraliflora_67
T. spiraliflora_68
T. albida_269
T. filifolia_49
T. filifolia_65
T. fuchsii_24
T. fuchsii_53
T. fuchsii var. stephanii_4
T. makoyana_22
T. makoyana_80
T. marnier-lapostollei_21
T. mima_11
T. bulbosa_16
T. bulbosa_29
T. ionantha var. maxima_24697
T. ionantha_219
T. van-hynigii_437
T. caput-medusae_32
T. caput-medusae_61
T. concolor_14
T. fasciculata_24655
T. fasciculata_24657
T. fasciculata_25390
T. fasciculata_24656
T. zuquensis_46
T. fasciculata_26
T. fasciculata_25334
T. fasciculata_30
T. festucoides_15
T. festucoides_64
T. juncea_6
T. schiedeana_12
T. schiedeana_41
T. butzii_3
T. butzii_87
T. tricolor x fasciculata_27
T. tricolor_42
T. tricolor_8
T. flabellata_51
T. flabellata_94
T. variabilis_43
T. guatemalensis_19
T. guatemalensis_95
T. leiboldiana_31
T. leiboldiana_57
T. punctulata_2
T. punctulata_5
T. gymnobotrya_17
T. gymnobotrya_18
T. kirchoffiana_28
T. kirchoffiana_56
T. macdougallii_78
T. cossonii_38
T. cossonii_39
T. achyrostachys_58
T. achyrostachys_9
T. carnosa var. boliviensis_92
T. complanata_90
0.002

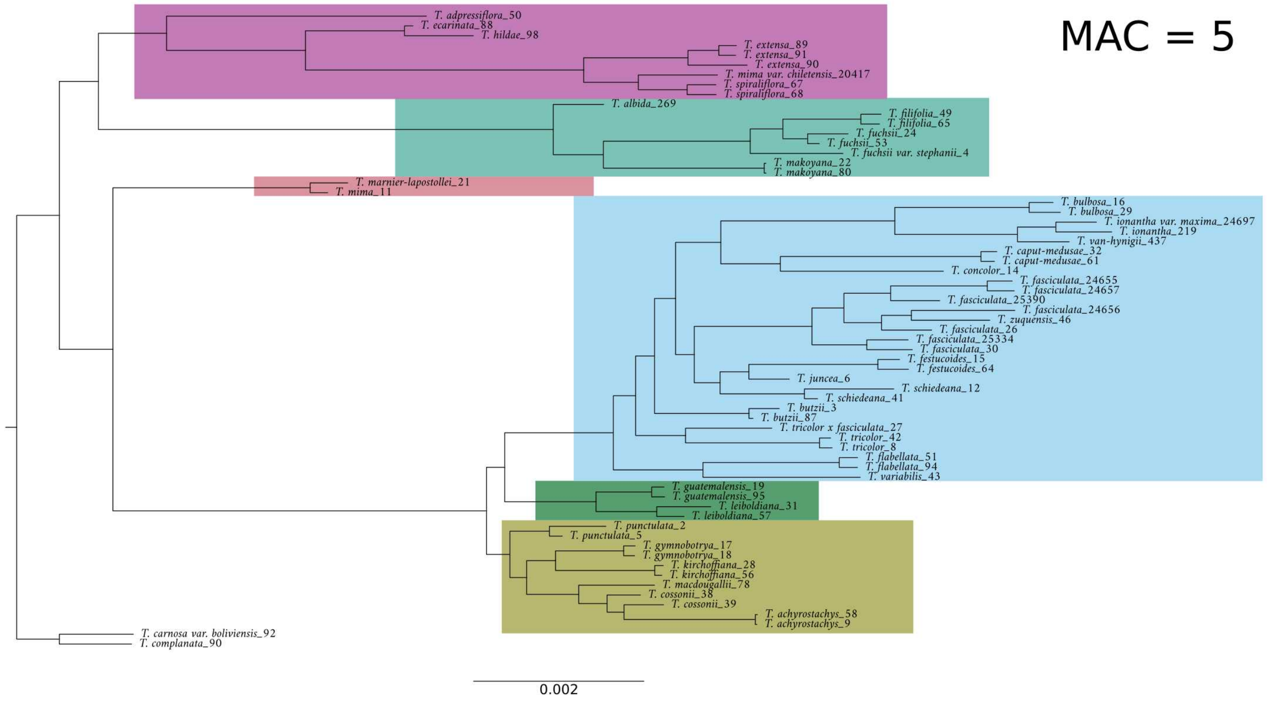
MAC = 5
T. adpressiflora_50
T. ecarinata_88
T. hildae_98
T. extensa_89
T. extensa_91
T. extensa_90
T. mima var. chiletensis_20417
T. spiraliflora_67
T. spiraliflora_68
T. albida_269
T. filifolia_49
T. filifolia_65
T. fuchsii_24
T. fuchsii_53
T. fuchsii var. stephanii_4
T. makoyana_22
T. makoyana_80
T. marnier-lapostollei_21
T. mima_11
T. bulbosa_16
T. bulbosa_29
T. ionantha var. maxima_24697
T. ionantha_219
T. van-hynigii_437
T. caput-medusae_32
T. caput-medusae_61
T. concolor_14
T. fasciculata_24655
T. fasciculata_24657
T. fasciculata_25390
T. fasciculata_24656
T. zuquensis_46
T. fasciculata_26
T. fasciculata_25334
T. fasciculata_30
T. festucoides_15
T. festucoides_64
T. juncea_6
T. schiedeana_12
T. schiedeana_41
T. butzii_3
T. butzii_87
T. tricolor x fasciculata_27
T. tricolor_42
T. tricolor_8
T. flabellata_51
T. flabellata_94
T. variabilis_43
T. guatemalensis_19
T. guatemalensis_95
T. leiboldiana_31
T. leiboldiana_57
T. punctulata_2
T. punctulata_5
T. gymnobotrya_17
T. gymnobotrya_18
T. kirchoffiana_28
T. kirchoffiana_56
T. macdougallii_78
T. cossonii_38
T. cossonii_39
T. achyrostachys_58
T. achyrostachys_9
T. carnosa var. boliviensis_92
T. complanata_90
0.002

**Astral trees**

All species trees were generated as described in the main text. Briefly, ML trees were generated on non-overlapping windows of 10kb. Windows with fewer than 40 SNPs were excluded. nodes with a bootstrap support below ten were collapsed. A species tree was inferred with ASTRAL-III. Below, the pies as the nodes represent quartet support for each topology, with blue representing support for the main topology and green for the second topology.

The relationships between clades are consistent, save for unusual placement of the *T. mima* clade at MAC=4. *T. mima* placement differed between the ML trees (above) and all species tree, hence cannot confidently be placed in this data-set. Few relationships were affected by the MAC filtering, like between species in the *T. punctulata* clade and specifically the monphyly of *T. punctulata*. Values of quartet support, reflecting discordance, differed minimally.

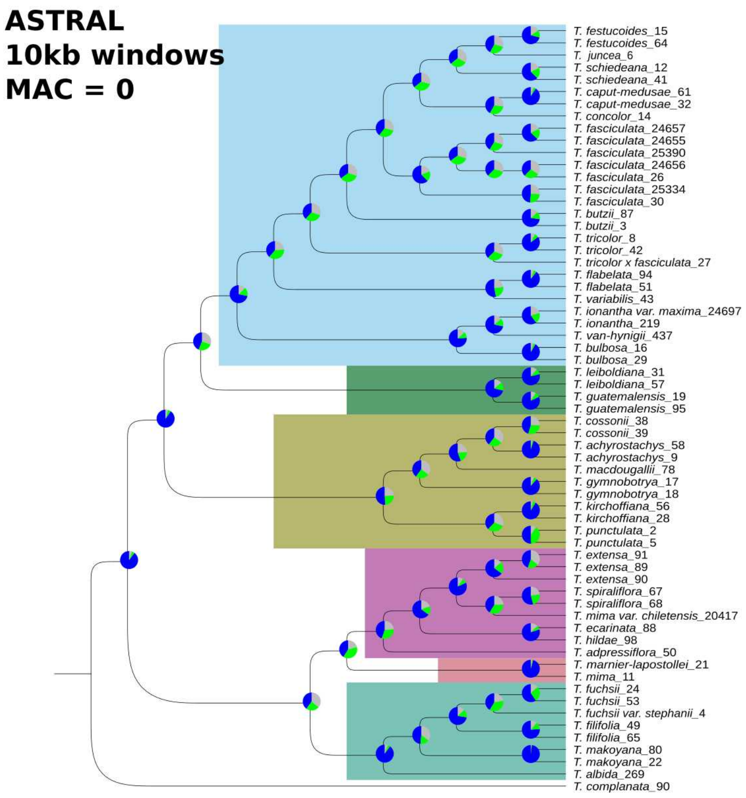
ASTRAL
10kb windows
MAC = 0
T. festucoides_15
T. festucoides_64
T. juncea_6
T. schiedeana_12
T. schiedeana_41
T. caput-medusae_61
T. caput-medusae_32
T. concolor_14
T. fasciculata_24657
T. fasciculata_24655
T. fasciculata_25390
T. fasciculata_24656
T. fasciculata_26
T. fasciculata_25334
T. fasciculata_30
T. butzii_87
T. butzii_3
T. tricolor_8
T. tricolor_42
T. tricolor x fasciculata_27
T. flabelata_94
T. flabelata_51
T. variabilis_43
T. ionantha var. maxima_24697
T. ionantha_219
T. van-hynigii_437
T. bulbosa_16
T. bulbosa_29
T. leiboldiana_31
T. leiboldiana_57
T. guatemalensis_19
T. guatemalensis_95
T. cossonii_38
T. cossonii_39
T. achyrostachys_58
T. achyrostachys_9
T. macdougallii_78
T. gymnobotrya_17
T. gymnobotrya_18
T. kirchoffiana_56
T. kirchoffiana_28
T. punctulata_2
T. punctulata_5
T. extensa_91
T. extensa_89
T. extensa_90
T. spiraliflora_67
T. spiraliflora_68
T. mima var. chiletensis_20417
T. ecarinata_88
T. hildae_98
T. adpressiflora_50
T. marnier-lapostollei_21
T. mima_11
T. fuchsii_24
T. fuchsii_53
T. fuchsii var. stephanii_4
T. filifolia_49
T. filifolia_65
T. makoyana_80
T. makoyana_22
T. albida_269
T. complanata_90

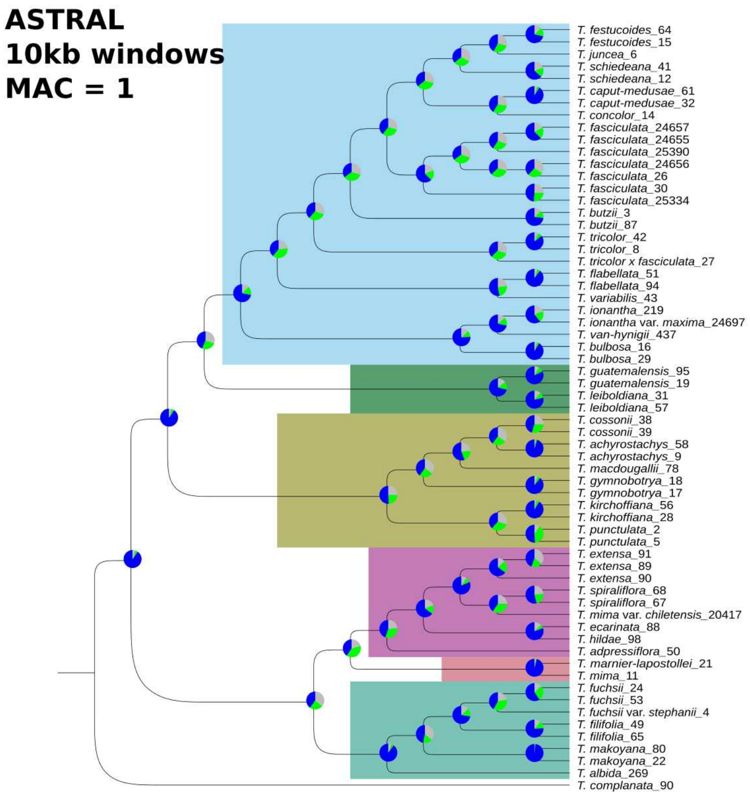
ASTRAL
10kb windows
MAC = 1
T. festucoides_64
T. festucoides_15
T. juncea_6
T. schiedeana_41
T. schiedeana_12
T. caput-medusae_61
T. caput-medusae_32
T. concolor_14
T. fasciculata_24657
T. fasciculata_24655
T. fasciculata_25390
T. fasciculata_24656
T. fasciculata_26
T. fasciculata_30
T. fasciculata_25334
T. butzii_3
T. butzii_87
T. tricolor_42
T. tricolor_8
T. tricolor x fasciculata_27
T. flabellata_51
T. flabellata_94
T. variabilis_43
T. ionantha_219
T. ionantha var. maxima_24697
T. van-hynigii_437
T. bulbosa_16
T. bulbosa_29
T. guatemalensis_95
T. guatemalensis_19
T. leiboldiana_31
T. leiboldiana_57
T. cossonii_38
T. cossonii_39
T. achyrostachys_58
T. achyrostachys_9
T. macdougallii_78
T. gymnobotrya_18
T. gymnobotrya_17
T. kirchoffiana_56
T. kirchoffiana_28
T. punctulata_2
T. punctulata_5
T. extensa_91
T. extensa_89
T. extensa_90
T. spiraliflora_68
T. spiraliflora_67
T. mima var. chiletensis_20417
T. ecarinata_88
T. hildae_98
T. adpressiflora_50
T. marnier-lapostollei_21
T. mima_11
T. fuchsii_24
T. fuchsii_53
T. fuchsii var. stephanii_4
T. filifolia_49
T. filifolia_65
T. makoyana_80
T. makoyana_22
T. albida_269
T. complanata_90

**ASTRAL**
**10kb windows**
**MAC = 2**

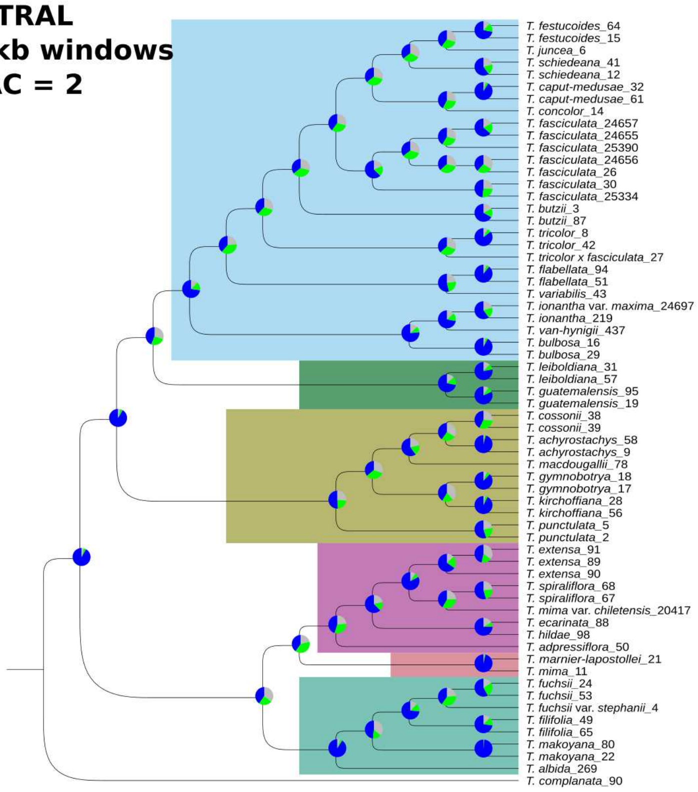


**ASTRAL**
**10kb windows**
**MAC = 3**

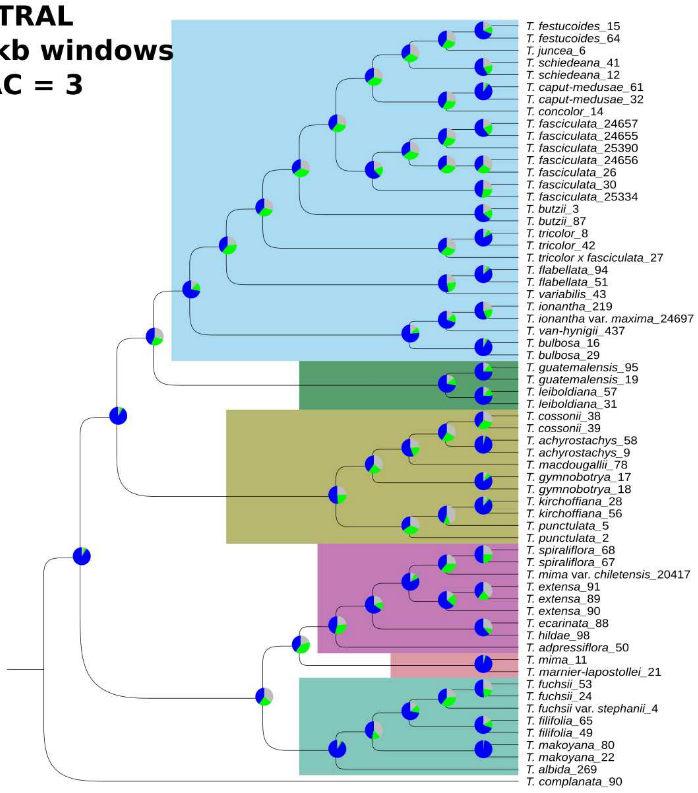

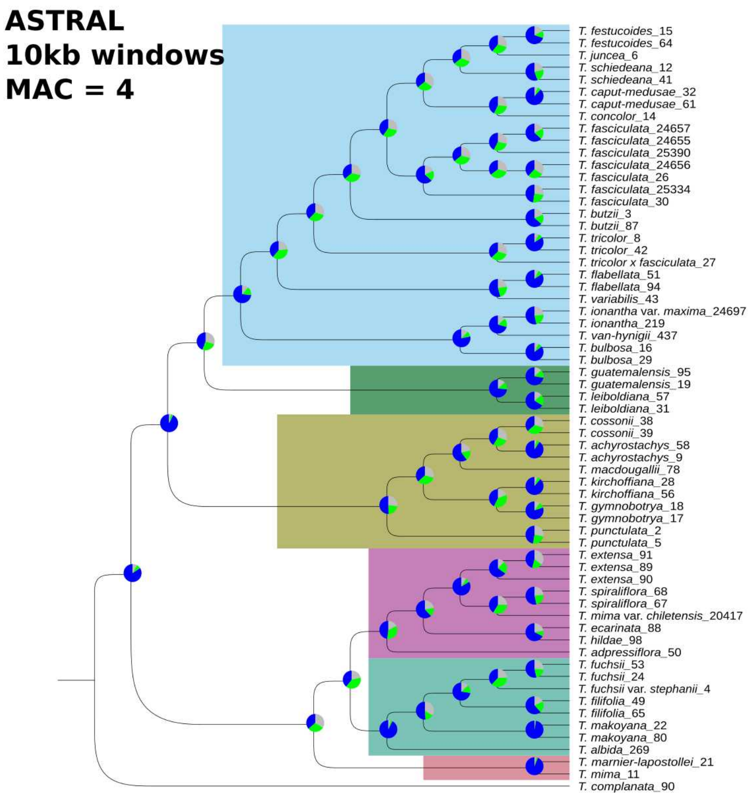
ASTRAL
10kb windows
MAC = 4
T. festucoides_15
T. festucoides_64
T. juncea_6
T. schiedeana_12
T. schiedeana_41
T. caput-medusae_32
T. caput-medusae_61
T. concolor_14
T. fasciculata_24657
T. fasciculata_24655
T. fasciculata_25390
T. fasciculata_24656
T. fasciculata_26
T. fasciculata_25334
T. fasciculata_30
T. butzii_3
T. butzii_87
T. tricolor_8
T. tricolor_42
T. tricolor x fasciculata_27
T. flabellata_51
T. flabellata_94
T. variabilis_43
T. ionantha var. maxima_24697
T. ionantha_219
T. van-hynigii_437
T. bulbosa_16
T. bulbosa_29
T. guatemalensis_95
T. guatemalensis_19
T. leiboldiana_57
T. leiboldiana_31
T. cossonii_38
T. cossonii_39
T. achyrostachys_58
T. achyrostachys_9
T. macdougallii_78
T. kirchoffiana_28
T. kirchoffiana_56
T. gymnobotrya_18
T. gymnobotrya_17
T. punctulata_2
T. punctulata_5
T. extensa_91
T. extensa_89
T. extensa_90
T. spiraliflora_68
T. spiraliflora_67
T. mima var. chiletensis_20417
T. ecarinata_88
T. hildae_98
T. adpressiflora_50
T. fuchsii_53
T. fuchsii_24
T. fuchsii var. stephanii_4
T. filifolia_49
T. filifolia_65
T. makoyana_22
T. makoyana_80
T. albida_269
T. marnier-lapostollei_21
T. mima_11
T. complanata_90

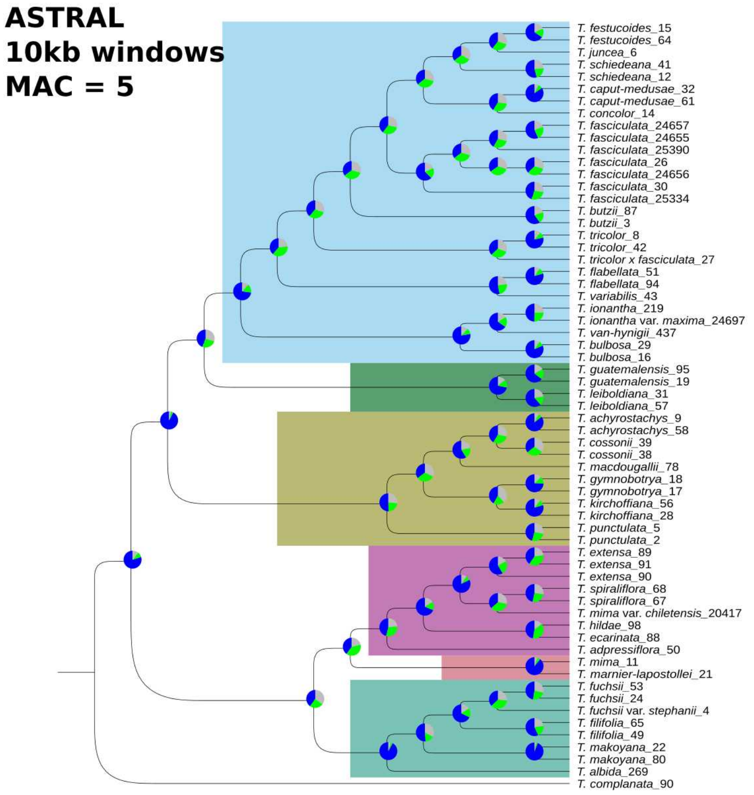
ASTRAL
10kb windows
MAC = 5
T. festucoides_15
T. festucoides_64
T. juncea_6
T. schiedeana_41
T. schiedeana_12
T. caput-medusae_32
T. caput-medusae_61
T. concolor_14
T. fasciculata_24657
T. fasciculata_24655
T. fasciculata_25390
T. fasciculata_26
T. fasciculata_24656
T. fasciculata_30
T. fasciculata_25334
T. butzii_87
T. butzii_3
T. tricolor_8
T. tricolor_42
T. tricolor x fasciculata_27
T. flabellata_51
T. flabellata_94
T. variabilis_43
T. ionantha_219
T. ionantha var. maxima_24697
T. van-hynigii_437
T. bulbosa_29
T. bulbosa_16
T. guatemalensis_95
T. guatemalensis_19
T. leiboldiana_31
T. leiboldiana_57
T. achyrostachys_9
T. achyrostachys_58
T. cossonii_39
T. cossonii_38
T. macdougallii_78
T. gymnobotrya_18
T. gymnobotrya_17
T. kirchoffiana_56
T. kirchoffiana_28
T. punctulata_5
T. punctulata_2
T. extensa_89
T. extensa_91
T. extensa_90
T. spiraliflora_68
T. spiraliflora_67
T. mima var. chiletensis_20417
T. hildae_98
T. ecarinata_88
T. adpressiflora_50
T. mima_11
T. marnier-lapostollei_21
T. fuchsii_53
T. fuchsii_24
T. fuchsii var. stephanii_4
T. filifolia_65
T. filifolia_49
T. makoyana_22
T. makoyana_80
T. albida_269
T. complanata_90

**Genome-wide D-statistic**

All D-statistics inference was performed as described in the main text. Briefly, we used the original implementation of the *D*-statistic in Dsuite v.0.5r45. We set no *a priori* knowledge of taxon relationships, but let Dsuite order each trio so that the BBAA pattern is more common to focus on topologies with minimal discordant patterns. We used the full VCF file with *T. complanata* as an outgroup.

**MAC=0**

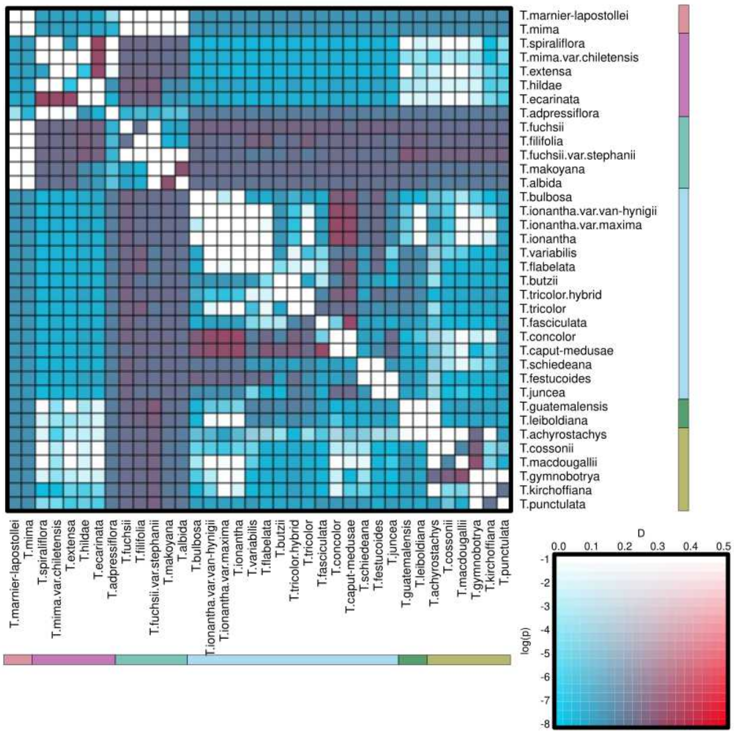


**MAC=1**

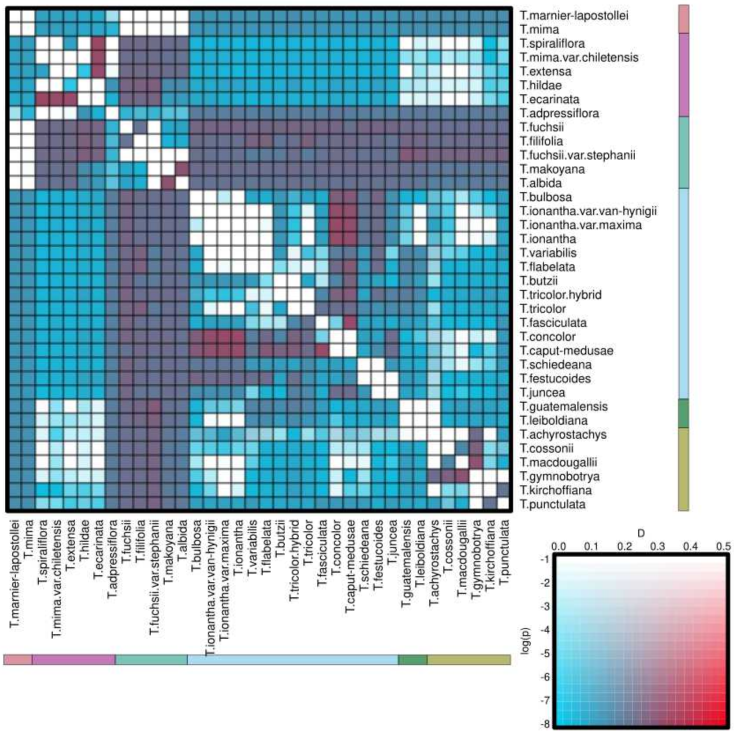

#### MAC=2

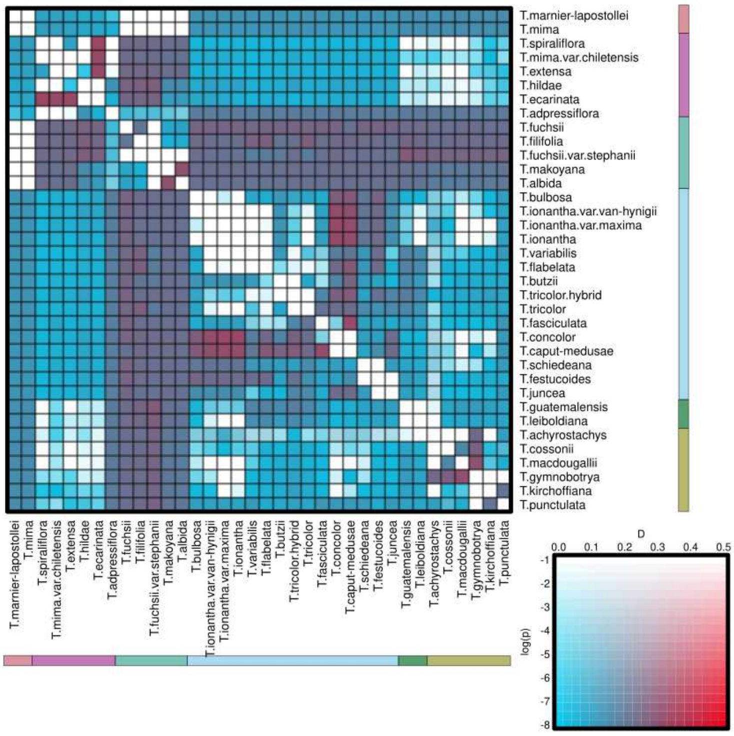
T.marnier-lapostollei
T.mima
T.spiraliflora
T.mima.var.chiletensis
T.extensa
T.hildae
T.ecarinata
T.adpressiflora
T.fuchsii
T.filifolia
T.fuchsii.var.stephanii
T.makoyana
T.albida
T.bulbosa
T.ionantha.var.van-hynigii
T.ionantha.var.maxima
T.ionantha
T.variabilis
T.flabelata
T.butzii
T.tricolor.hybrid
T.tricolor
T.fasciculata
T.concolor
T.caput-medusae
T.schiedeana
T.festucoides
T.juncea
T.guatemalensis
T.leiboldiana
T.achyrostachys
T.cossonii
T.macdougallii
T.gymnobotrya
T.kirchoffiana
T.punctulata
D
0.0
0.1
0.2
0.3
0.4
0.5
log(p)
-1
-2
-3
-4
-5
-6
-7
-8

#### MAC=3

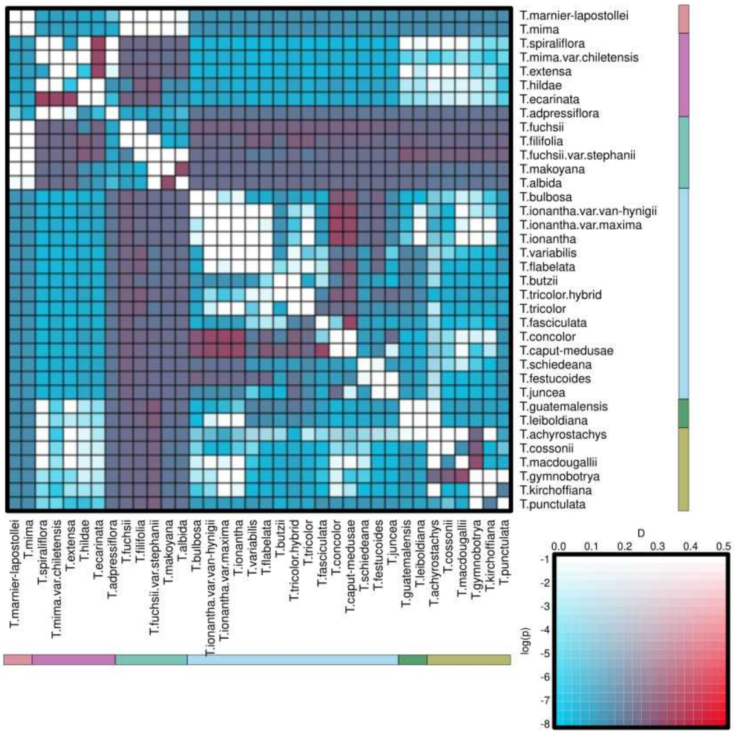
T.marnier-lapostollei
T.mima
T.spiraliflora
T.mima.var.chiletensis
T.extensa
T.hildae
T.ecarinata
T.adpressiflora
T.fuchsii
T.filifolia
T.fuchsii.var.stephanii
T.makoyana
T.albida
T.bulbosa
T.ionantha.var.van-hynigii
T.ionantha.var.maxima
T.ionantha
T.variabilis
T.flabelata
T.butzii
T.tricolor.hybrid
T.tricolor
T.fasciculata
T.concolor
T.caput-medusae
T.schiedeana
T.festucoides
T.juncea
T.guatemalensis
T.leiboldiana
T.achyrostachys
T.cossonii
T.macdougallii
T.gymnobotrya
T.kirchoffiana
T.punctulata
D
0.0
0.1
0.2
0.3
0.4
0.5
log(p)
-1
-2
-3
-4
-5
-6
-7
-8
